# Helixer–*de novo* Prediction of Primary Eukaryotic Gene Models Combining Deep Learning and a Hidden Markov Model

**DOI:** 10.1101/2023.02.06.527280

**Authors:** Felix Holst, Anthony Bolger, Christopher Günther, Janina Maß, Sebastian Triesch, Felicitas Kindel, Niklas Kiel, Nima Saadat, Oliver Ebenhöh, Björn Usadel, Rainer Schwacke, Marie Bolger, Andreas P.M. Weber, Alisandra K. Denton

**Author notes:** These authors contributed equally (Felix Holst, Anthony Bolger).

## Abstract

Gene structural annotation is a critical step in obtaining biological knowledge from genome sequences yet remains a major challenge in genomics projects. Current *de novo* Hidden Markov Models are limited in their capacity to model biological complexity; while current pipelines are resource-intensive and their results vary in quality with the available extrinsic data. Here, we build on our previous work in applying Deep Learning to gene calling to make a fully applicable, fast and user friendly tool for predicting primary gene models from DNA sequence alone. The quality is state-of-the-art, with predictions scoring closer by most measures to the references than to predictions from other *de novo* tools. Helixer’s predictions can be used as is or could be integrated in pipelines to boost quality further. Moreover, there is substantial potential for further improvements and advancements in gene calling with Deep Learning.

Helixer is open source and available at https://github.com/weberlab-hhu/Helixer

A web interface is available at https://www.plabipd.de/helixer_main.html

## 1 Introduction

To accelerate research progress in biology and bioengineering we need accurate *in silico* models to capitalize on rapidly increasing amounts of genomics data. Gene calling, or structural gene annotation, is critical to extracting biological knowledge from a genome, yet gene calling in eukaryotes lags far behind genome sequencing and assembly in quality, ease and speed [Michael and VanBuren, 2020, Yandell and Ence, 2012]. Improving gene calling will strengthen the foundation for all downstream applications, ranging from target-gene characterization, to transcriptomics and proteomics, genome-wide association studies and more.

Historically, eukaryotic gene calling was primarily performed by (generalized) Hidden Markov Models such as Genemark [Ter-Hovhannisyan et al., 2008], FGENESH [Solovyev et al., 2006] or AUGUSTUS [Stanke and Waack, 2003, Stanke et al., 2006]. These models lack the capacity to fully model biological complexity on their own, and therefore are today used as parts of data integration pipelines such as MAKER2 [Holt and Yandell, 2011], PASA [Haas et al., 2003] and BRAKER2 [Bru°na et al., 2021]. These pipelines require wetlab data and substantial computational resources, to the point they are often the rate-limiting step in a genome project. Inconsistent availability of such resources leads to a heterogenous quality of the resulting gene models. Errors are still being identified in gene models of extensively studied species such as human and mouse [Oz-Levi et al., 2019]. While in less-studied species, the overall lower annotation quality is disruptive for large-scale analyses, and can necessitate project-specific re-annotation [Wang et al., 2021]. Finally, many newly sequenced species entirely lack an annotation, with only circa 24% of latest Eukaryotic assemblies on NCBI having an accompanying annotation, far behind the 89% annotated in prokaryotes. Better tools are needed to easily produce consistent high quality annotations.

Deep Learning (DL) is a transformative technology where massive networks with the capacity to make extraordinarily complex and non-linear fits are trained on large amounts of data. DL has achieved astonishing performance in fields ranging from machine translation (e.g. GPT-3; Brown et al. [2020]), playing strategy games (e.g. AlphaGo; Silver et al. [2016]), deciphering mathematical equations [Lample and Charton, 2019, Fawzi et al., 2022] and to protein folding [Jumper et al., 2021]. In the last several years, DL networks have been employed for some elements of gene calling with promising results. DL has been used to predict genes in prokaryotes [Amin et al., 2018]; whether RNAs are coding or non-coding [Hill et al., 2018]; whether human sequences are exons or introns [Singh et al., 2021]; splice site type and strength [Louadi et al., 2019, Zeng and Li, 2022]; and to predict elements of a gene model, such as start codons, splice sites, and poly-adenylation sites [Wei et al., 2021, Liu et al., 2022]. That is, Deep Learning has already shown promising results in tasks related to – or that are a subset of – eukaryotic gene annotation.

In our previous work we developed Helixer: a proof of principle eukaryotic cross-species gene classifier that predicts the genic class of each base pair with very high quality [Stiehler et al., 2020]. While Helixer made substantial performance gains compared to an existing *de novo* predictor, and even exceeded the quality of *some* references in consistency with independent RNAseq data; Helixer’s base-wise predictions, as other applications of Deep Learning mentioned above, did not yet produce finalized gene models.

Here, we present Helixer as an application-ready tool, combining various improvements to our previous work and a a Hidden Markov Model-based post-processing tool named HelixerPost. Helixer takes the genome sequence as input (in fasta format) and outputs genome wide structural annotations for primary gene models (in gff3 format). Pre-trained models are now provided for fungal and invertebrate genomes, in addition to the previously available plant and vertebrate genomes. No extrinsic data nor species-specific retraining is required.

## 2 Implementation

Helixer takes DNA sequence as input, makes base-wise predictions for genic class and phase with pre-trained Deep Neural Networks, and processes these predictions with a Hidden Markov Model to into primary gene models (Fig.S1).

### 2.1 Retained Helixer architecture

Unless otherwise specified, the setup, processing and architecture was retained from our previous work [Stiehler et al., 2020]. In brief, Helixer implements a Deep Neural Network that takes substrings of the genomic DNA sequence as input, and outputs genic class for each base pair. The input is encoded as a one-hot vector of C, A, T, G. The output was a one-hot encoding of Intergenic, UTR, CDS, Intron for each base pair, which is still included.

### 2.2 Straight forward modifications

#### Layer type

We move from the stacked bidirectional Long Short-Term Memory (bLSTM) architecture used before, to a hybrid architecture comprised of initial convolutional layers followed by bLSTM layers.

#### Validation set

Validation after each training epoch is now performed on a cross-species ‘validation set’, i.e. every species not currently assigned to training nor reserved for testing, which provides closer to predictive-target feedback than the previous within-genome split.

#### Applicable range

In addition to vertebrates and land plants, models were trained and evaluated to predict on invertebrate and fungal data.

#### Performance

Performance improvements, in particular regarding data-import, allowed for an ∼5 x improvement in speed, which in turn allowed for usage of more training genomes.

### 2.3 Species selection

While developing Helixer, we found that the generalization performance of a trained model is highly dependent on both the quality and the quantity of the training genomes. Finding the optimal trade-off proved to be difficult and often a bit counter-intuitive.

To automatically obtain more generalizable, higher quality, and consistent models, the process of splitting training and validation genomes was subjected to a custom hyperparameter optimization, which essentially selected promising random sets of training genomes via 2-fold cross-validation, and then remixed the best genomes from the folds for a final evaluation.

First, the available training and validation genomes were divided randomly into two folds. Second, within each fold, a number (Tab.S14) of models were trained each on randomized divisions of the fold into training and validation genomes. Third, the performance (genic F1) of each model was evaluated on all the genomes in the other fold. Fourth, the best two divisions of training and validation in each fold were recombined. So given the best two splits in fold A as A1 and A2, and the best two splits in fold B as B1 and B2, models were then trained on A1 + B1, A1 + B2, A2 + B1, A2 + B2. All remaining non-test species were used for validation. Fifth, final evaluation was performed on the intersect of all validation sets from step four, and this was then used for the final ranking of the models. The top models can be used for ensembling, but here the best model was used alone unless otherwise specified.

This was implemented here: https://github.com/alisandra/SpeciesSelector

As necessary, a last round of human expertise and hyperparameter optimization was employed to finalize the selection.

### 2.4 Improved reflection of biological importance of predictive task via the loss function

#### Transition weights

The base-wise probabilities output by the original version of Helixer [Stiehler et al., 2020] were scored against the references with the categorical cross-entropy loss function, and the penalty for each base pair varied at most by the setting of class weights (to compensate for class imbalance). This setup does not fully reflect the biological significance of different mistakes made by the network; and modifications to the Helixer architecture presented here address this.

The previous setup weighted all non-masked base pairs in an annotation class equally, and the network could achieve a low loss by predicting the broad regions correctly and confidently, while accumulating only a small penalty for comparatively few uncertain predictions around class transitions. However, biologically speaking, four of the class transitions (start codon, stop codon, donor splice site and acceptor splice site) can cause a non-sense prediction if they are off by even a single base pair by introducing a frame shift in the final protein sequence. To better reflect the biological importance of these transitions in the loss function, we implemented transition weighting; which up-weighted the loss directly before and after a transition. The transcript start and end were not up-weighted as preliminary analyses (not shown) indicated this was not beneficial, potentially due to the extreme noise in the transcript start and end labels be it from errors, biological variation, or both.

In more detail: For all transitions transcription start site, start codon, donor splice site, poly-adenylation site, stop codon, acceptor splice site the indices of last base pair of the proceeding class, and the first base pair of the next class were identified. Where *any* transition occurred in a pool, the whole pool was up-weighted (generally 12 × for start & stop codons, and 3× for splice sites). Where two transitions occurred in the same pool (e.g. start codon and donor splice site), weights were summed. Non-transition pools received a weight of 1. The resulting weights were used to multiply the sample weights (which are additionally adjusted according to error-masks as previously described.)

#### Phase prediction

To further associate the loss function with the resulting protein sequence, we added a second matrix ‘phases’ as an auxiliary prediction. This was also beneficial for post-processing (see below). The target phases matrix is a one-hot vector for the classes None, phase 0, phase 1, phase 2. When the main target class is CDS, this is set to the phase (number of base pairs until the start of the next codon); in all other regions it is set to None. The overall loss is the weighted sum of the primary loss (weight=0.8), and the categorical cross entropy loss of the phase predictions (weight=0.2).

### 2.5 Post-processing into final primary gene models

We implemented a post-processing tool, HelixerPost: https://github.com/TonyBolger/HelixerPost, that takes the base-wise predictions of genic class and phase as input and outputs a final gff3 file of primary gene models.

HelixerPost makes a double pass of the genome. In the first pass, genic regions are identified as sections where the average non-intergenic prediction of a sliding window (recommended 100bp) stays above a given threshold (recommended 0.1) and peaks at or above a higher threshold (recommended 0.8).

In the second pass, the primary gene model(s) most consistent with the base-wise predictions in each genic region are determined using a Markov Model which encodes biologically plausible states and transitions. The major states and transitions of the Markov Model are similar to those standardly used in gene calling, with various states representing intergenic, 5’ UTR, CDS, and 3’ UTR sequence. Specific CDS states are used to represent start and stop codons. Coding phase is also encoded within the states for start, regular and stop codons, thus requiring 3 start codon states, 3 regular codon states and, perhaps surprisingly, 4 stop codon states, with the additional stop state needed to distinguish between TAx and TGx partial stop codons. Intron state is encoded using an additional set of substates, for both UTR states and 8 of the 10 CDS states. The Start2 and Stop2 states do not need additional intron states since introns starting immediately after start and stop codons are functionally equivalent to introns starting immediately after regular codons, and anywhere in the 3’ UTR respectively. Three different forms of intron splicing are represented, GT-AG, GC-AG, both of which are mediated by U2, and AC-AT as mediated by U12. Two states are used for each intron form, representing the start and continuation of each intron form, thus requiring 6 additional states for each primary state. Thus, in total, the HMM uses 73 states, with 1 intergenic, 2 UTR states, 10 CDS states, and 6 additional intron substates each for 10 of the 12 UTR/CDS states.

Minimal intron lengths are implemented by pausing the HMM state for additional bases when an intron start state is detected (49 additional bases for U2 mediated introns and 29 for U12 mediated introns). In all other cases, comsumption of each base is associated with a HMM state transition, according to set of possible valid state transitions.

The optimal scoring path through this Markov Model for a given underlying sequence and set of base-wise predictions is determined with the Viterbi algorithm [Viterbi, 1967], a dynamic programming algorithm. The scoring system penalizes discrepancies where the state of the Markov Model differs from the base-wise predictions produced by Helixer based on their confidence. In addition, HMM states which require specific sequence contexts, such as start and stop codons, and intron donor/acceptor sites, are strongly penalized in the absence of those sequences, and thus unlikely to be chosen as the optimum.

For performance reasons, all calculations are performed in negative log2 space, allowing multiplicative probability to be maximized by minimizing the total penalty accumulated.

### 2.6 Iterative Release

Note that as models were released and employed by the community as progress was made during development, all such transitional “best models” have been included for the evaluation; regardless of whether they differ from the final models on hyperparameters or the training species set. The exact parameters (Tab.S9, S10, S11, S12) and training species (Tab. S2.2, S6, S7, S8) are listed for transitional and current best models.

### 2.7 Nomenclature

Released models have been assigned an ID, as follows <lineage>_<release>_<key>_<rank>. Where the lineage indicates the appropriate application lineage of the trained model, the release indicates the code version with which it is compatible. The key is a brief marker of ‘a’ for automatic species selection and ‘m’ for manual, which can be either entirely manual or automatic followed by manual tuning. Finally, rank indicates relative model performance within the affore mentioned categories (lower is better).

Tag v0.3.0 was used or is functionally equivalent to the version used troughout, unless otherwise specified.

### 2.8 Usability

As Helixer and its sub-components are now directly practically applicable, we also made major improvements to the usability to facilitate said application. In particular, all three components can now be conveniently installed together via Docker or Singularity. Further, all components can be used together for inference with a convenience wrapper, ‘Helixer.py‘, so that the user inputs a genomic fasta file, and receives a final gff3 (standard annotation format) file as output. Instructions and examples are included.

## 3 Methods

### 3.1 Data

The training and evaluation data was acquired from RefSeq and Phytozome13 and split into the combined training and validation set separately for each phylogenetic group as the project progressed. Details and species splits are listed in section S2.1.

### 3.2 Parameters

Non-default and result-affecting parameters for all tools listed below are listed in Tab.S13.

### 3.3 Annotation quality comparison

Two established HMM tools were used to set baseline expectations for a *de novo* gene calling tool. AUGUSTUS (3.3.2) [Stanke et al., 2006] was used as a high performance baseline where trained models were available (Tab. S15), but for feasibility reasons, no re-training was performed here. Therefore, for the complete test set, GenemarkES (4.71_lic) [Ter-Hovhannisyan et al., 2008], which uses unsupervised training was additionally employed. Nevertheless data is missing for GenemarkES for the 10.7Gbp plant *Triticum dicoccoides*, as GenemarkES did not finish given 24 threads within 1 week.

For comparing Helixer’s prediction to these tools and the reference, primary transcripts were used unless otherwise specified. Primary transcripts are defined as the splice variant producing the longest protein.

### 3.4 Helixer Test Inference

Inference for test species was performed with an ensemble (element-wise average of raw predictions) of the best two models for fungi (fungi_v0.3_a_0100 and fungi_v0.3_a_0200), and due to time constraints for developing and evaluating the models with only the single best model (default) for the other three groups. Subsequence length and other parameters were set to easily contain typical gene lengths for the phylogenetic group (Tab. S16).

### 3.5 Evaluation

#### F1 scores

Reported F1 scores are the combined base-wise F1 for CDS, UTR, and intron (Genic) or CDS and intron (Subgenic), with details described in Stiehler et al. [2020]. For the broad evaluation including training and validation genomes as well as for the validation genomes during training, the F1 scores were calculated on a random sample of 800 of the subsequences from each genome. For test genomes, F1 scores are calculated genome-wide.

#### Homology comparisons

Proteome completeness was estimated with BUSCO (5.2.2) [Simão et al., 2015]. For the plant test genomes, completeness and annotatability were further estimated by comparison to curated gene families by assigning Mapman4 protein categories [Schwacke et al., 2019] with Mercator4 (5.0) [Bolger et al., 2021]. More generally, comparability between proteomes was further assessed by clustering into orthougroups with OrthoFinder (2.5.4) [Emms and Kelly, 2019]. For the sake of comparison, clustering was performed on 12 plant test genomes (those that finished in 1-week for GenemarkES); and consistency was further double-checked for all 13 species with only the references and Helixer’s predictions.

### 3.6 *Arabidopsis thaliana*-annotation comparison

Pairwise comparison of the structural annotations from Helixer to TAIR10 and Araport11 for the model species *A. thaliana*.

As Helixer predicts only a single splice variant per locus, the number of gene loci was used for counting purposes. First, identical protein sequences were identified from each reference annotation versus Helixer. If any splice variant from the comparison reference genome was present in the Helixer set, it was labelled as ‘identical’. The remaining set of sequences were tested using BLAST (E-value 10^*−*8^, Altschul et al. [1997]) to find homologous sequences between the reference and Helixer. This identified similar protein-coding sequences between each pairwise comparison. Finally, the remaining sequences were compared to sequences from 420 genome projects representing all streptophytes clades, to identify orthologous sequences. Orthologous candidates were verified by triangles of Reciprocal Best Hits [Tatusov et al., 1997].

For the figure, the same RNAseq data and processing was used as in [Stiehler et al., 2020].

### 3.7 Ablations

To assess the actual effect of various optimizations included during development the best plant model was compared to plant models trained without phase, with lower transition weights or with an LSTM instead of Hybrid model. Exactly how parameters differed from land_plant_v0.3_a_0080 is listed (Tab. S17), all other training parameters were held constant. Ablations were evaluated on all plant test genomes except the very large *T. dicoccoides*. Inference parameters differing from other analyses can be found here (Tab. S18).

### 3.8 Benchmarking

Benchmarking was performed on the following machines. A) Workstation with an AMD Ryzen 9 5950X CPU, GeForce GTX 1080 Ti graphics card, 32GiB 2133MHz DDR4 RAM, and a Samsung 980 PRO 2TB NVMe SSD. B) Workstation with Intel(R) Xeon(R) CPU E5-2623 v4 @ 2.60GHz, GeForce GTX 1080 Ti graphics card, 64GiB 2400MHz DDR4 ECC RAM, and an INTEL SSDSC2BB24 SSD. C) for comparison between Helixer and the other tools on a workstation with Intel(R) Xeon(R) W-2125 CPU @ 4.00GHz, 32GB (4x 8GB) DDR4 PC2666 ECC RAM, GeForce GTX 1080 graphics card, and an Intel D3-S4510 240GB SSD. All benchmarks were performed single-threaded, with performance-affecting parameters as specified here (Tab. S19).

## 4 Results

### 4.1 The released Helixer models show state-of-the-art performance compared to existing *de novo* gene calling tools and to previous Helixer models

As our previous work [Stiehler et al., 2020] set the state-of-the-art for base-wise predictions, we first compared the latest models to models trained with the hand selected 6 (vertebrate) or 9 (plant) species used before, as well as to all intermittently released models (Fig. 1, S2, S3, S4, S5, S6, S7, S8). The best models released here (vertebrate_v0.3_m_0080, invertebrate_v0.3_m_0100, land_plant_v0.3_a_0080, and fungi_v0.3_a_0100), had the highest median Genic F1 for their phylogenetic target range, and showed more balanced performance across said range, compared to other models. Nevertheless, no model was consistently better for *all* species, so all plotted models are released in order to allow researchers to select the model likely to perform best for their species of interest.

**Figure 1:**
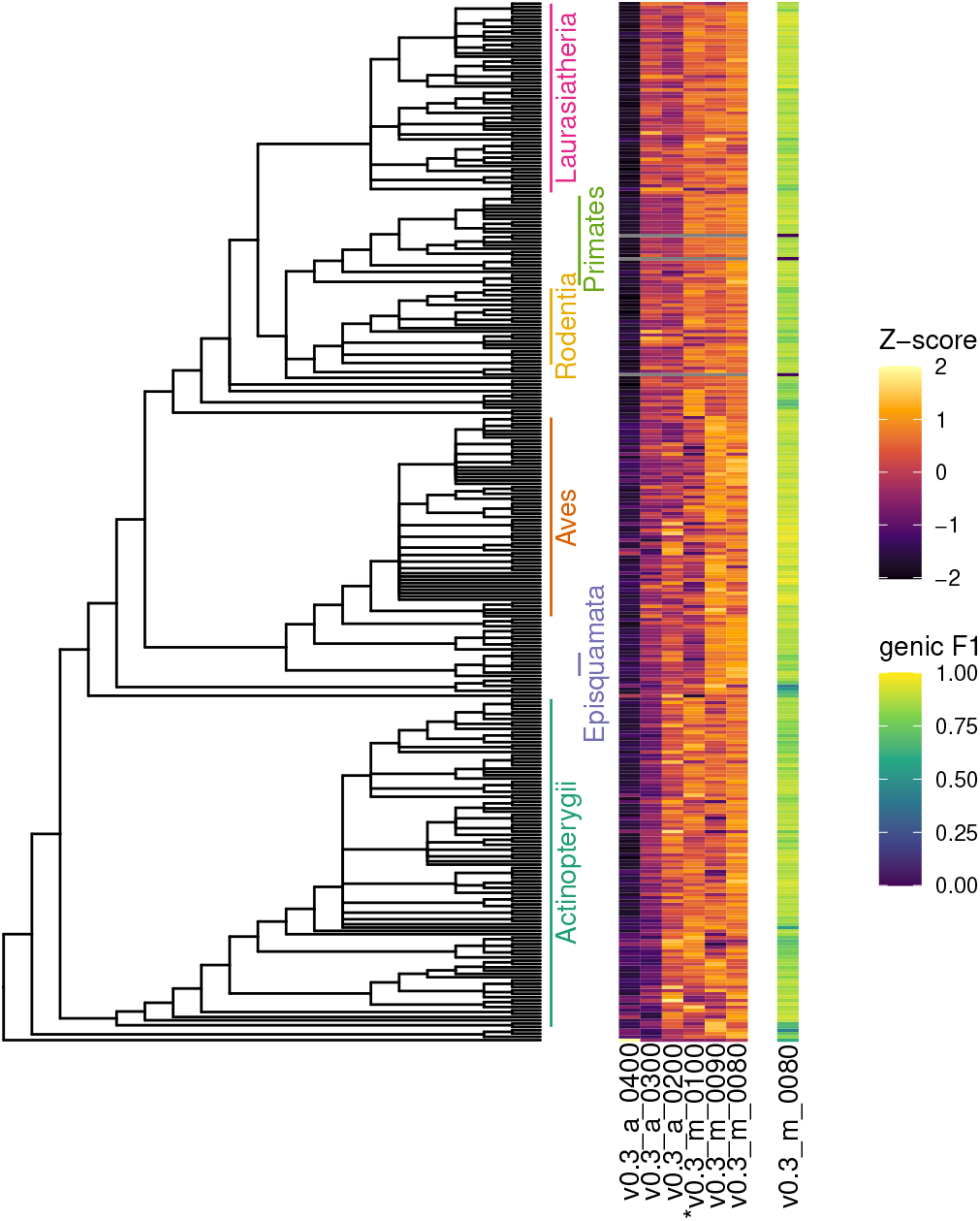
Performance of candidate and best vertebrate models across vertebrates. The absolute performance is measured with genic F1 on a random selection of 800 subsequences of each species and displayed for the overal best v0.3_m_0080 model (right). For peceptibility, differences between the genic F1 of models are displayed as a Z-score (middle). Models marked with * used the same training species as in Stiehler et al. [2020]. Grey indicates data is not available.

While promising, high base-wise performance of raw Helixer predictions does not necessarily imply that the performance is maintained through post processing. Therefore we further computed the Subgenic F1 for the final gene models output by HelixerPost on the selected test species (Fig. 2a, S9, S11, S13, S15, Tab. S20). As in our previous work, the Subgenic F1 was chosen as evaluation metric - instead of the more comprehensive Genic F1 - for comparability with exiting HMM tools that don’t (necessarily) predict UTRs.

**Figure 2:**
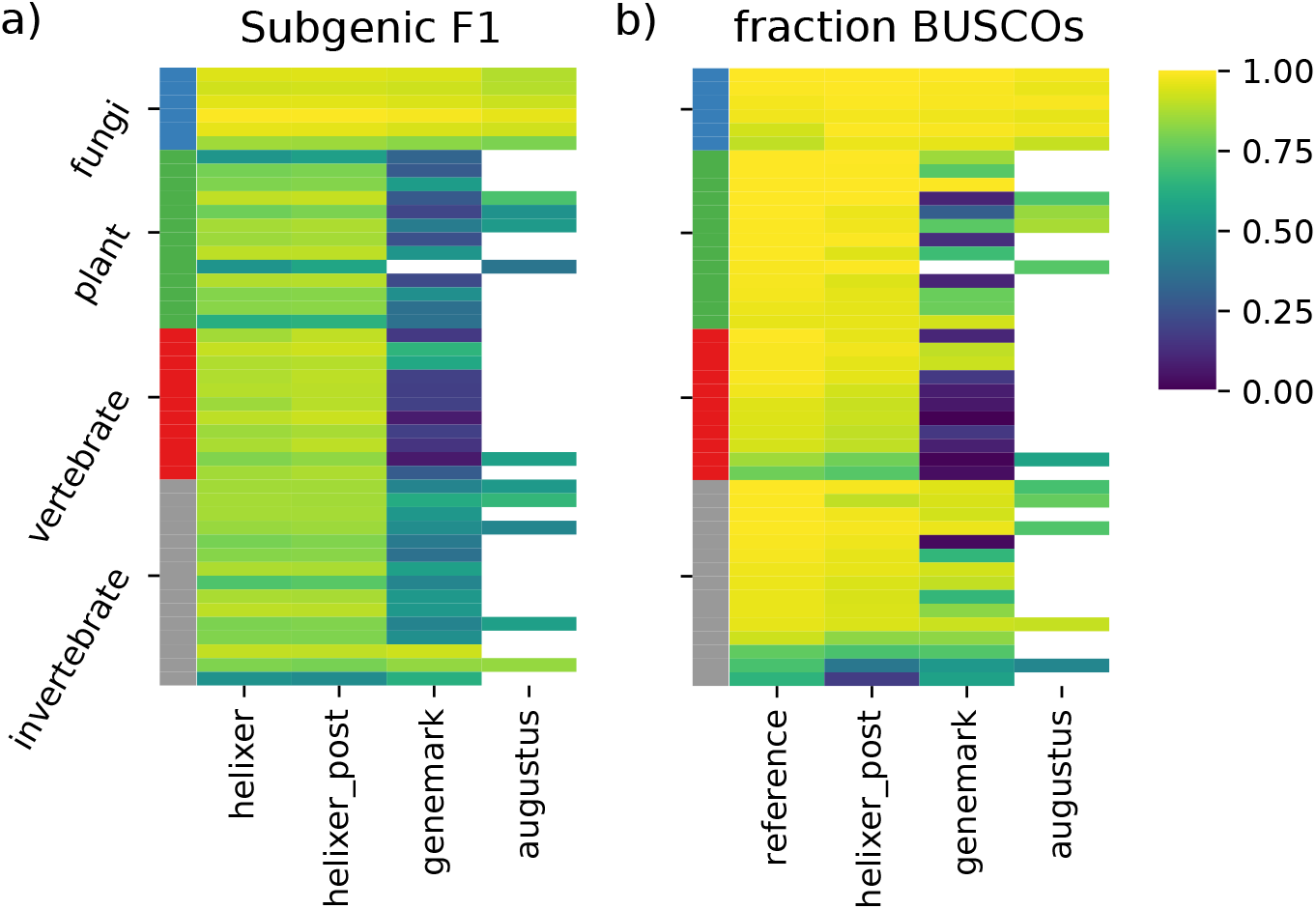
Test set annotation quality measured as (a) Subgenic F1 compared to the reference; and (b) as fraction of complete BUSCOs found in final annotation. The labels ‘helixer’ and ‘helixer_post’ indicate raw predictions from the neural network, and those post-processed by HelixerPost, respectively. White indicates data is not available.

Importantly, we see that predictions post-processed via HelixerPost have very similar performance to the raw Helixer predictions, with even a small increase in 33 of 45 test species. Additionally, we inferred gene models with two existing HMM tools, GenemarkES, and – where trained models were available for the test species or a closely related species – AUGUSTUS. The Subgenic F1 of HelixerPost was higher than Genemark in 42 of 45 species, and than AUGUSTUS in 14 of 15 species, with an average margin of 0.38 and 0.17 respectively.

To move from base-wise to protein level evaluation, the completeness of predicted proteomes [Simão et al., 2015] was quantified for the reference and all three predictive tools (Fig. 2b, S10, S12, S14, S16, Tab. S21). Here, the reference consistently had the highest performance (39 of 45 species); followed closely by HelixerPost, then AUGUSTUS, and GenemarkES with average reductions in BUSCOs found of (3.7, 12.8, and 33.9% respectively). The HelixerPost annotation contained more BUSCOs than GenemarkES and AUGUSTUS in 41 of 45, and 14 of 15 species, respectively. Interestingly, the three invertebrates where HelixerPost scored behind GenemarkES in Subgenic F1 and BUSCO count, had the overall lowest BUSCO count in the reference annotations. This hints at larger challenges related to annotating these genomes, be it exceptional divergence or simply a paucity of well-annotated genomes in close phylogenetic proximity to use either for training or for the homology mapping step of an annotation pipeline. However, it may also simply reflect that the invertebrate models are the newest and have seen less optimization.

### 4.2 Ablation analyses

As many changes were made during development, we performed an ablation analysis by individually excluding each of the major changes, training a new model, then predicting and comparing prediction quality to the final model (Fig. 3).

**Figure 3:**
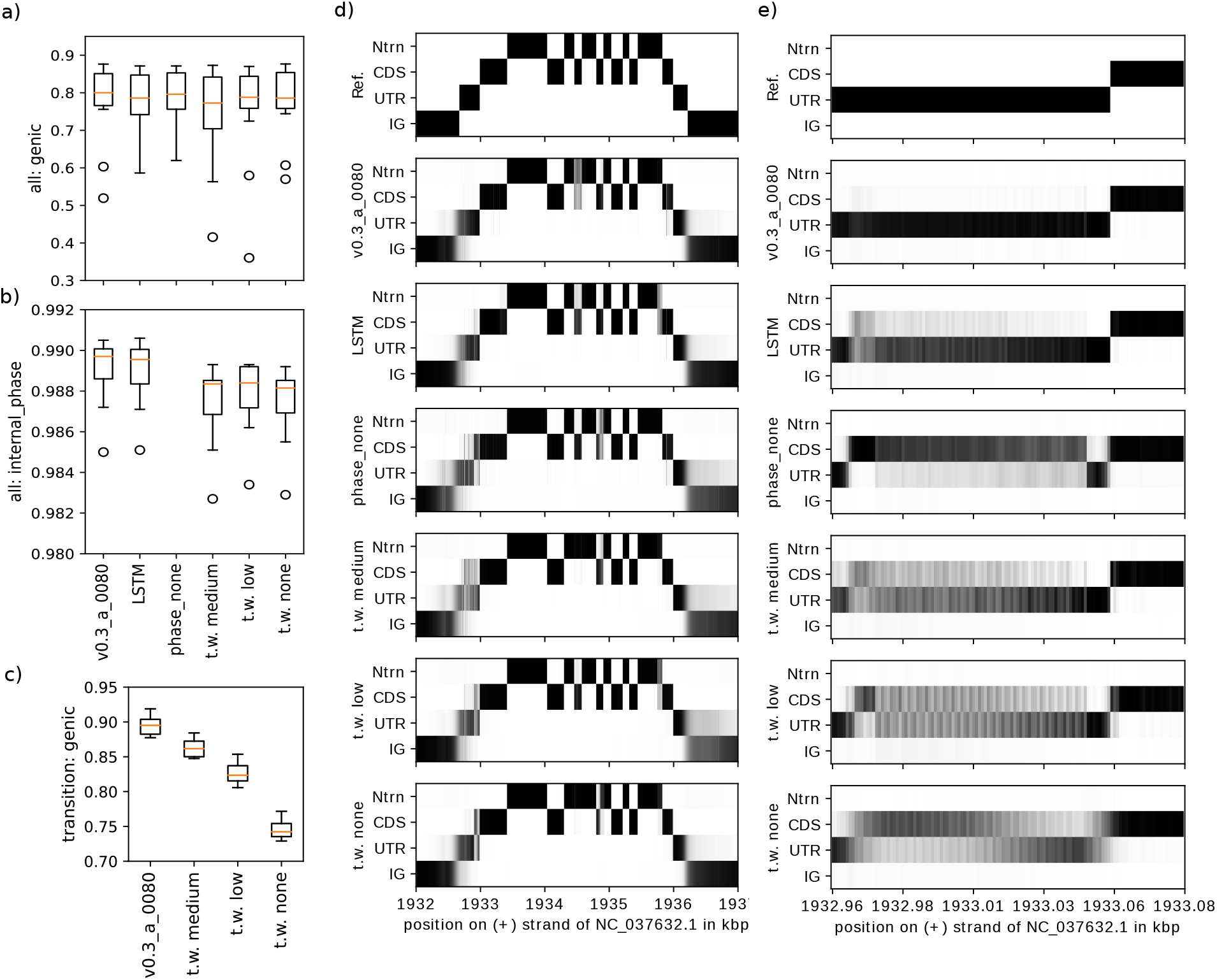
Performance of models with or without various optmizations to the models (phase, transition weights, Hybrid vs LSTM) on test plant species. v0.3_a_0080 is the final model and contains all optimizations; LSTM has phase and the final transition weighting, but uses the pure LSTM architecture used in Stiehler et. al 2020; the phase_none model was trained without the phase matrix, the t.w. medium, low, and none models use progressively lower transition weights compared to v0.3_a_0080 (see methods). a) genic F1 for all postions, b) accuracy of internal phase predictions, i.e. accuracy as fraction of phase prediction when both the reference and the predictions had a non-none phase. This captures only frameshift errors. c) effect of transition weights on the genic F1 score *only* at the basepairs before or after a transition. d) example prediction of the genic class matrix compared to the reference 1-hot encoding, white=0, black=1 with grays being intermediate values. e) a zoom in of specifically the same gene, specifically around the start codon.

While none of the changes had a major effect on the overall Genic F1 score (Fig. 3a), this was not the target of most of the changes. Without transition weights (Fig. 3e), the network displays high uncertainty immediately around start and stop codons, as well as acceptor and donor splice sites. All of the low, medium, and final transition weights push the networks towards sharper transitions in an example gene, both right at the start codon (Fig. 3e), and more widely helping to clear up uncertainty throughout the UTR (Fig. 3d), with the final transition weights being most effective. To quantify this on a larger scale, we calculated the Genic F1 specifically for the base pairs immediately before or after transitions. We see that progressively higher transition weights resulted in the expected increase in transition Genic F1, but with the largest difference being between low transition weights, and none (Fig. 3c). Moreover, the models with reduced transition weights show markedly more internal phase mistakes, i.e. where both reference and predictions have a 0-2 codon phase, but the phase do not match (Fig. 3b).

Interestingly, including the prediction of the phase matrix and loss also appears to have improved the accuracy around transitions, at least in this example (Fig. 3d, e). But the most important difference for the phase_none model is simply that without phase, the predictions aren’t suitable for post processing into final gene models.

The LSTM model and the Hybrid model have similar performance.

We acknowledge that this analysis could be made more conclusive by addressing the effect of random starting parameters by adding replicates, or where applicable (the number and arrangement of trainable parameters is only constant for the transition weight ablations), recording and reusing a random seed.

### 4.3 Helixer’s annotations approach reference quality

The F1 metrics shown so far treat the references as the ‘ground truth’; however, the provided references themselves are ultimately the output of a gene annotation pipeline, i.e. data-supported predictions. Measuring performance against such references is particularly limited for understanding how good the absolute performance is as it approaches that of the provided reference. In the BUSCO analysis above we already observed that Helixer’s performance was close to that of the reference. Therefore, we used two homology-based methods to evaluate in more detail the quality of the references and predictions alike (Fig. 4, S17, S18, S19, S20, S21, Tab. S22) for the plant test species.

**Figure 4:**
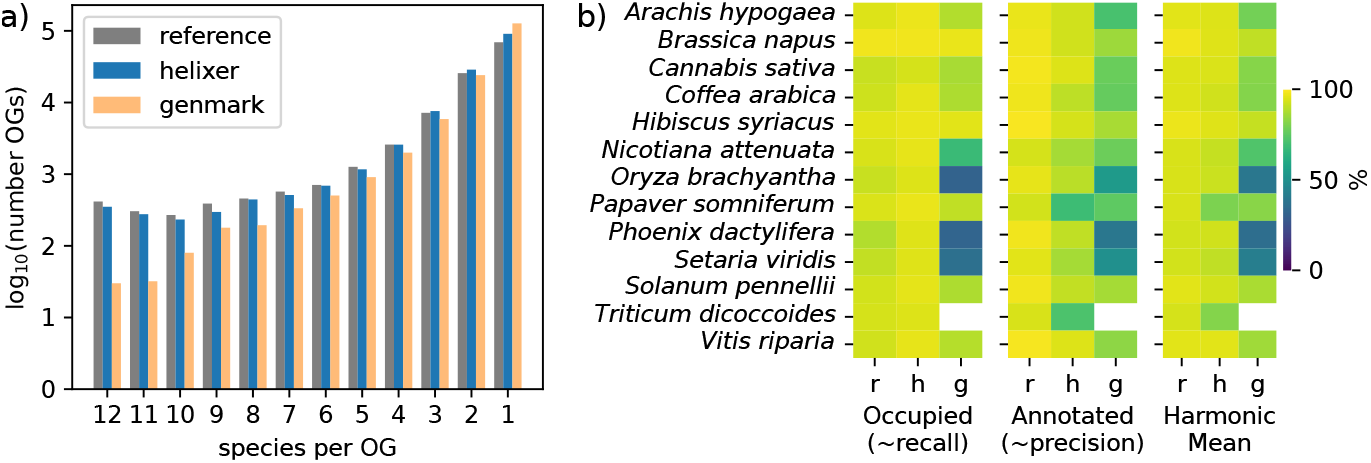
Homology based evaluation of annotations. a) Orthogroup occupancy (number of species represented out of 12) for orthogroups based on the reference, Helixer’s, and Genemark’s annotations (each clustered individually). b) Comparison of proteomes to currated Mapman4 protein annotations. r=reference, h=helixer (post processed), g=genemark. The harmonic mean is that of the %Annotated and the %Occupied, and here is conceptually a proxy for the F1.

First, we use the ability of a set of proteomes to form orthogroups as a reference-free indicator for quality, which is particularly relevant when considering annotation applicability for comparative genomic tasks.

The more accurate annotation is expected to have more orthogroups with all 12 species represented and generally more orthogroups with higher numbers of species represented. Here, the reference performs best with 0.38% of orthogroups containing all 12 species, and 36.5% containing two or more species; followed by Helixer at 0.26% and 31.1% for the same statistics, and Genemark at 0.019% and 21.2%. The reference has the most orthogroups with 4 or more species, Helixer, with 2 or 3 species, and Genemark with only 1 species (Fig. 4a). Notably, Helixer’s performance by these metrics was more similar to that of the reference, i.e. to extrinsic data-supported predictions, than to Genemarks predictions, which, comparable to Helixer’s are made from DNA sequence alone. Moreover, all of the reference annotations were respectable (minimum complete BUSCOs = 96.5%, and 8 of the species above 99%).

Second, given that the orthogroup-based numbers could potentially be skewed by consistent cross-species mistakes, such as mis-annotation of transposons, we used the Mapman4 protein annotations (curated plant protein coding gene families) to evaluate annotation quality. This further gives us a proxy for precision (the % of proteins that could be annotated) and recall (the % of gene families occupied by a protein). Measured this way, the reference had an average precision, recall, and harmonic mean (of precision and recall) of (0.966, 0.931, 0.948), Helixer (0.878, 0.958, 0.914) and Genemark (0.719, 0.742, 0.724); respectively (Fig. 4b). Thus the *de novo* annotations from Helixer, show higher recall than the references for the plant species test set; but are behind the references on precision, particularly for the specific species *Papaver somniferum* and *Triticum dicoccoides* (Fig. 4b, S17, S19, S21, Tab. S22); which we note have the most fragmented assemblies leading to an idea on how to further improve Helixer’s predictions (supplemental section S3).

### 4.4 Applicability for filling gaps in even good references

We compared the annotations produced by Helixer for the model plant *A. thaliana*, to the established high quality reference annotations: TAIR10 and Araport11. When comparing Helixer to Araport11 (Fig. 5a), the majority (∼70%) of gene loci were found to have an exact match in Helixer. From the remaining ∼30%, around 27% are highly similar but varied in their length. The unmatched sequences which remained from Helixer and Araport11 were considered as true annotations if the sequences were found to be conserved in at least 20 streptophytes species. For the Araport11 annotations, 93 of 1401 were found to be true annotations while 102 of 711 Helixer annotations were identified as true. Similar results were found when comparing Helixer with TAIR10 annotations (Fig. S22, S23).

**Figure 5:**
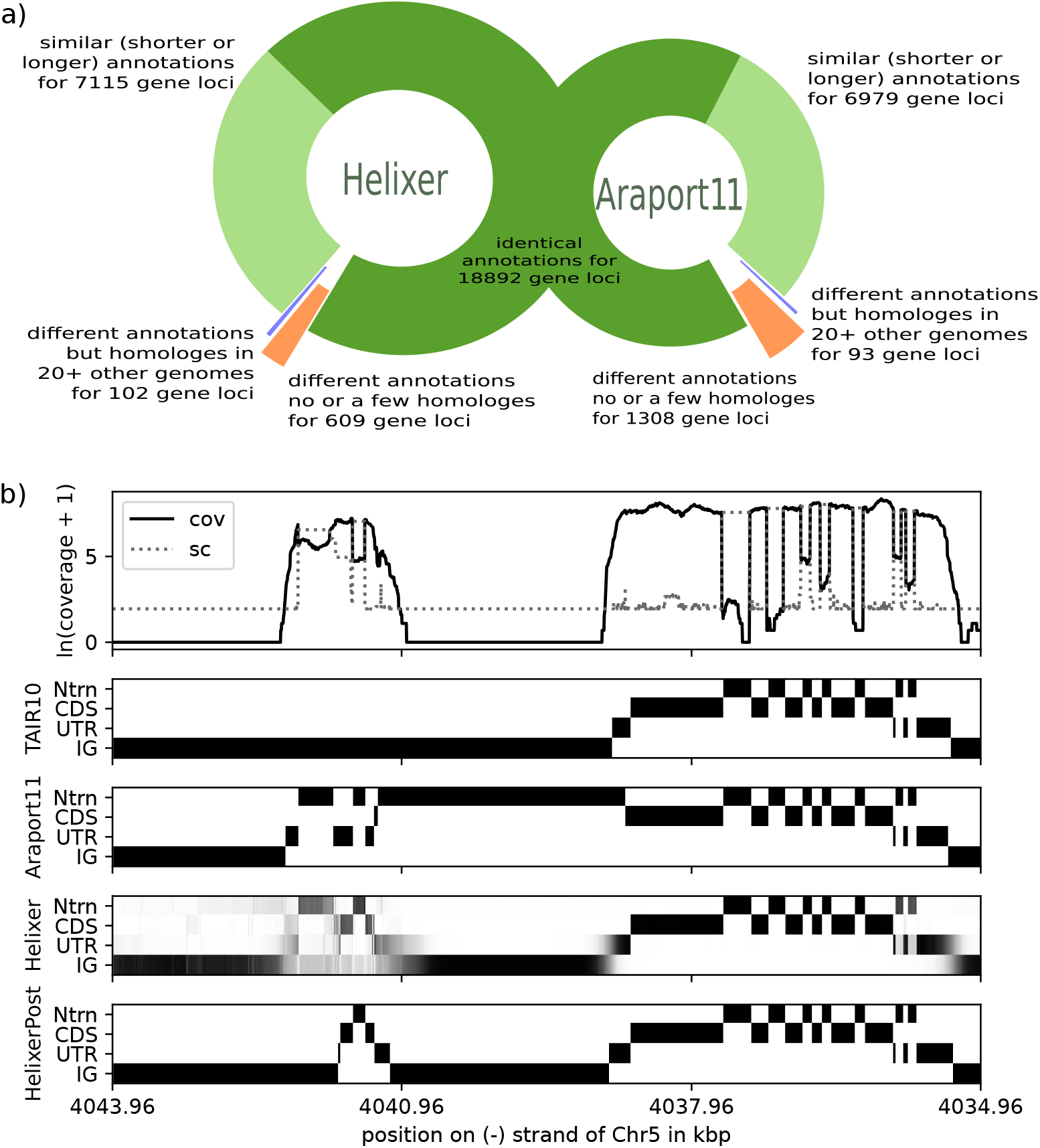
Comparison of the Arabidopsis thaliana proteome predicted by Helixer to the existing Araport11 annotation. a) breakdown of the overlaps and differences between predicted proteomes. b) visualization of expression and available annotations at the genomic loci of the Phosphatidylinositol N-acetylglucosaminyltransferase *γ* subunit (Chr5, - strand, from 4041091 to 4041627, via HelixerPost).

From these 102 Helixer annotations not found in the Araport11 reference annotation, a notable example is the Phosphatidylinositol N-acetylglucosaminyltransferase *γ* subunit. This complex is known to be active in *A. thaliana* [Lalanne et al., 2004, Beihammer et al., 2020] *and the γ* subunit locus identified by Helixer shows expression (Fig. 5b, S24). However, this *γ* subunit is entirely missing from TAIR10, and has a chimeric annotation with the next gene in Araport11, resulting in a mere 4bp of out-of-frame overlap between the resulting protein annotations.

While the final post-processed annotation from Helixer has truncations in at least UTR and introns relative to both the RNAseq data and the raw predictions, the resulting protein was long enough for homology-based identification.

This highlights the power of Helixer to complement and improve even the most polished reference annotations.

### 4.5 Benchmarking

The computational load for Helixer comprises three main parts, namely conversion of input sequence into numerical form, prediction with the neutral network, and the HMM for post-processing of the predictions. The first two are expected to scale linearly with genomic sequence length with the qualifier that here, the length includes any necessary padding that is necessary at sequence ends or where scaffolds are shorter than the specified subsequence length. The HMM is expected to scale linearly with the cumulative length of candidate gene regions. Testing this in practice, we observe approximately linear performance within phylogenetic groups, but slightly different coefficients between them, likely related to differences in typical gene length and number (Fig. 6).

**Figure 6:**
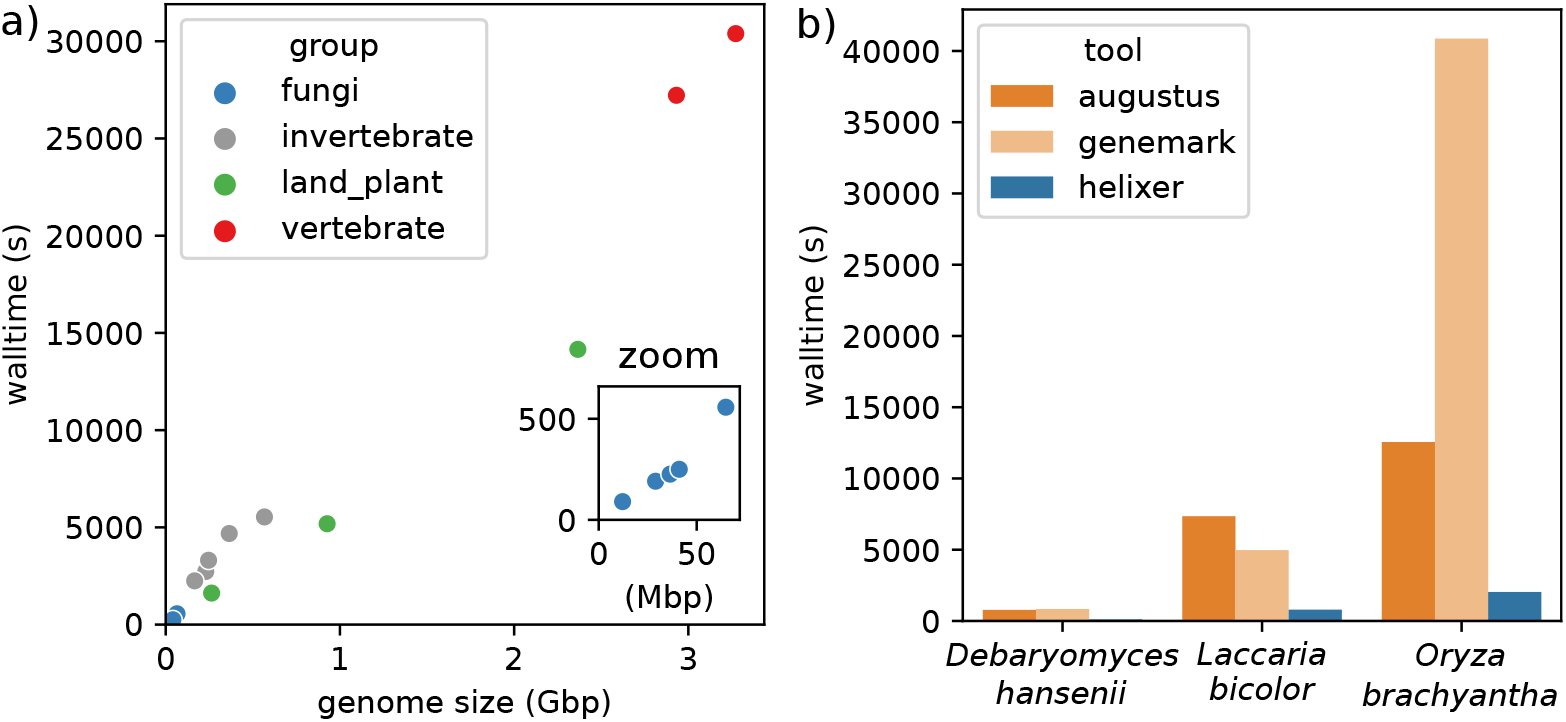
Benchmarking. a) Helixer walltime on test genomes and *Homo sapiens* on “workstation A”; b) comparison in single-threaded walltime between *de novo* tools

When comparing to GenemarkES and Augustus on the smallest and largest fungi test genomes, and smallest test plant genome, Helixer required 6.2-20.1 fold lower walltime in single threaded mode. However, we acknowledge the other tools have more complete multi-threading, and don’t require a graphics processing unit (GPU).

In single-threaded mode, Helixer can annotate the 263Mbp *Oryza brachyantha* genome in 27min, and the 3.3Gbp human genome in just under 8.5 hours (Fig. 6a). Exact speed will vary by system, particularly in regard to the speed of the HDD or SSD (Fig. S25), and of course the GPU (not shown).

## 5 Discussion

Helixer as a whole is the first Deep Learning-based tool for producing full eukaryotic gene models, and demonstrates what can be accomplished with a state-of-the-art *de novo* gene caller. The predicted primary gene models approach the quality of those references produced via data-supported pipelines and sometimes even manual curation, while requiring only a single tool, no data beyond the DNA sequence, and a very modest compute time. Although the overall quality does not yet match that of extrinsic data-supported pipelines, the production of a standardized output format allows for easy incorporation as part of any such pipeline; which has the potential to immediately improve the final quality of predictions. This is probable as we demonstrated that Helixer’s predictions are already able to find areas for improvement in major model species. Moreover, the resulting annotations have been gainfully applied for comparative genomics [Triesch et al., 2022]. The combined high-enough quality and also feasible-to-produce structural annotations, even for non-experts, are likely useful ‘as is’ for many projects.

While Helixer is the first fully applicable Deep Learning-based gene caller, it is not the pinnacle of what can be achieved by applying Deep Learning to structural gene annotation. We identify two major divisions for improvement, namely in modeling and data handling.

On the side of modeling, there are several promising options that should be explored in future work. For instance, end-to-end prediction is generally recommended to leverage the full power of Deep Learning. Others have shown that Deep Learning can effectively directly predict transitions such as splice sites [Zeng and Li, 2022, Zhang et al., 2016, Wang et al., 2019]. It is conceptually possible to encode a full gene structure as a series of transition tokens and positions, and then predict structure with a many-to-many model architecture such as those used for large language modeling [Brown et al., 2020, Vaswani et al., 2017]. This would have the advantage of being readily extensible for alternative splicing, but bring with it the challenge of an extremely sparse encoding. This could compound with erroneous or arbitrary labels (see below) and result in difficulties to get the model to converge during training. However, encoding the data as such would simplify application of major powerful techniques from language modeling, such as unsupervised pre-training [Devlin et al., 2018, Dai and Le, 2015]. Additionally, this would shorten the output length compared to base-wise encoding, and thus reduce the memory required for low-bias transformer architectures that have achieved extraordinary results across fields [Vaswani et al., 2017, Devlin et al., 2018, Avsec et al., 2021].

On the side of data, there are both extensive challenges and opportunities for improvement. Generally, there is no shortage of total data, but obtaining quality data for a phylogenetically broad and balanced selection of species is a major issue. We have every reason to think data quality is currently limiting network performance, for instance in Stiehler et al. [2020] (and here) we observed the lowest scoring predictions for UTR, notably where there was the highest discrepancy between references and independent RNAseq data was found, likely indicating errors in the reference annotations.

Several options exist to address data quality issues. The conceptually simplest – but poorly scaling – option is investing time, money and expertise specifically in improving the reference annotation quality of the training genomes. Indeed, effort is being continuously invested in improving reference annotations already. These improvements can be utilized simply by retraining Helixer, and automated selection of training species will promote adaptability to even large changes in available input data.

An intermediate option would be to use public extrinsic data for identifying and then differentially weighting regions of higher or lower quality in the reference and thereby reduce the noise the network sees while training.

Finally, there are pure modeling options such as the unsupervised pre-training mentioned above [Devlin et al., 2018, Dai and Le, 2015] or pseudo-labeling [Arazo et al., 2019]. These of course could also be combined, and an iterative approach where state-of-the-art *de novo* predictions from Helixer are incorporated into data-supported pipelines, to create state-of-the-art references that are in turn used to train a Deep Learning *de novo* caller would be an effort-intensive but highly reliable route to major improvement.

Naturally, there is also the option to train Deep Learning models to use RNAseq, CAGE or other extrinsic information as input. This is very likely to boost performance when both training and inference are performed with the highest quality of extrinsic data; but handling the potentially high variability of the extrinsic data at inference time will be an obstacle in its own right.

The above ideas are neither exhaustive, nor tested; however, we hope the potential to make further large gains in performance is clear. Perhaps most importantly, given the power of Deep Learning both shown here and impressively in other genomics tasks [Avsec et al., 2021, Jumper et al., 2021, Alipanahi et al., 2015, Zeng and Li, 2022], we hope that both researcher interest and available resources will start to reach a critical mass where modeling tasks previously considered untenable, such as achieving reference quality *de novo* annotations, can be accomplished. This will free up time and resources that previously flowed into both using more cumbersome tools, and moreover into large amounts of troubleshooting and manual validation, and accelerate research progress.

## Acknowledgments

Computational infrastructure and support were provided by the Centre for Information and Media Technology at Heinrich Heine University Düsseldorf. This work was supported by the Deutsche Forschungsgemeinschaft (DFG, German Research Foundation) under Germany’s Excellence Strategy–EXC-2048/1–Project ID: 390686111; by the DFG project 497667402; and by the BMBF-funded de.NBI Cloud within the German Network for Bioinformatics Infrastructure (de.NBI) (031A537B, 031A533A, 031A538A, 031A533B, 031A535A, 031A537C, 031A534A, 031A532B).

## Notes

### Competing Interest Statement

The authors have declared no competing interest.

### Summary of Updates

Fixed typos in affiliations. Fixed exact wording of affiliations & institute naming. More precise assignment of dual affiliations. Clean up extraneous '.' characters in title & affiliations. Updated relevant acknowledgement text to exactly match the HHU HPC's template.

