## Supplemental Information for "Helixer–*de novo* Prediction of Primary Eukaryotic Gene Models Combining Deep Learning and a Hidden Markov Model"

#### S1 Figures

#### S1.1 Network Architechture

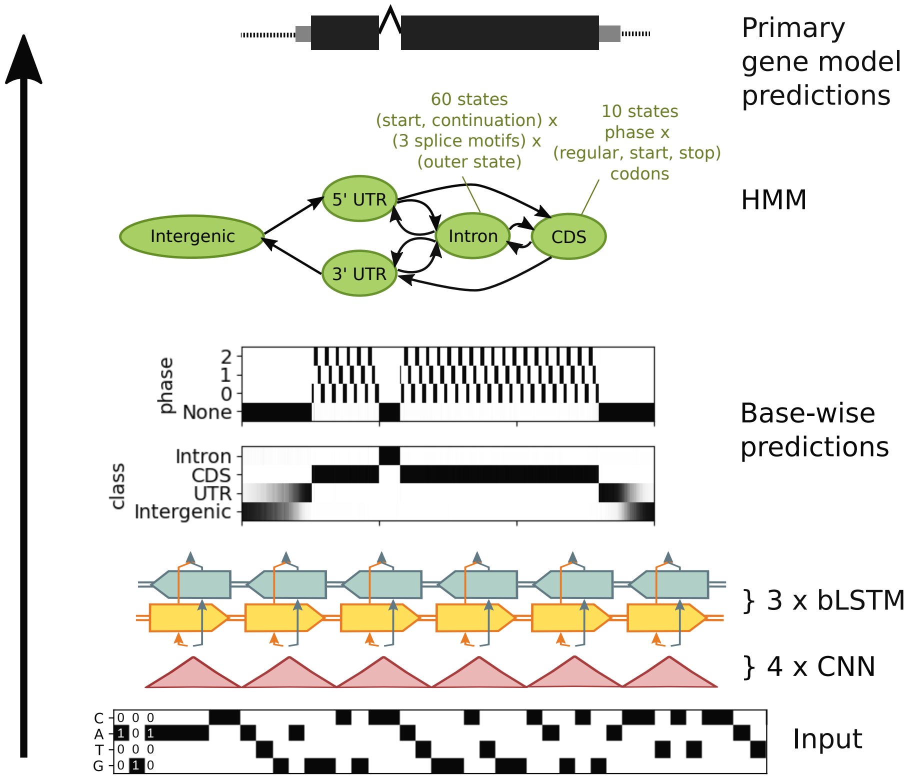


Figure S1: Visual summary of Helixer's predictive process. Not to scale. Hyperparameters are examples and may vary.

##### S1.2 Invertebrate, Plant and Fungi Model Comparison (Genic F1)

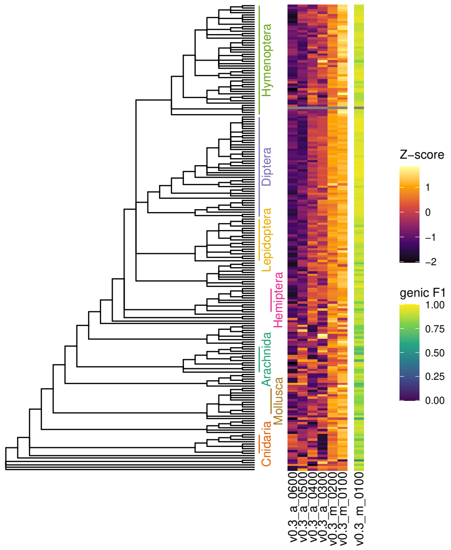


Figure S2: Performance of candidate and best invertebrate models across invertebrates. The absolute performance is measured with genic F1 on a random selection of 800 subsequences of each species and displayed for the overall best v0.3_m_0100 model (right). For peceptibility, differences between the genic F1 of models are displayed as a Z-score (middle).

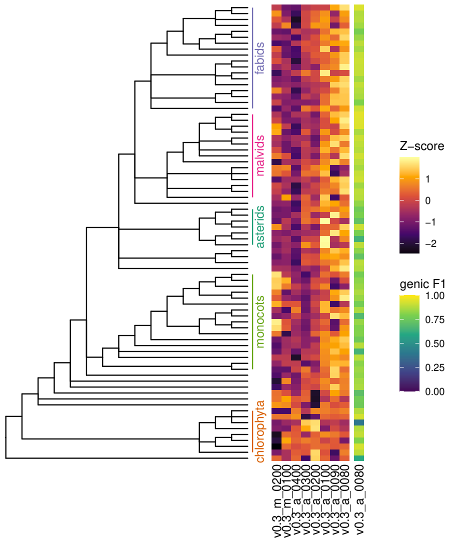


Figure S3: Performance of candidate and best land plant models across land plants. The absolute performance is measured with genic F1 on a random selection of 800 subsequences of each species and displayed for the overall best v0.3_a_0080 model (right). For peceptibility, differences between the genic F1 of models are displayed as a Z-score (middle). Models marked with * used the same training species as in Stiehler et al. (2020).

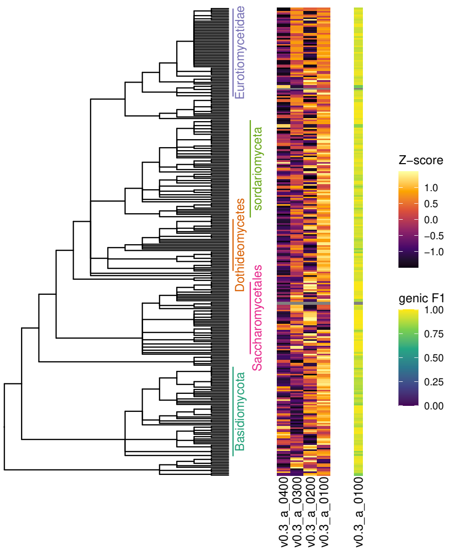


Figure S4: Performance of candidate and best fungi models across fungi. The absolute performance is measured with genic F1 on a random selection of 800 subsequences of each species and displayed for the overall best v0.3_a_0100 model (right). For peceptibility, differences between the genic F1 of models are displayed as a Z-score (middle).

#### S1.3 Vertebrate Performance With Species Labels

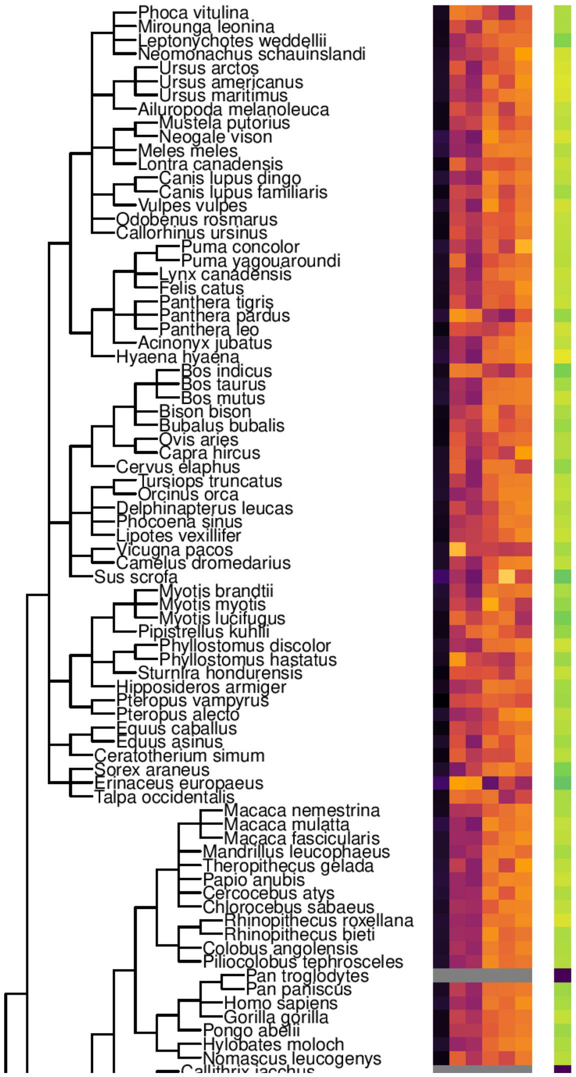
Phoca vitulina
Mirounga leonina
Leptonychotes weddellii
Neomonachus schauinslandi
Ursus arctos
Ursus americanus
Ursus maritimus
Ailuropoda melanoleuca
Mustela putorius
Neogale vison
Meles meles
Lontra canadensis
Canis lupus dingo
Canis lupus familiaris
Vulpes vulpes
Odobenus rosmarus
Callorhinus ursinus
Puma concolor
Puma yagouaroundi
Lynx canadensis
Felis catus
Panthera tigris
Panthera pardus
Panthera leo
Acinonyx jubatus
Hyaena hyaena
Bos indicus
Bos taurus
Bos mutus
Bison bison
Bubalus bubalis
Ovis aries
Capra hircus
Cervus elaphus
Tursiops truncatus
Orcinus orca
Delphinapterus leucas
Phocoena sinus
Lipotes vexillifer
Vicugna pacos
Camelus dromedarius
Sus scrofa
Myotis brandtii
Myotis myotis
Myotis lucifugus
Pipistrellus kuhlii
Phyllostomus discolor
Phyllostomus hastatus
Sturnira hondurensis
Hipposideros armiger
Pteropus vampyrus
Pteropus alecto
Equus caballus
Equus asinus
Ceratotherium simum
Sorex araneus
Erinaceus europaeus
Talpa occidentalis
Macaca nemestrina
Macaca mulatta
Macaca fascicularis
Mandrillus leucophaeus
Theropithecus gelada
Papio anubis
Cercocebus atys
Chlorocebus sabaeus
Rhinopithecus roxellana
Rhinopithecus bieti
Colobus angolensis
Piliocolobus tephrosceles
Pan troglodytes
Pan paniscus
Homo sapiens
Gorilla gorilla
Pongo abelii
Hylobates moloch
Nomascus leucogenys

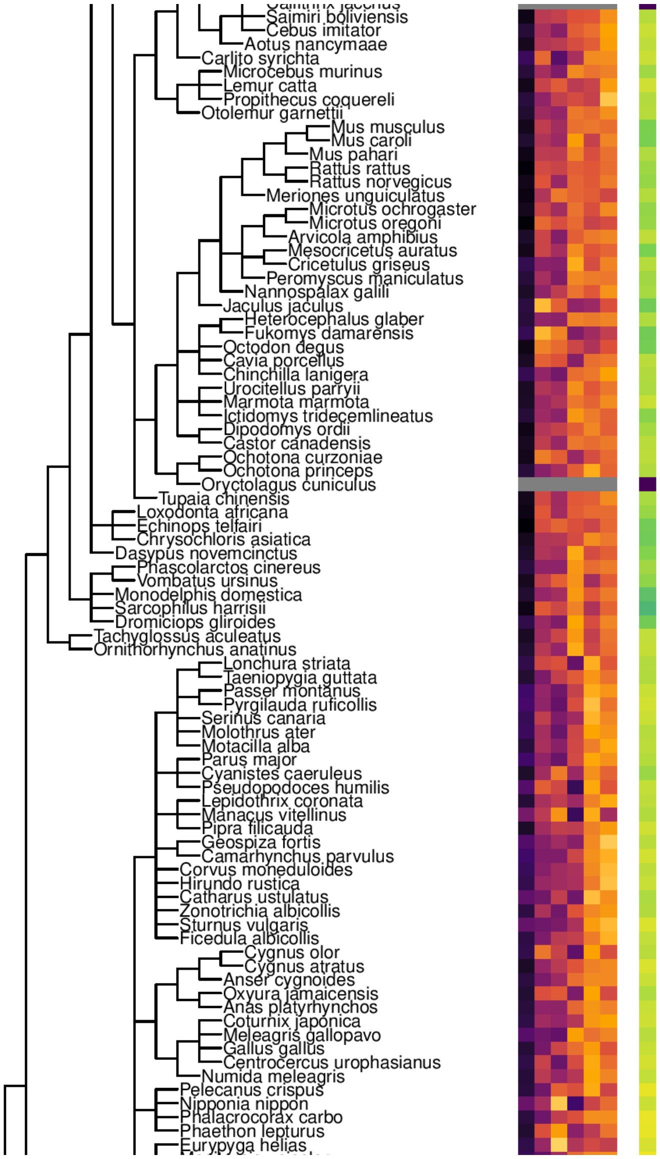
Saimiri boliviensis
Cebus imitator
Aotus nancymaae
Carlito syrichta
Microcebus murinus
Lemur catta
Propithecus coquereli
Otolemur garnettii
Mus musculus
Mus caroli
Mus pahari
Rattus rattus
Rattus norvegicus
Meriones unguiculatus
Microtus ochrogaster
Microtus oregoni
Arvicola amphibius
Mesocricetus auratus
Cricetulus griseus
Peromyscus maniculatus
Nannospalax galili
Jaculus jaculus
Heterocephalus glaber
Fukomys damarensis
Octodon degus
Cavia porcellus
Chinchilla lanigera
Urocitellus parryii
Marmota marmota
Ictidomys tridecemlineatus
Dipodomys ordii
Castor canadensis
Ochotona curzoniae
Ochotona princeps
Oryctolagus cuniculus
Tupaia chinensis
Loxodonta africana
Echinops telfairi
Chrysochloris asiatica
Dasypus novemcinctus
Phascolarctos cinereus
Vombatus ursinus
Monodelphis domestica
Sarcophilus harrisii
Dromiciops gliroides
Tachyglossus aculeatus
Ornithorhynchus anatinus
Lonchura striata
Taeniopygia guttata
Passer montanus
Pyrgilauda ruficollis
Serinus canaria
Molothrus ater
Motacilla alba
Parus major
Cyanistes caeruleus
Pseudopodoces humilis
Lepidothrix coronata
Manacus vitellinus
Pipra filicauda
Geospiza fortis
Camarhynchus parvulus
Corvus moneduloides
Hirundo rustica
Catharus ustulatus
Zonotrichia albicollis
Sturnus vulgaris
Ficedula albicollis
Cygnus olor
Cygnus atratus
Anser cygnoides
Oxyura jamaicensis
Anas platyrhynchos
Coturnix japonica
Meleagris gallopavo
Gallus gallus
Centrocercus urophasianus
Numida meleagris
Pelecanus crispus
Nipponia nippon
Phalacrocorax carbo
Phaethon lepturus
Eurypyga helias

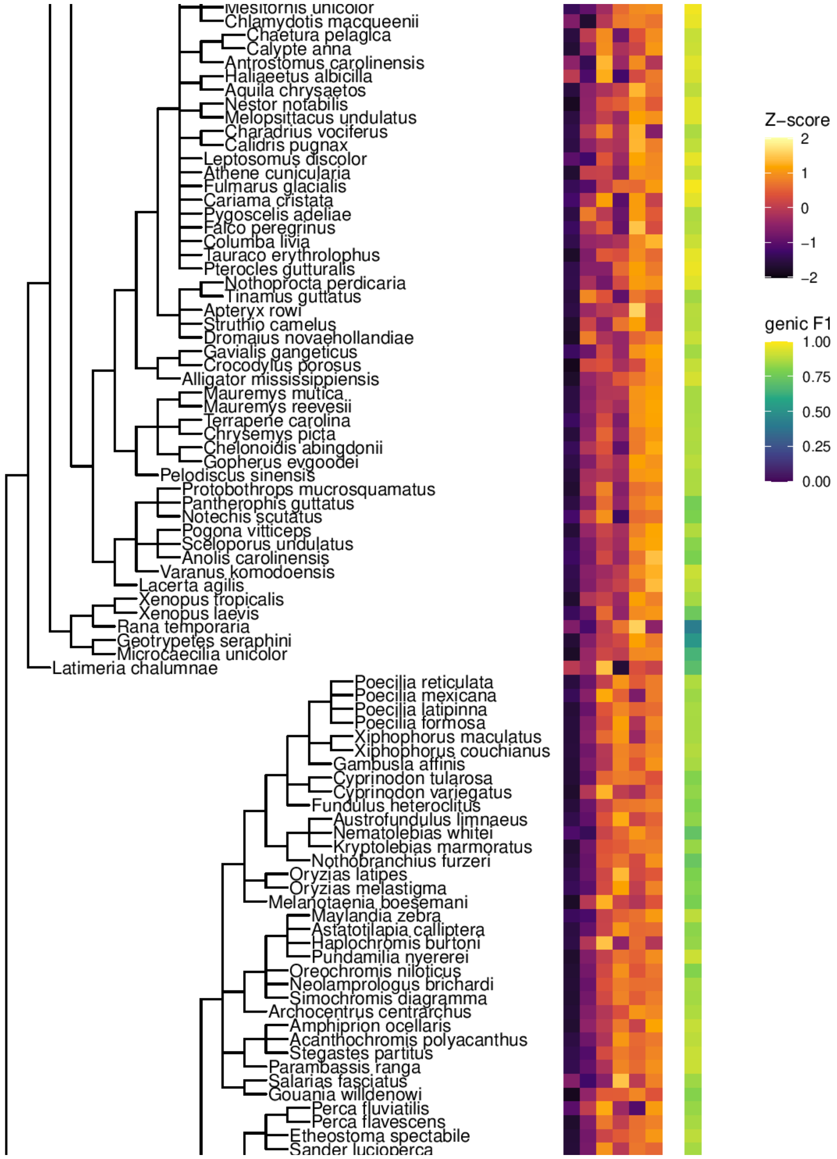
Chlamydotis macqueenii
Chaetura pelagica
Calypte anna
Antrostomus carolinensis
Haliaeetus albicilla
Aquila chrysaetos
Nestor notabilis
Melopsittacus undulatus
Charadrius vociferus
Calidris pugnax
Leptosomus discolor
Athene cunicularia
Fulmarus glacialis
Cariama cristata
Pygoscelis adeliae
Falco peregrinus
Columba livia
Tauraco erythrolophus
Pterocles gutturalis
Nothoprocta perdicaria
Tinamus guttatus
Apteryx rowi
Struthio camelus
Dromaius novaehollandiae
Gavialis gangeticus
Crocodylus porosus
Alligator mississippiensis
Mauremys mutica
Mauremys reevesii
Terrapene carolina
Chrysemys picta
Chelonoidis abingdonii
Gopherus evgoodei
Pelodiscus sinensis
Protobothrops mucrosquamatus
Pantherophis guttatus
Notechis scutatus
Pogona vitticeps
Sceloporus undulatus
Anolis carolinensis
Varanus komodoensis
Lacerta agilis
Xenopus tropicalis
Xenopus laevis
Rana temporaria
Geotrypetes seraphini
Microcaecilia unicolor
Latimeria chalumnae
Poecilia reticulata
Poecilia mexicana
Poecilia latipinna
Poecilia formosa
Xiphophorus maculatus
Xiphophorus couchianus
Gambusia affinis
Cyprinodon tularosa
Cyprinodon variegatus
Fundulus heteroclitus
Austrofundulus limnaeus
Nematolebias whitei
Kryptolebias marmoratus
Nothobranchius furzeri
Oryzias latipes
Oryzias melastigma
Melanotaenia boesemani
Maylandia zebra
Astatotilapia calliptera
Haplochromis burtoni
Pundamilia nyererei
Oreochromis niloticus
Neolamprologus brichardi
Simochromis diagramma
Archocentrus centrarchus
Amphiprion ocellaris
Acanthochromis polyacanthus
Stegastes partitus
Parambassis ranga
Salarias fasciatus
Gouania willdenowi
Perca fluviatilis
Perca flavescens
Etheostoma spectabile
Sander lucioperca
Z−score
2
1
0
−1
−2
genic F1
1.00
0.75
0.50
0.25
0.00

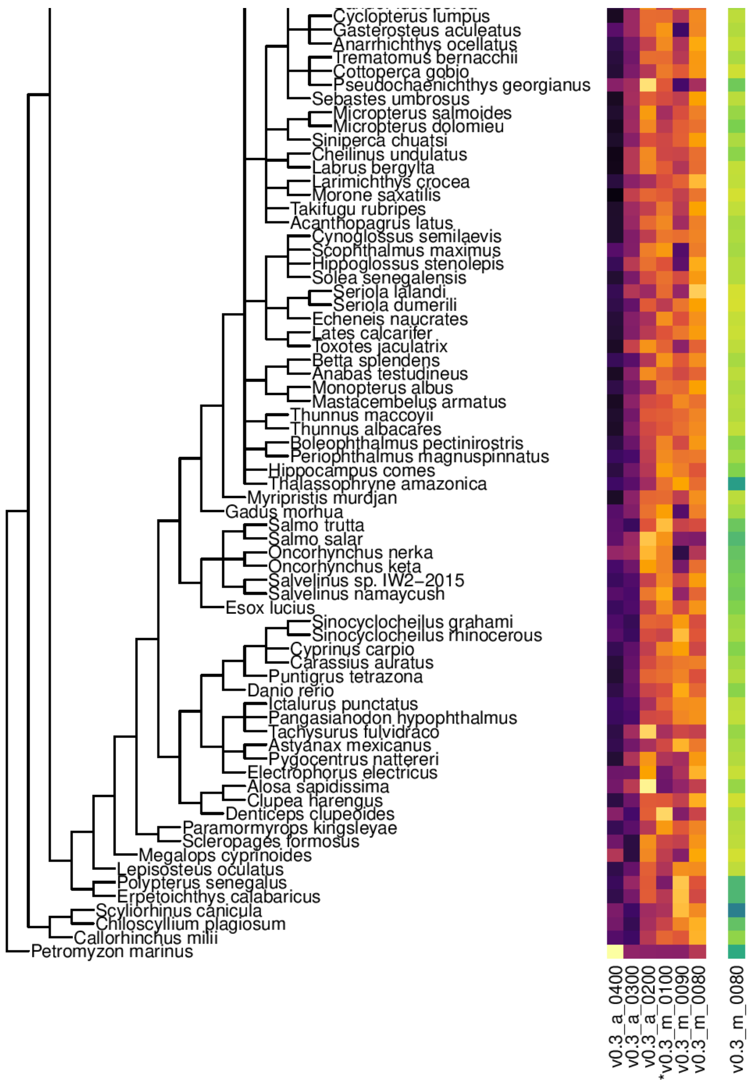


Figure S5: Performance of candidate and best vertebrate models across vertebrates. The absolute performance is measured with genic F1 on a random selection of 800 subsequences of each species and displayed for the overall best v0.3_m_0080 model (right). For peceptibility, differences between the genic F1 of models are displayed as a Z-score (middle). Models marked with * used the same training species as in Stiehler et al. (2020).

#### S1.4 Invertebrate Performance With Species Labels

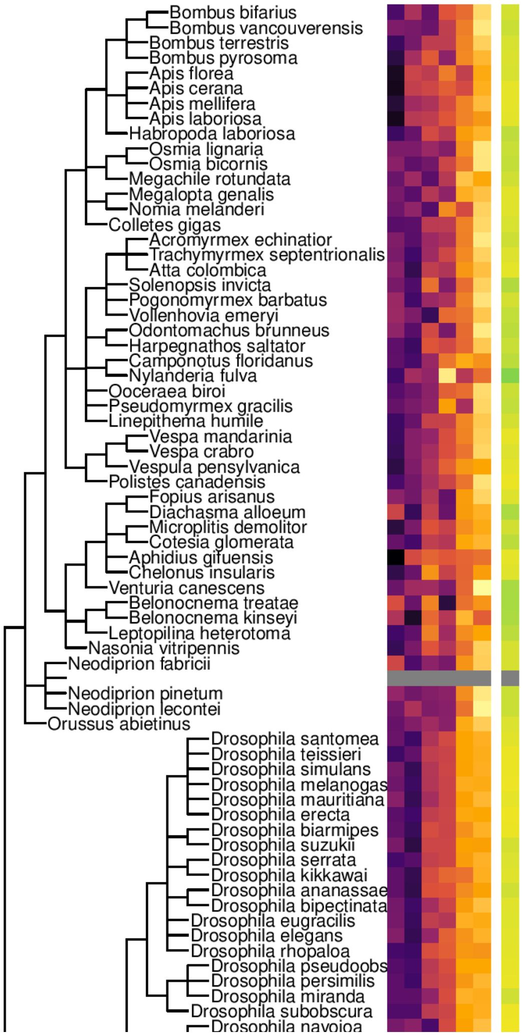
Bombus bifarius
Bombus vancouverensis
Bombus terrestris
Bombus pyrosoma
Apis florea
Apis cerana
Apis mellifera
Apis laboriosa
Habropoda laboriosa
Osmia lignaria
Osmia bicornis
Megachile rotundata
Megalopta genalis
Nomia melanderi
Colletes gigas
Acromyrmex echinatior
Trachymyrmex septentrionalis
Atta colombica
Solenopsis invicta
Pogonomyrmex barbatus
Vollenhovia emeryi
Odontomachus brunneus
Harpegnathos saltator
Camponotus floridanus
Nylanderia fulva
Ooceraea biroi
Pseudomyrmex gracilis
Linepithema humile
Vespa mandarinia
Vespa crabro
Vespula pensylvanica
Polistes canadensis
Fopius arisanus
Diachasma alloeum
Microplitis demolitor
Cotesia glomerata
Aphidius gifuensis
Chelonus insularis
Venturia canescens
Belonocnema treatae
Belonocnema kinseyi
Leptopilina heterotoma
Nasonia vitripennis
Neodiprion fabricii
Neodiprion pinetum
Neodiprion lecontei
Orussus abietinus
Drosophila santomea
Drosophila teissieri
Drosophila simulans
Drosophila melanogas
Drosophila mauritiana
Drosophila erecta
Drosophila biarmipes
Drosophila suzukii
Drosophila serrata
Drosophila kikkawai
Drosophila ananassae
Drosophila bipectinata
Drosophila eugracilis
Drosophila elegans
Drosophila rhopaloa
Drosophila pseudoobs
Drosophila persimilis
Drosophila miranda
Drosophila subobscura
Drosophila navojoa

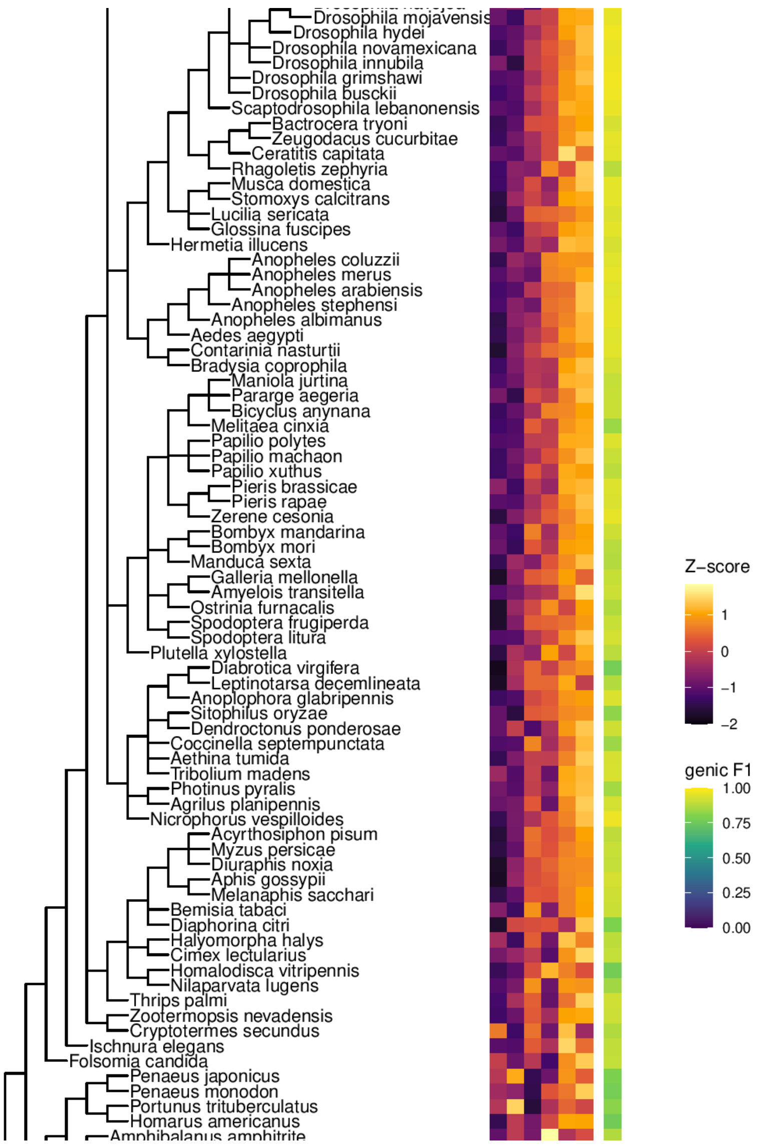
Drosophila mojavensis
Drosophila hydei
Drosophila novamexicana
Drosophila innubila
Drosophila grimshawi
Drosophila busckii
Scaptodrosophila lebanonensis
Bactrocera tryoni
Zeugodacus cucurbitae
Ceratitis capitata
Rhagoletis zephyria
Musca domestica
Stomoxys calcitrans
Lucilia sericata
Glossina fuscipes
Hermetia illucens
Anopheles coluzzii
Anopheles merus
Anopheles arabiensis
Anopheles stephensi
Anopheles albimanus
Aedes aegypti
Contarinia nasturtii
Bradysia coprophila
Maniola jurtina
Pararge aegeria
Bicyclus anynana
Melitaea cinxia
Papilio polytes
Papilio machaon
Papilio xuthus
Pieris brassicae
Pieris rapae
Zerene cesonia
Bombyx mandarina
Bombyx mori
Manduca sexta
Galleria mellonella
Amyelois transitella
Ostrinia furnacalis
Spodoptera frugiperda
Spodoptera litura
Plutella xylostella
Diabrotica virgifera
Leptinotarsa decemlineata
Anoplophora glabripennis
Sitophilus oryzae
Dendroctonus ponderosae
Coccinella septempunctata
Aethina tumida
Tribolium madens
Photinus pyralis
Agrilus planipennis
Nicrophorus vespilloides
Acyrthosiphon pisum
Myzus persicae
Diuraphis noxia
Aphis gossypii
Melanaphis sacchari
Bemisia tabaci
Diaphorina citri
Halyomorpha halys
Cimex lectularius
Homalodisca vitripennis
Nilaparvata lugens
Thrips palmi
Zootermopsis nevadensis
Cryptotermes secundus
Ischnura elegans
Folsomia candida
Penaeus japonicus
Penaeus monodon
Portunus trituberculatus
Homarus americanus
Amphibalanus amphitrite
Z−score
1
0
−1
−2
genic F1
1.00
0.75
0.50
0.25
0.00

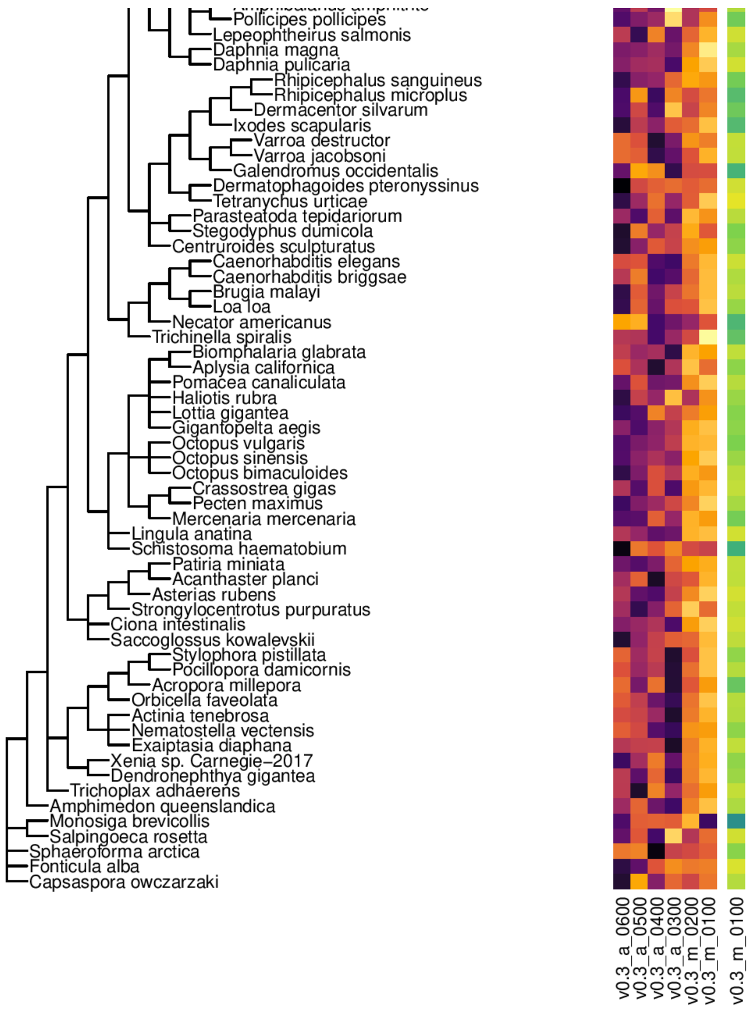


Figure S6: Performance of candidate and best invertebrate models across invertebrates. The absolute performance is measured with genic F1 on a random selection of 800 subsequences of each species and displayed for the overall best v0.3_m_0100 model (right). For peceptibility, differences between the genic F1 of models are displayed as a Z-score (middle).

##### S1.5 Plant Performance With Species Labels

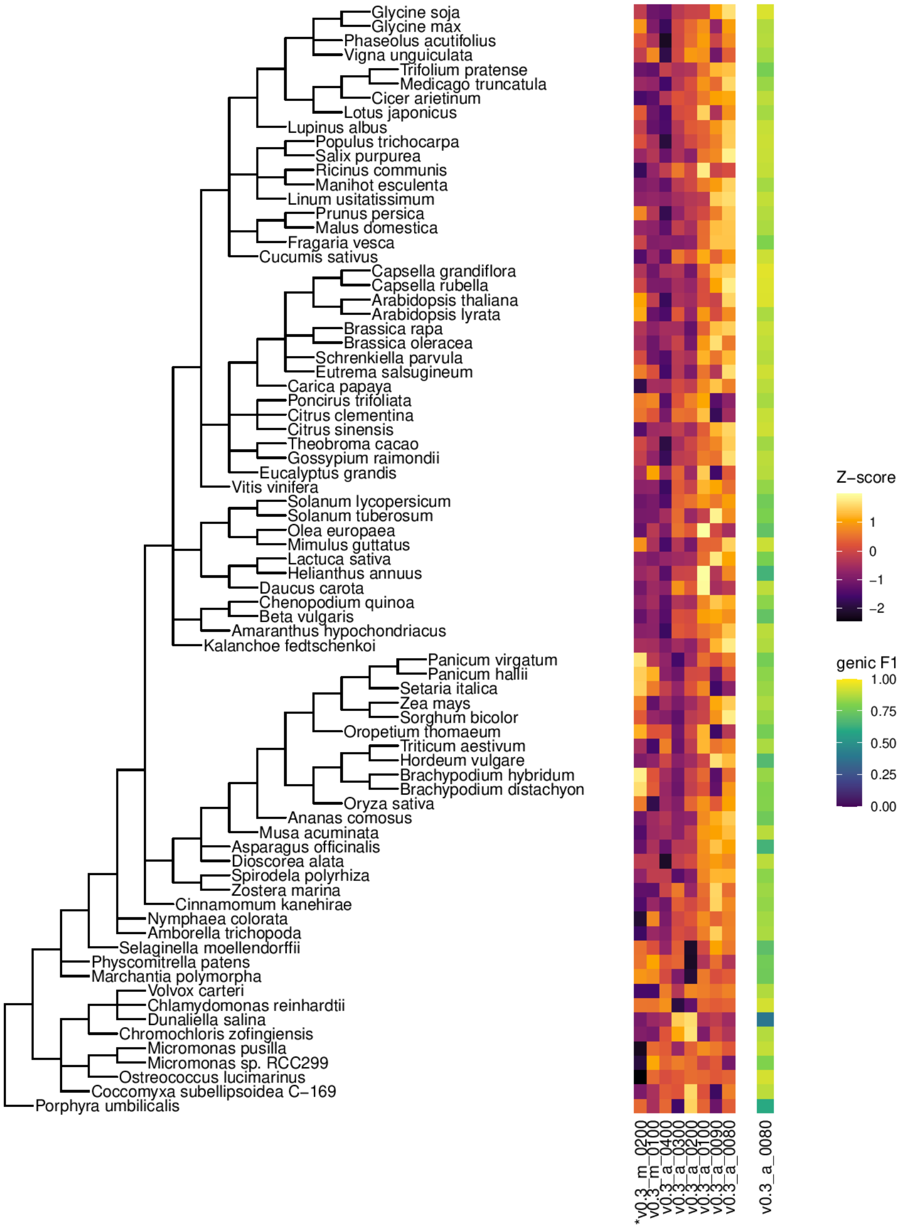


Figure S7: Performance of candidate and best land plant models across land plants. The absolute performance is measured with genic F1 on a random selection of 800 subsequences of each species and displayed for the overall best v0.3_m_0080 model (right). For peceptibility, differences between the genic F1 of models are displayed as a Z-score (middle). Models marked with * used the same training species as in Stiehler et al. (2020).

##### S1.6 Fungi Performance With Species Labels

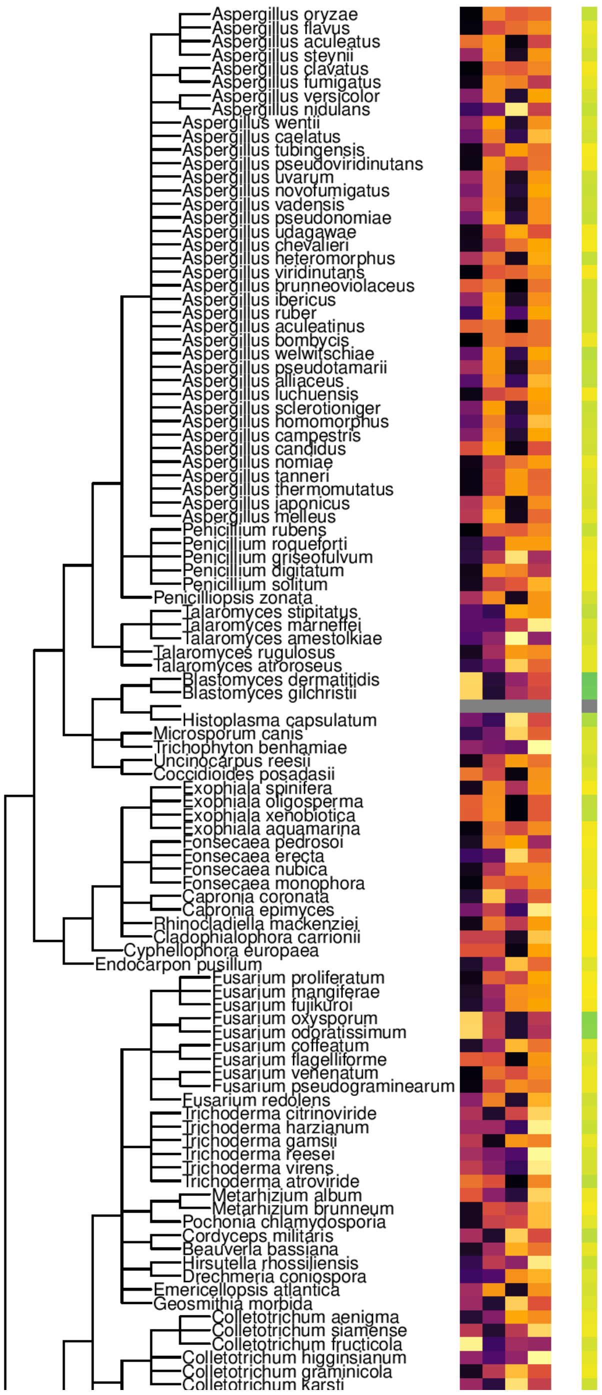
Aspergillus oryzae
Aspergillus flavus
Aspergillus aculeatus
Aspergillus steynii
Aspergillus clavatus
Aspergillus fumigatus
Aspergillus versicolor
Aspergillus nidulans
Aspergillus wentii
Aspergillus caelatus
Aspergillus tubingensis
Aspergillus pseudoviridinutans
Aspergillus uvarum
Aspergillus novofumigatus
Aspergillus vadensis
Aspergillus pseudonomiae
Aspergillus udagawae
Aspergillus chevalieri
Aspergillus heteromorphus
Aspergillus viridinutans
Aspergillus brunneoviolaceus
Aspergillus ibericus
Aspergillus ruber
Aspergillus aculeatinus
Aspergillus bombycis
Aspergillus welwitschiae
Aspergillus pseudotamarii
Aspergillus alliaceus
Aspergillus luchuensis
Aspergillus sclerotioniger
Aspergillus homomorphus
Aspergillus campestris
Aspergillus candidus
Aspergillus nomiae
Aspergillus tanneri
Aspergillus thermomutatus
Aspergillus japonicus
Aspergillus melleus
Penicillium rubens
Penicillium roqueforti
Penicillium griseofulvum
Penicillium digitatum
Penicillium solitum
Penicilliopsis zonata
Talaromyces stipitatus
Talaromyces marneffei
Talaromyces amestolkiae
Talaromyces rugulosus
Talaromyces atroroseus
Blastomyces dermatitidis
Blastomyces gilchristii
Histoplasma capsulatum
Microsporum canis
Trichophyton benhamiae
Uncinocarpus reesii
Coccidioides posadasii
Exophiala spinifera
Exophiala oligosperma
Exophiala xenobiotica
Exophiala aquamarina
Fonsecaea pedrosoi
Fonsecaea erecta
Fonsecaea nubica
Fonsecaea monophora
Capronia coronata
Capronia epimyces
Rhinocladiella mackenziei
Cladophialophora carrionii
Cyphellophora europaea
Endocarpon pusillum
Fusarium proliferatum
Fusarium mangiferae
Fusarium fujikuroi
Fusarium oxysporum
Fusarium odoratissimum
Fusarium coffeatum
Fusarium flagelliforme
Fusarium venenatum
Fusarium pseudograminearum
Fusarium redolens
Trichoderma citrinoviride
Trichoderma harzianum
Trichoderma gamsii
Trichoderma reesei
Trichoderma virens
Trichoderma atroviride
Metarhizium album
Metarhizium brunneum
Pochonia chlamydosporia
Cordyceps militaris
Beauveria bassiana
Hirsutella rhossiliensis
Drechmeria coniospora
Emericellopsis atlantica
Geosmithia morbida
Colletotrichum aenigma
Colletotrichum siamense
Colletotrichum fructicola
Colletotrichum higginsianum
Colletotrichum graminicola
Colletotrichum karsti

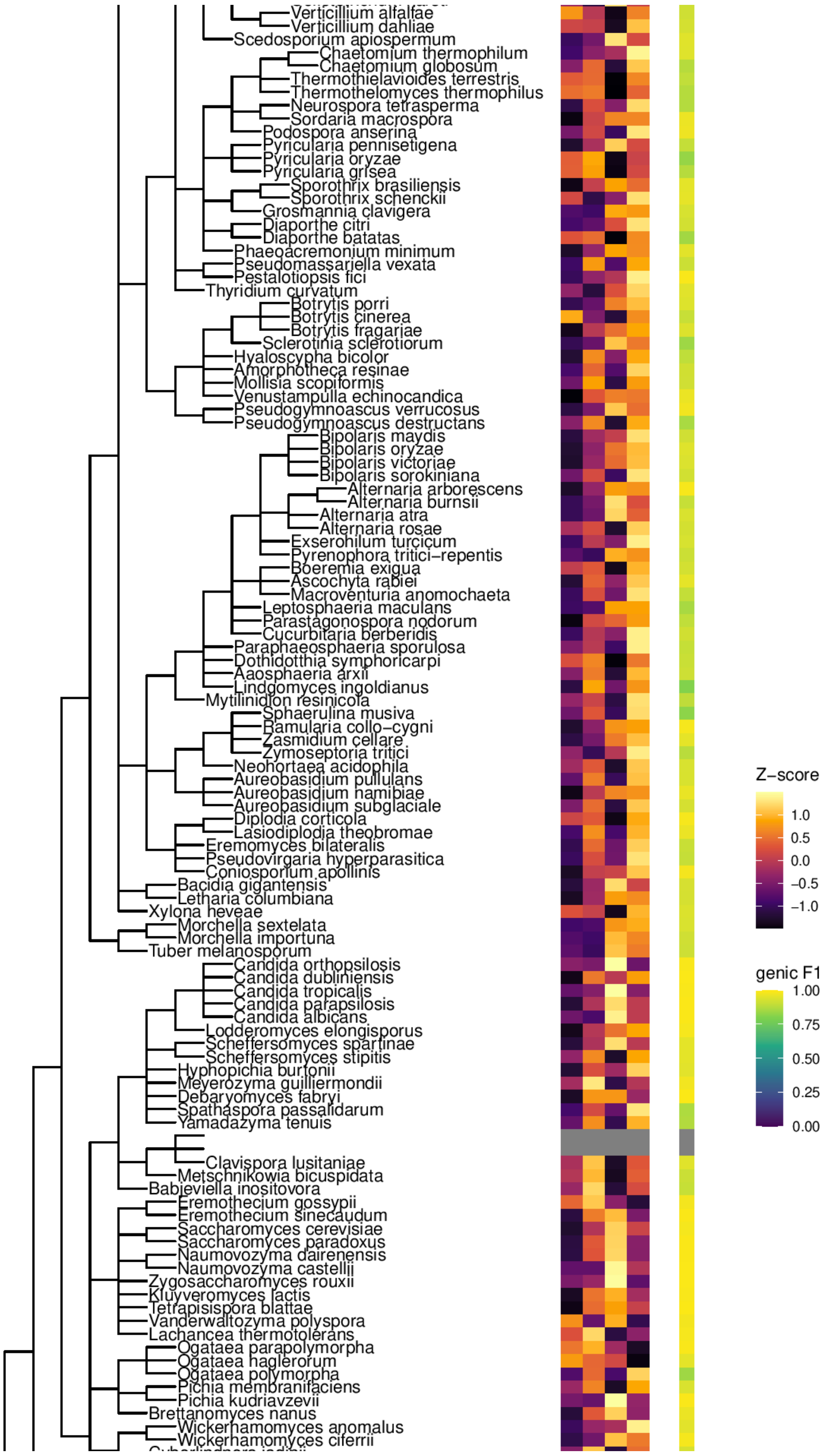
Verticillium alfalfae
Verticillium dahliae
Scedosporium apiospermum
Chaetomium thermophilum
Chaetomium globosum
Thermothielavioides terrestris
Thermothelomyces thermophilus
Neurospora tetrasperma
Sordaria macrospora
Podospora anserina
Pyricularia pennisetigena
Pyricularia oryzae
Pyricularia grisea
Sporothrix brasiliensis
Sporothrix schenckii
Grosmannia clavigera
Diaporthe citri
Diaporthe batatas
Phaeoacremonium minimum
Pseudomassariella vexata
Pestalotiopsis fici
Thyridium curvatum
Botrytis porri
Botrytis cinerea
Botrytis fragariae
Sclerotinia sclerotiorum
Hyaloscypha bicolor
Amorphotheca resinae
Mollisia scopiformis
Venustampulla echinocandica
Pseudogymnoascus verrucosus
Pseudogymnoascus destructans
Bipolaris maydis
Bipolaris oryzae
Bipolaris victoriae
Bipolaris sorokiniana
Alternaria arborescens
Alternaria burnsii
Alternaria atra
Alternaria rosae
Exserohilum turcicum
Pyrenophora tritici−repentis
Boeremia exigua
Ascochyta rabiei
Macroventuria anomochaeta
Leptosphaeria maculans
Parastagonospora nodorum
Cucurbitaria berberidis
Paraphaeosphaeria sporulosa
Dothidotthia symphoricarpi
Aaosphaeria arxii
Lindgomyces ingoldianus
Mytilinidion resinicola
Sphaerulina musiva
Ramularia collo−cygni
Zasmidium cellare
Zymoseptoria tritici
Neohortaea acidophila
Aureobasidium pullulans
Aureobasidium namibiae
Aureobasidium subglaciale
Diplodia corticola
Lasiodiplodia theobromae
Eremomyces bilateralis
Pseudovirgaria hyperparasitica
Coniosporium apollinis
Bacidia gigantensis
Letharia columbiana
Xylona heveae
Morchella sextelata
Morchella importuna
Tuber melanosporum
Candida orthopsilosis
Candida dubliniensis
Candida tropicalis
Candida parapsilosis
Candida albicans
Lodderomyces elongisporus
Scheffersomyces spartinae
Scheffersomyces stipitis
Hyphopichia burtonii
Meyerozyma guilliermondii
Debaryomyces fabryi
Spathaspora passalidarum
Yamadazyma tenuis
Clavispora lusitaniae
Metschnikowia bicuspidata
Babjeviella inositovora
Eremothecium gossypii
Eremothecium sinecaudum
Saccharomyces cerevisiae
Saccharomyces paradoxus
Naumovozyma dairenensis
Naumovozyma castellii
Zygosaccharomyces rouxii
Kluyveromyces lactis
Tetrapisispora blattae
Vanderwaltozyma polyspora
Lachancea thermotolerans
Ogataea parapolymorpha
Ogataea haglerorum
Ogataea polymorpha
Pichia membranifaciens
Pichia kudriavzevii
Brettanomyces nanus
Wickerhamomyces anomalus
Wickerhamomyces ciferrii
Z−score
1.0
0.5
0.0
−0.5
−1.0
genic F1
1.00
0.75
0.50
0.25
0.00

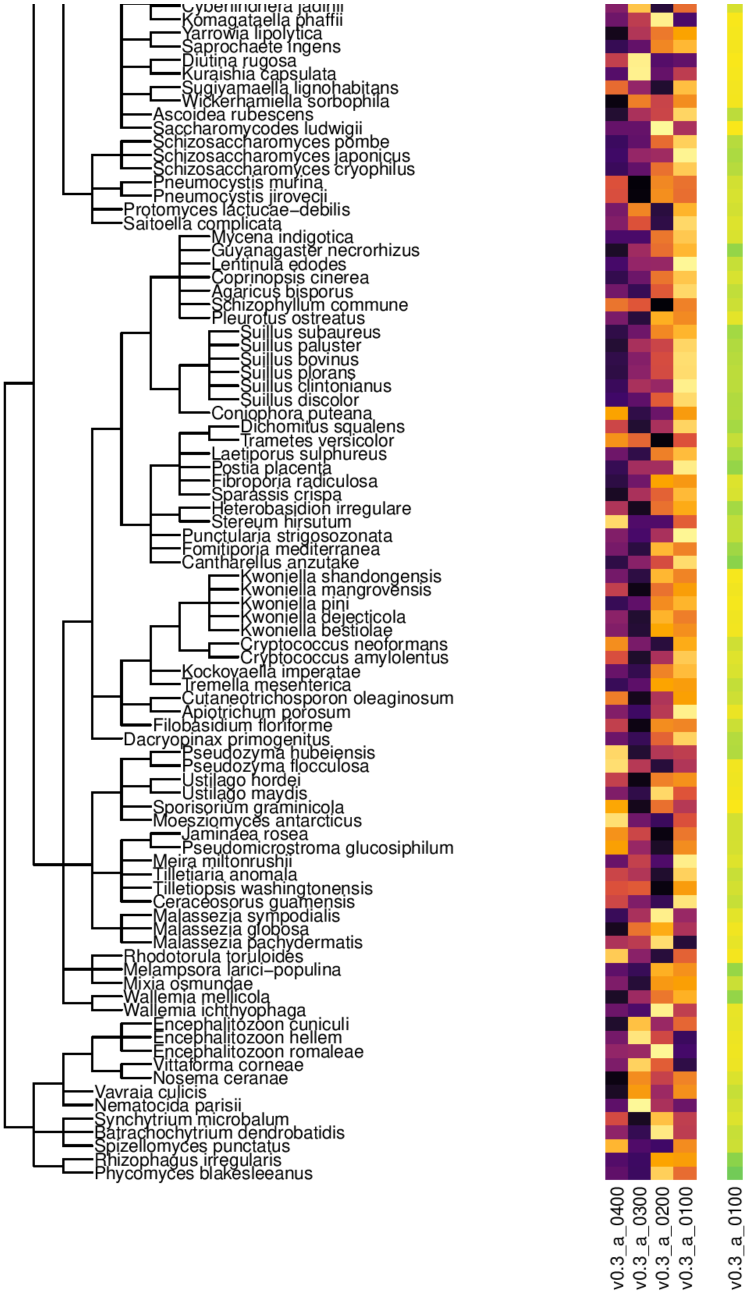


Figure S8: Performance of candidate and best fungi models across fungi. The absolute performance is measured with genic F1 on a random selection of 800 subsequences of each species and displayed for the overall best v0.3_a_0100 model (right). For peceptibility, differences between the genic F1 of models are displayed as a Z-score (middle).

##### S1.7 Relative Performance of Annotations

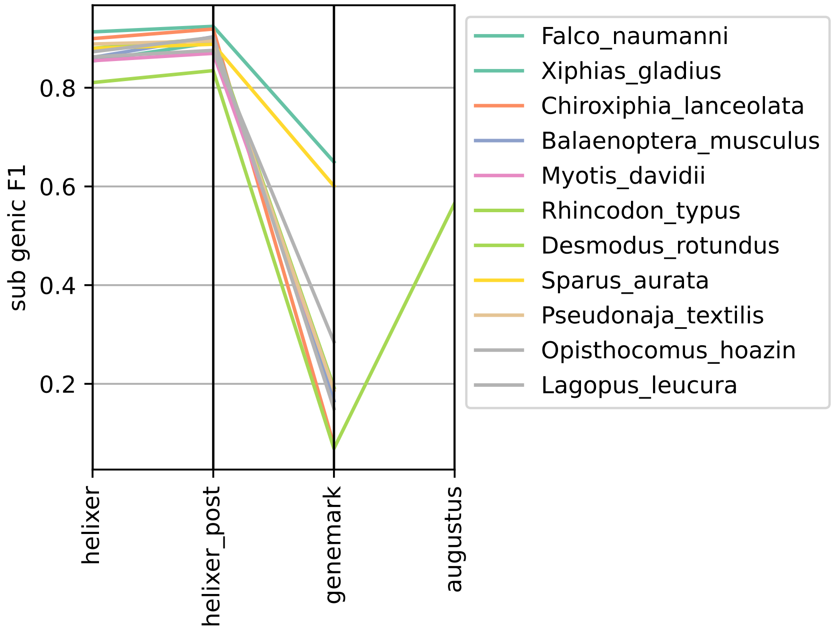


Figure S9: Test set annotation quality for vertebrates measured as Subgenic F1 compared to the reference

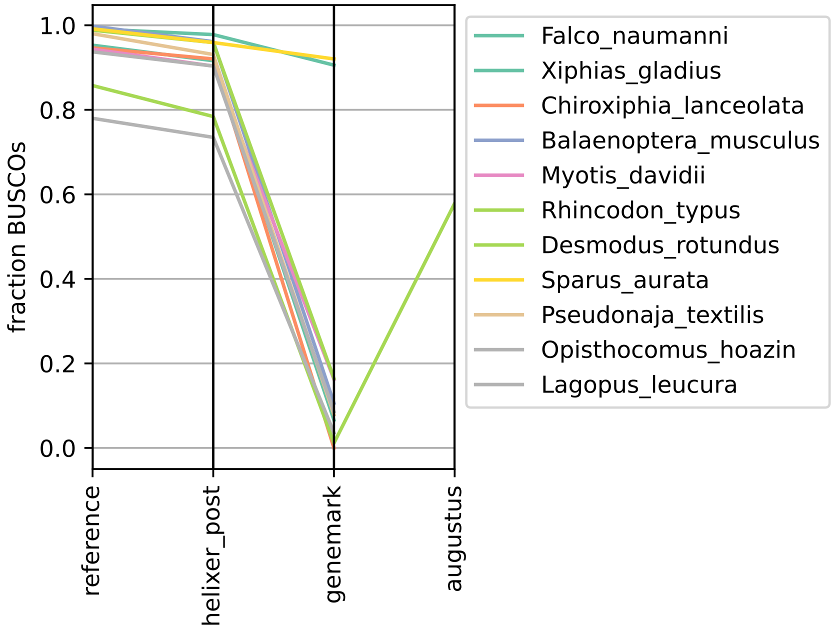


Figure S10: Test set annotation quality for vertebrates measured as fraction of complete BUSCOs found in final annotation.

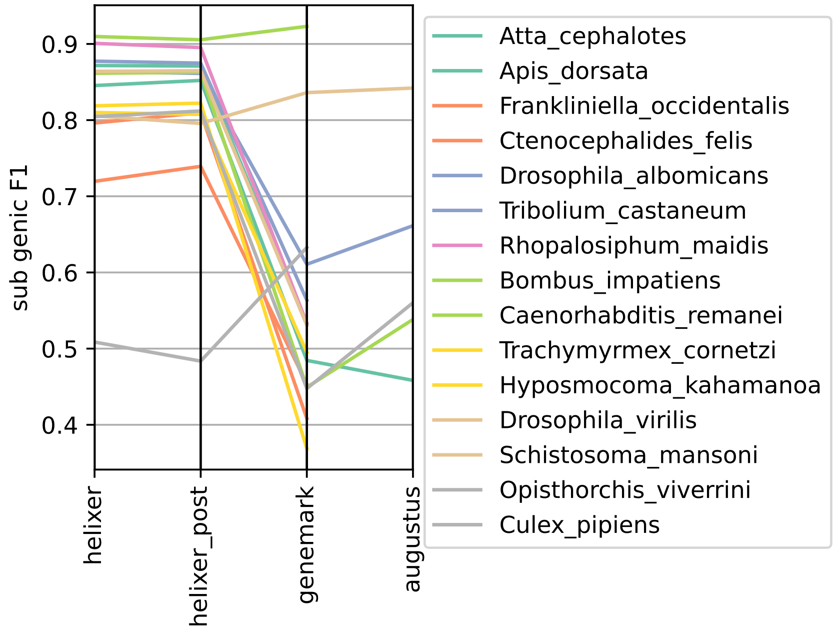


Figure S11: Test set annotation quality for invertebrates measured as Subgenic F1 compared to the reference

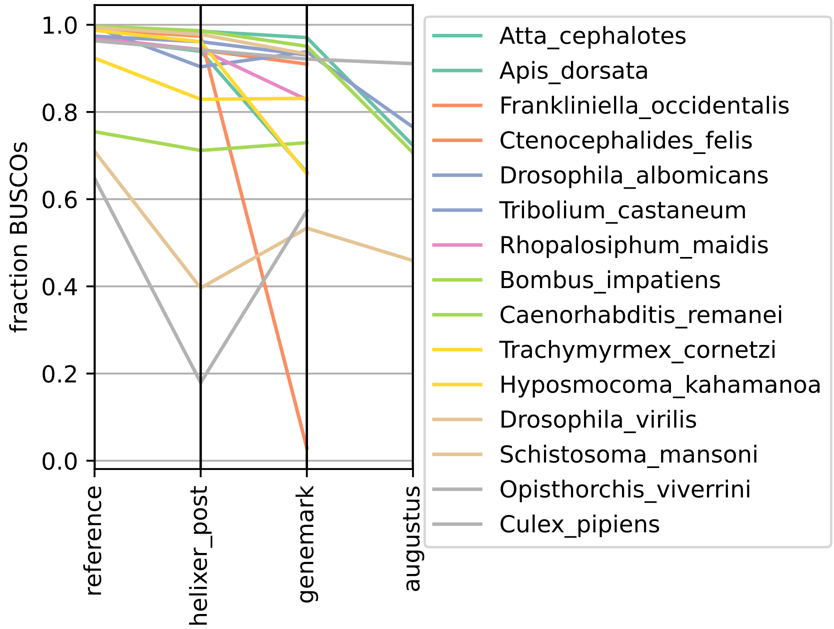


Figure S12: Test set annotation quality for invertebrates measured as fraction of complete BUSCOs found in final annotation.

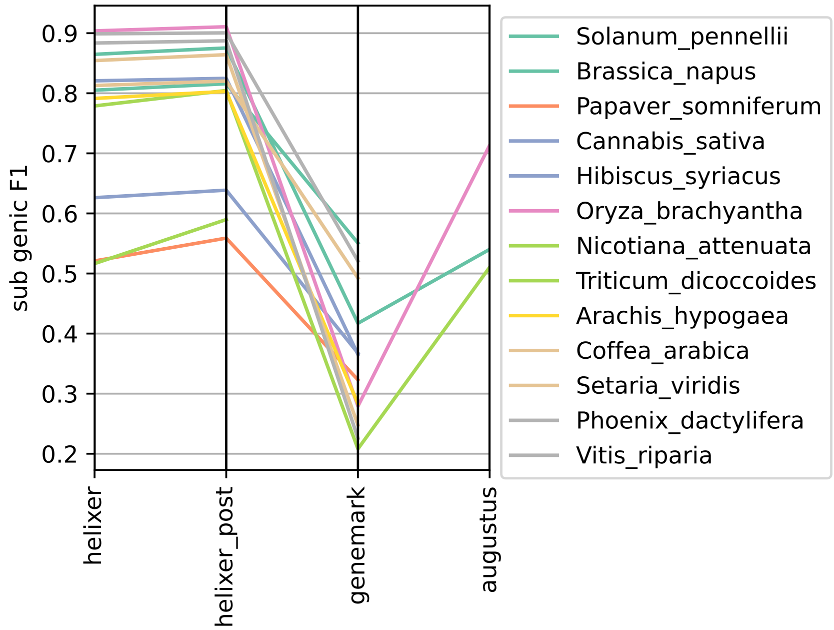


Figure S13: Test set annotation quality for plants measured as Subgenic F1 compared to the reference

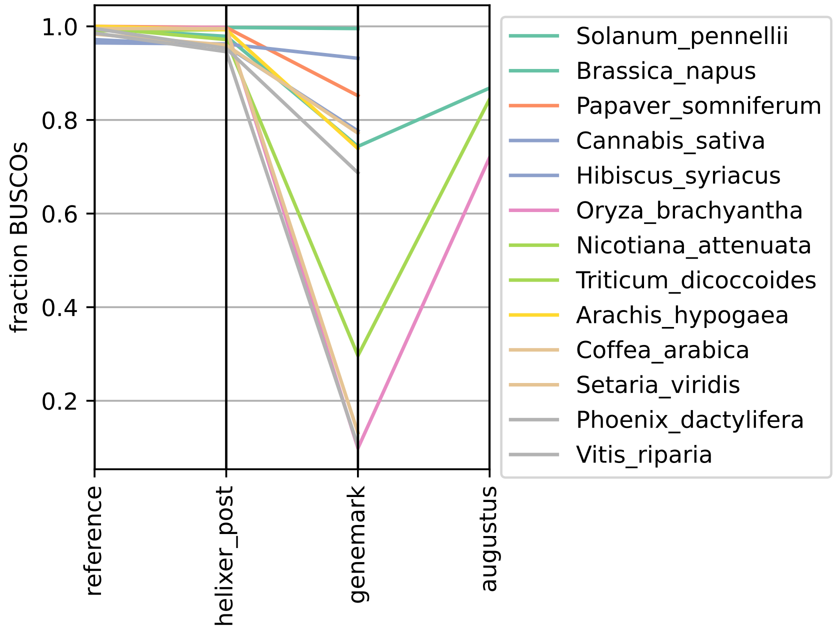


Figure S14: Test set annotation quality for plants measured as fraction of complete BUSCOs found in final annotation.

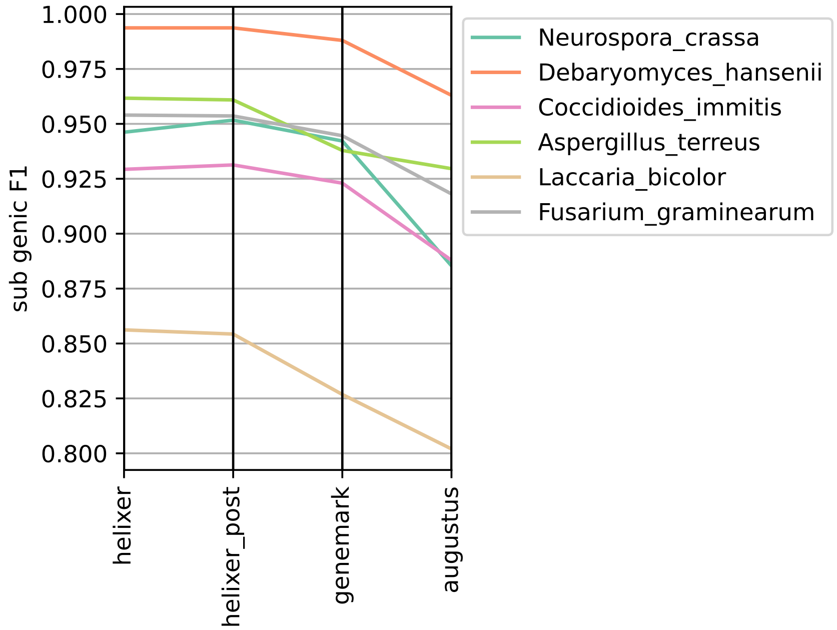


Figure S15: Test set annotation quality for fungis measured as Subgenic F1 compared to the reference

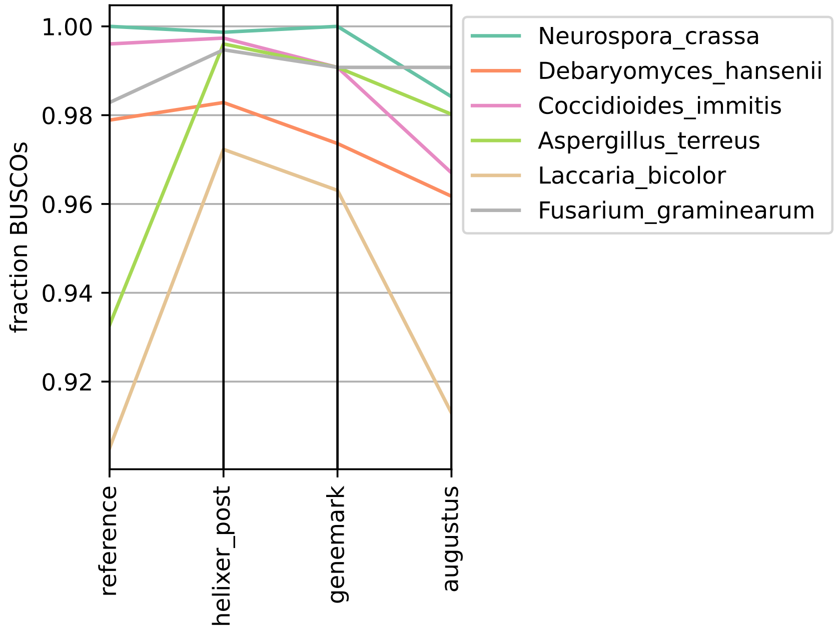


Figure S16: Test set annotation quality for fungis measured as fraction of complete BUSCOs found in final annotation.

#### S1.8 Orthogroup based quality control

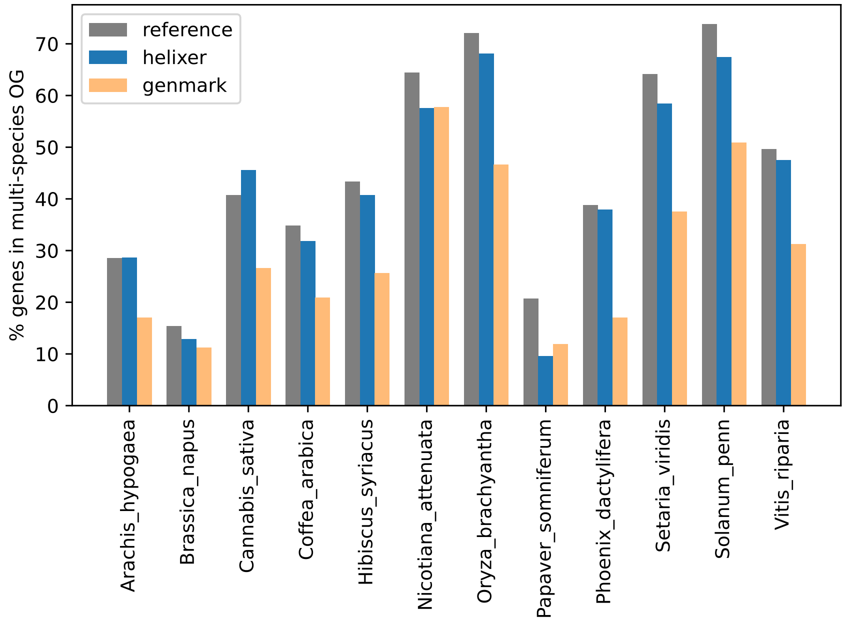


Figure S17: Precision proxy of proteome annotations as estimated with orthogroups. Specifically, the percentage of genes from each species that were assigned to an orthogroup containing at least one other species.

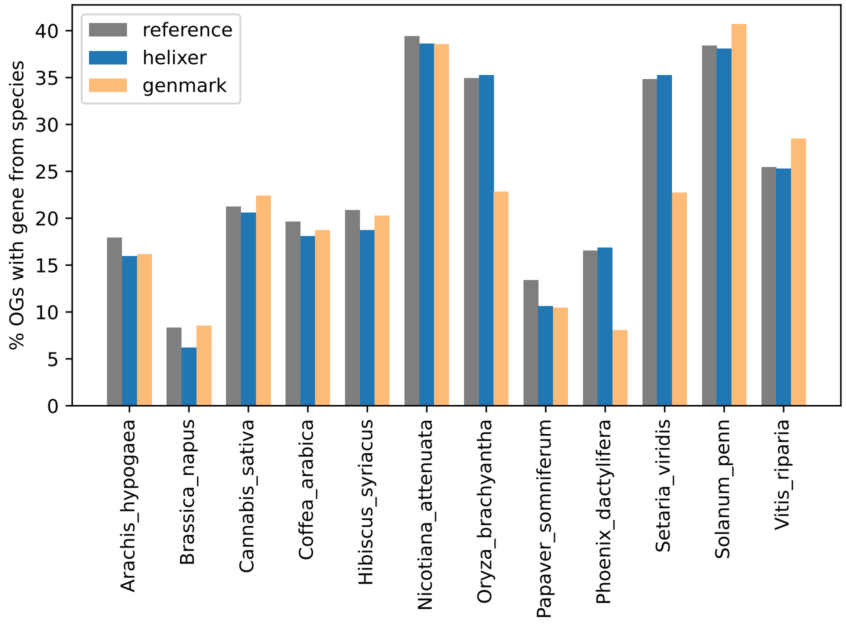


Figure S18: Recall proxy of proteome annotations as estimated with orthogroups. Specifically the percentage of orthogroups that contain a gene from each given species.

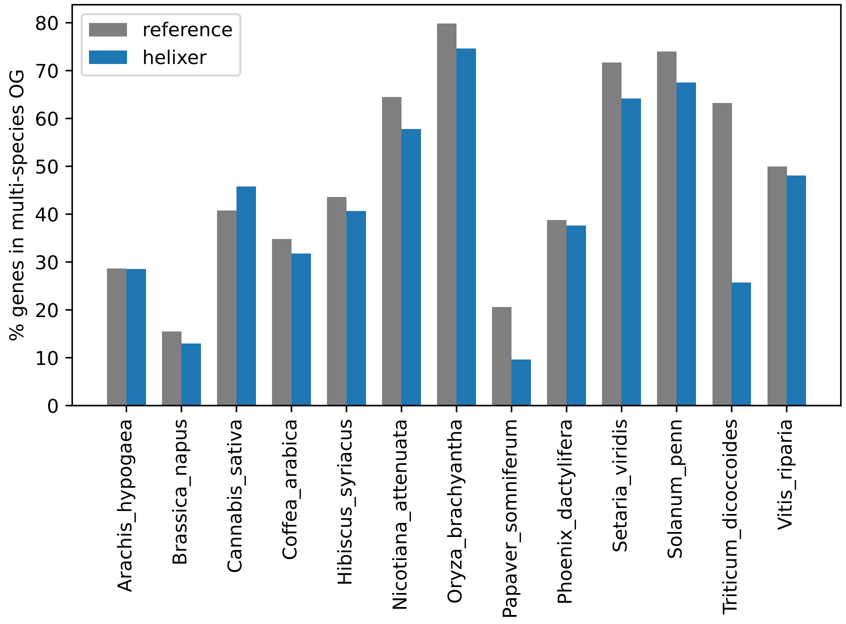


Figure S19: Precision proxy of proteome annotations as estimated with orthogroups. Specifically, the percentage of genes from each species that were assigned to an orthogroup containing at least one other species. Here with all 13 plant test species, but without GenemarkES (missing annotation for *T. dicoccoides*).

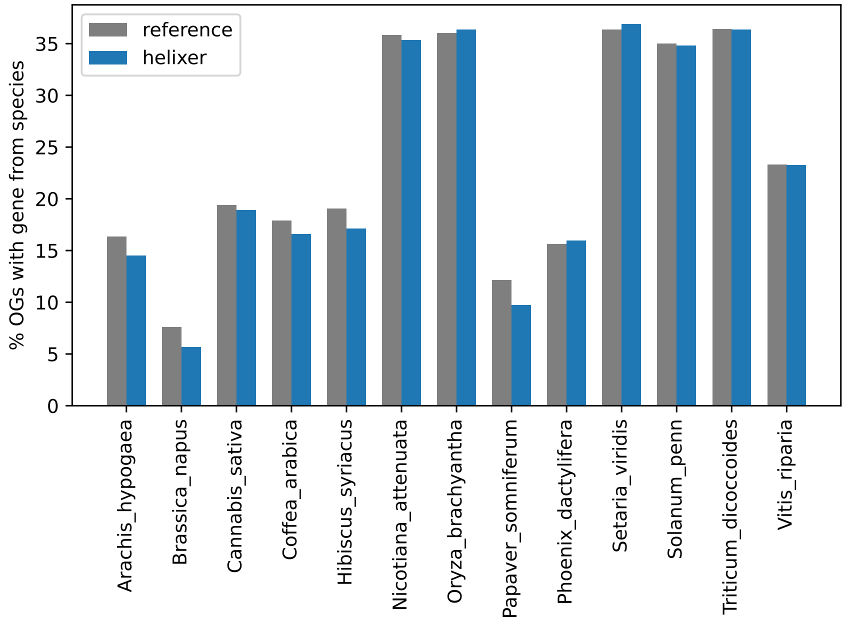


Figure S20: Recall proxy of proteome annotations as estimated with orthogroups. Specifically the percentage of orthogroups that contain a gene from each given species. Here with all 13 plant test species, but without GenemarkES (missing annotation for *T. dicoccoides*)

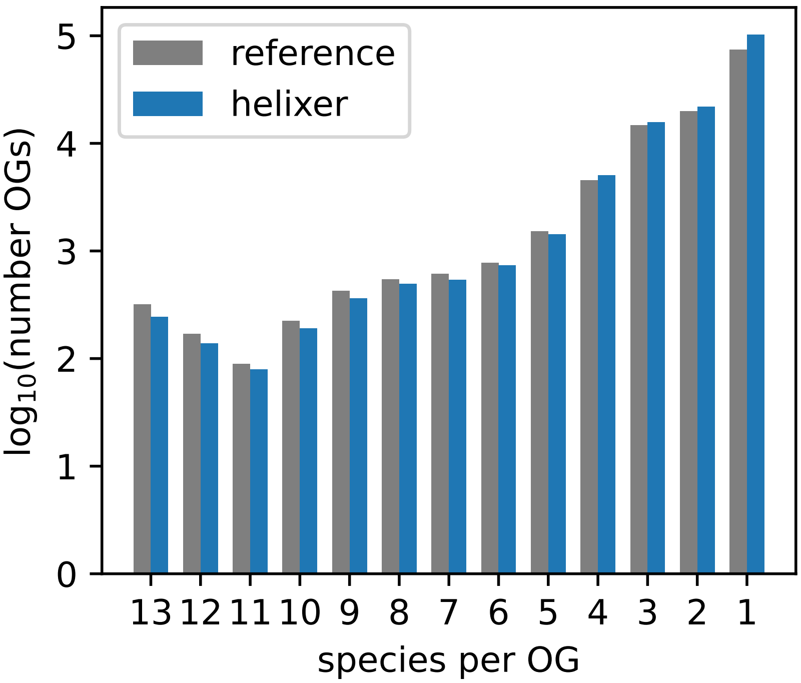


Figure S21: Orthogroup occupancy (number of species represented out of 13) for orthogroups based on the reference and Helixer's annotations (each clustered individually)

##### S1.9 *Arabidopsis thaliana*

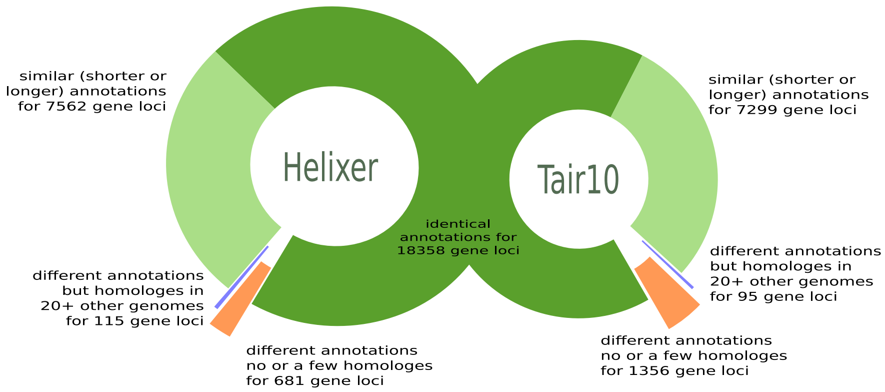


Figure S22: Comparison of the *A. thaliana* proteome predicted by Helixer to the existing TAIR10 annotation. A breakdown of the overlaps and differences between predicted proteomes.

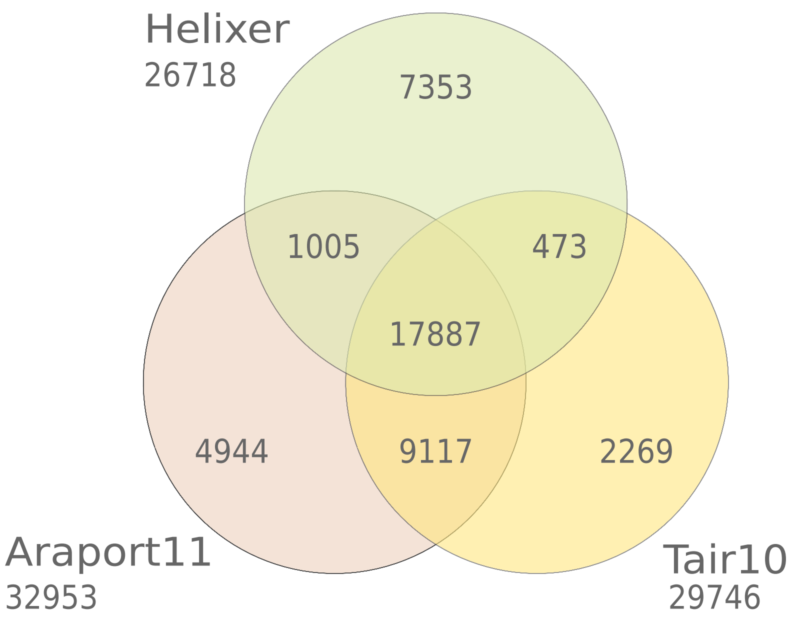


Figure S23: Comparison of the *A. thaliana* proteome predicted by Helixer to the existing TAIR10 and Araport11 annotations. Venn diagram of *identical* proteins. Note that here all splice variants, not loci, are counted.

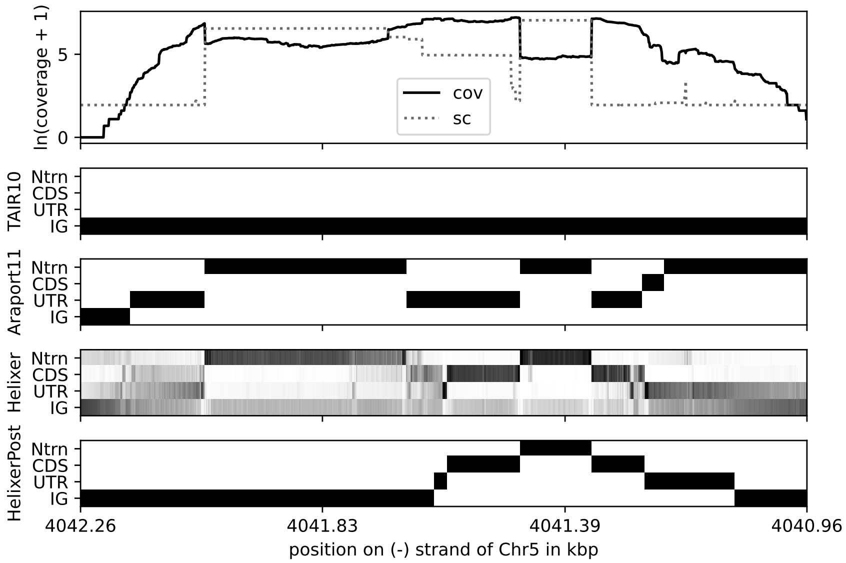


Figure S24: Comparison of the *A. thaliana* proteome predicted by Helixer to the existing TAIR10 and Araport11 annotations. Visualization of expression and available annotations zoomed into the genomic locus of the Phosphatidylinositol N-acetylglucosaminyltransferase $\gamma$ subunit (Chr5, - strand, from 4041091 to 4041627, via HelixerPost).

##### S1.10 Benchmarking

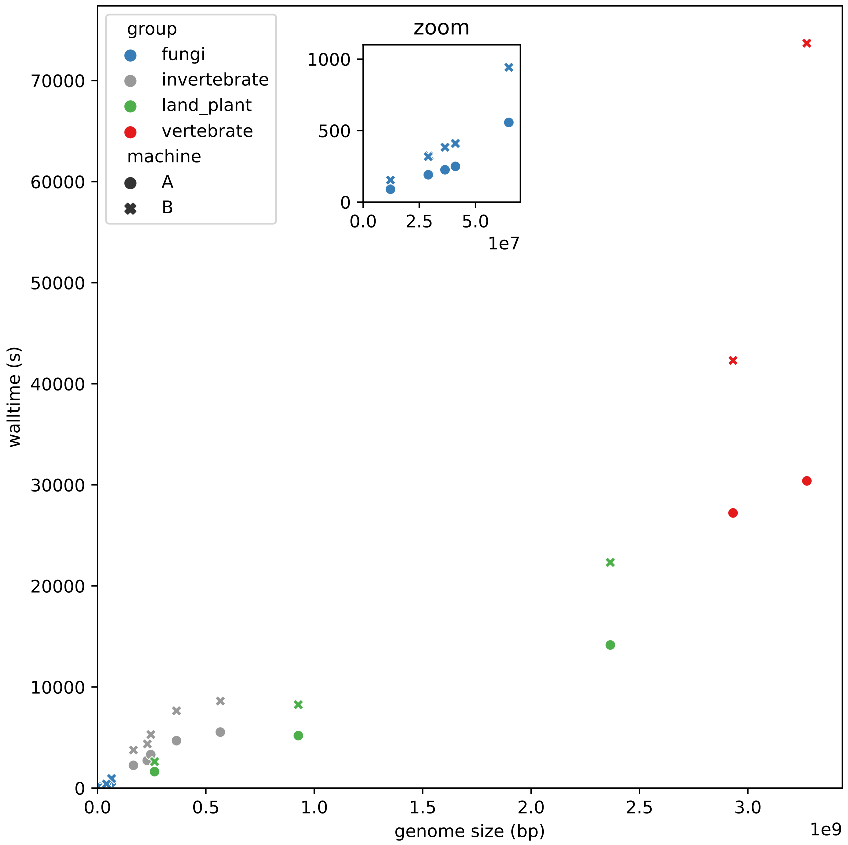


Figure S25: Benchmarking. Helixer walltime on some test genomes and *H. sapiens*. Comparison between two different workstations (see methods).

### S2 Tables

#### S2.1 Full Datasets

Due to the iterative nature of development, datasets were acquired from different sources at different times, as listed below

##### S2.1.1 Fungi

Fungi data for training, validation, and testing were acquired from RefSeq on March 4th, 2022. Test genomes were assigned randomly *within* genomes that had a matching or close AUGUSTUS trained model available and *within* those that did not.

All species assigned to test that had a close AUGUSTUS model were actually used as test species in this manuscript, all other test species remain reserved.

Table S1: All fungi genomes, set assignment and versions

| scientific | ID | version | split |
|---|---|---|---|
| Aaosphaeria arxii | Aaosphaeria_arxii | GCF_010015735.1_Aaoar1 | train_val |
| Acaromyces ingoldii | Acaromyces_ingoldii | GCF_003144295.1_Acain1 | test |
| Agaricus bisporus | Agaricus_bisporus | GCF_000300575.1_Agabi_varbisH97_2 | train_val |
| Alternaria alternata | Alternaria_alternata | GCF_001642055.1_Altal1 | test |
| Alternaria arborescens | Alternaria_arborescens | GCF_004154835.1_ASM415483v1 | train_val |
| Alternaria atra | Alternaria_atra | GCF_907166805.1_ALTATR162 | train_val |
| Alternaria burnsii | Alternaria_burnsii | GCF_013036055.1_ASM1303605v1 | train_val |
| Alternaria rosae | Alternaria_rosae | GCF_020736505.1_Altro1 | train_val |
| Amorphotheca resinae | Amorphotheca_resinae | GCF_003019875.1_Amore1 | train_val |
| Apiotrichum porosum | Apiotrichum_porosum | GCF_003942205.1_ASM394220v1 | train_val |
| Aplosporella prunicola | Aplosporella_prunicola | GCF_010093885.1_Aplpr1 | test |
| Arthroderma uncinatum | Arthroderma_uncinatum | GCF_011692745.1_ASM1169274v1 | test |
| Ascochyta rabiei | Ascochyta_rabiei | GCF_004011695.1_Arabiei_Me14 | train_val |
| Ascoidea rubescens | Ascoidea_rubescens | GCF_001661345.1_Ascru1 | train_val |
| Aspergillus aculeatinus | Aspergillus_aculeatinus | GCF_003184765.1_Aspacu1 | train_val |
| Aspergillus aculeatus | Aspergillus_aculeatus | GCF_001890905.1_Aspac1 | train_val |
| Aspergillus alliaceus | Aspergillus_alliaceus | GCF_009176365.1_Aspalli1 | train_val |
| Aspergillus bombycis | Aspergillus_bombycis | GCF_001792695.1_ASM179269v1 | train_val |
| Aspergillus brunneoviolaceus | Aspergillus_brunneoviolaceus | GCF_003184695.1_Aspbru1 | train_val |
| Aspergillus caelatus | Aspergillus_caelatus | GCF_009193585.1_Aspcae1 | train_val |
| Aspergillus campestris | Aspergillus_campestris | GCF_002847485.1_Aspcam1 | train_val |
| Aspergillus candidus | Aspergillus_candidus | GCF_002847045.1_Aspcand1 | train_val |
| Aspergillus chevalieri | Aspergillus_chevalieri | GCF_016861735.1_AchevalieriM1_assembly01 | train_val |
| Aspergillus clavatus | Aspergillus_clavatus | GCF_000002715.2_ASM271v1 | train_val |
| Aspergillus costaricensis | Aspergillus_costaricensis | GCF_003184835.1_Aspcos1 | test |
| Aspergillus eucalypticola | Aspergillus_eucalypticola | GCF_003184535.1_Aspeuc1 | test |
| Aspergillus fijiensis | Aspergillus_fijiensis | GCF_003184825.1_Aspfij1 | test |
| Aspergillus fischeri | Aspergillus_fischeri | GCF_000149645.2_ASM14964v3 | test |
| Aspergillus flavus | Aspergillus_flavus | GCF_014117465.1_ASM1411746v1 | train_val |
| Aspergillus fumigatus | Aspergillus_fumigatus | GCF_000002655.1_ASM265v1 | train_val |
| Aspergillus glaucus | Aspergillus_glaucus | GCF_001890805.1_Aspgl1 | test |
| Aspergillus heteromorphus | Aspergillus_heteromorphus | GCF_003184545.1_Asphet1 | train_val |
| Aspergillus homomorphus | Aspergillus_homomorphus | GCF_003184865.1_Asphom1 | train_val |
| Aspergillus ibericus | Aspergillus_ibericus | GCF_003184845.1_Aspibe1 | train_val |
| Aspergillus japonicus | Aspergillus_japonicus | GCF_003184785.1_Aspjap1 | train_val |
| Aspergillus lentulus | Aspergillus_lentulus | GCF_010724455.1_ASM1072445v1 | test |
| Aspergillus luchuensis | Aspergillus_luchuensis | GCF_016861625.1_AkawachiiIFO4308_assembly01 | train_val |
| Aspergillus melleus | Aspergillus_melleus | GCF_016097325.1_ASM1609732v1 | train_val |
| Aspergillus mulundensis | Aspergillus_mulundensis | GCF_003369625.1_ASM336962v1 | test |
| Aspergillus neoniger | Aspergillus_neoniger | GCF_003184625.1_Aspneo1 | test |
| Aspergillus nidulans | Aspergillus_nidulans | GCF_000149205.2_ASM14920v2 | train_val |
| Aspergillus niger | Aspergillus_niger | GCF_000002855.3_ASM285v2 | test |
| Aspergillus nomiae | Aspergillus_nomiae | GCF_001204775.2_ASM120477v2 | train_val |
| Aspergillus novofumigatus | Aspergillus_novofumigatus | GCF_002847465.1_Aspnov1 | train_val |
| Aspergillus ochraceoroseus | Aspergillus_ochraceoroseus | GCF_002846915.1_Aspergillus_ochraceoroseus_IBT_... | test |
| Aspergillus oryzae | Aspergillus_oryzae | GCF_000184455.2_ASM18445v3 | train_val |
| Aspergillus piperis | Aspergillus_piperis | GCF_003184755.1_Asppip1 | test |
| Aspergillus pseudonomiae | Aspergillus_pseudonomiae | GCF_009193645.1_Asppsen1 | train_val |
| Aspergillus pseudotamarii | Aspergillus_pseudotamarii | GCF_009193445.1_Asppset1 | train_val |
| Aspergillus pseudoviridinutans | Aspergillus_pseudoviridinutans | GCF_018340605.1_Asppvi_assembly01 | train_val |
| Aspergillus puulaauensis | Aspergillus_puulaauensis | GCF_016861865.1_ApuulaauensisMK2_assembly01 | test |
| Aspergillus ruber | Aspergillus_ruber | GCF_000600275.1_Eurhe1 | train_val |
| Aspergillus saccharolyticus | Aspergillus_saccharolyticus | GCF_003184585.1_Aspsac1 | test |
| Aspergillus sclerotioniger | Aspergillus_sclerotioniger | GCF_003184525.1_Aspscl1 | train_val |
| Aspergillus steynii | Aspergillus_steynii | GCF_002849105.1_Aspste1 | train_val |
| Aspergillus sydowii | Aspergillus_sydowii | GCF_001890705.1_Aspsy1 | test |
| Aspergillus tanneri | Aspergillus_tanneri | GCF_003426965.1_ASM342696v1 | train_val |
| Aspergillus terreus | Aspergillus_terreus | GCF_000149615.1_ASM14961v1 | test (used) |
| Aspergillus thermomutatus | Aspergillus_thermomutatus | GCF_002237265.1_ASM223726v2 | train_val |
| Aspergillus tubingensis | Aspergillus_tubingensis | GCF_013340325.1_ASM1334032v1 | train_val |
| Aspergillus udagawae | Aspergillus_udagawae | GCF_001078395.1_Aud_assembly02 | train_val |
| Aspergillus uvarum | Aspergillus_uvarum | GCF_003184745.1_Aspuva1 | train_val |
| Aspergillus vadensis | Aspergillus_vadensis | GCF_003184925.1_Aspvad1 | train_val |
| Aspergillus versicolor | Aspergillus_versicolor | GCF_001890125.1_Aspve1 | train_val |
| Aspergillus viridinutans | Aspergillus_viridinutans | GCF_018404265.1_Aspvir_assembly01 | train_val |
| Aspergillus welwitschiae | Aspergillus_welwitschiae | GCF_003344945.1_Aspwel1 | train_val |
| Aspergillus wentii | Aspergillus_wentii | GCF_001890725.1_Aspwe1 | train_val |
| Aureobasidium melanogenum | Aureobasidium_melanogenum | GCF_000721775.1_Aureobasidium_pullulans_var._me... | test |
| Aureobasidium namibiae | Aureobasidium_namibiae | GCF_000721765.1_Aureobasidium_pullulans_var._na... | train_val |
| Aureobasidium pullulans | Aureobasidium_pullulans | GCF_000721785.1_Aureobasidium_pullulans_var._pu... | train_val |
| Aureobasidium subglaciale | Aureobasidium_subglaciale | GCF_000721755.1_Aureobasidium_pullulans_var._su... | train_val |
| Babjeviella inositovora | Babjeviella_inositovora | GCF_001661335.1_Babin1 | train_val |
| Bacidia gigantensis | Bacidia_gigantensis | GCF_019456465.1_ASM1945646v1 | train_val |
| Batrachochytrium dendrobatidis | Batrachochytrium_dendrobatidis | GCF_000203795.1_v1.0 | train_val |
| Baudoinia panamericana | Baudoinia_panamericana | GCF_000338955.1_Bauco1 | test |
| Beauveria bassiana | Beauveria_bassiana | GCF_000280675.1_ASM28067v1 | train_val |
| Bipolaris maydis | Bipolaris_maydis | GCF_000354255.1_CocheC4_1 | train_val |
| Bipolaris oryzae | Bipolaris_oryzae | GCF_000523455.1_Cochliobolus_miyabeanus_v1.0 | train_val |
| Bipolaris sorokiniana | Bipolaris_sorokiniana | GCF_000338995.1_Cocsa1 | train_val |
| Bipolaris victoriae | Bipolaris_victoriae | GCF_000527765.1_Cochliobolus_victoriae_v1.0 | train_val |

| scientific | ID | version | split |
|---|---|---|---|
| Bipolaris zeicola | Bipolaris_zeicola | GCF_000523435.1_Cochliobolus_carbonum_v1.0 | test |
| Blastomyces dermatitidis | Blastomyces_dermatitidis | GCF_000003525.1_BD_ER3_V1 | train_val |
| Blastomyces gilchristii | Blastomyces_gilchristii | GCF_000003855.2_BD_SLH14081_V1 | train_val |
| Boeremia exigua | Boeremia_exigua | GCF_020726555.1_Boeex1 | train_val |
| Botrytis byssoidea | Botrytis_byssoidea | GCF_014898295.1_ASM1489829v1 | test |
| Botrytis cinerea | Botrytis_cinerea | GCF_000143535.2_ASM14353v4 | train_val |
| Botrytis deweyae | Botrytis_deweyae | GCF_014898535.1_ASM1489853v1 | test |
| Botrytis fragariae | Botrytis_fragariae | GCF_013461495.1_Bfra_R1V1 | train_val |
| Botrytis porri | Botrytis_porri | GCF_014898465.1_ASM1489846v1 | train_val |
| Botrytis sinoallii | Botrytis_sinoallii | GCF_014898435.1_ASM1489843v1 | test |
| Brettanomyces bruxellensis | Brettanomyces_bruxellensis | GCF_011074885.1_ASM1107488v2 | test |
| Brettanomyces nanus | Brettanomyces_nanus | GCF_011074865.1_ASM1107486v2 | train_val |
| Candida albicans | Candida_albicans | GCF_000182965.3_ASM18296v3 | train_val |
| Candida auris | _Candida_auris | GCF_002775015.1_Cand_auris_B11221_V1 | train_val |
| Candida dubliniensis | Candida_dubliniensis | GCF_000026945.1_ASM2694v1 | train_val |
| Candida duobushaemulonis | _Candida_duobushaemulonis | GCF_002926085.2_CanDuoHae_v1.0 | test |
| Candida glabrata | _Candida_glabrata | GCF_000002545.3_ASM254v2 | test |
| Candida haemuloni | _Candida_haemuloni | GCF_002926055.2_CanHae_1.0 | train_val |
| Candida orthopsilosis | Candida_orthopsilosis | GCF_000315875.1_ASM31587v1 | train_val |
| Candida parapsilosis | Candida_parapsilosis | GCF_000182765.1_ASM18276v2 | train_val |
| Candida pseudohaemulonii | _Candida_pseudohaemulonii | GCF_003013735.1_Cand_pseudohaemulonii_B12108 | test |
| Candida tropicalis | Candida_tropicalis | GCF_000006335.3_ASM633v3 | train_val |
| Cantharellus anzutake | Cantharellus_anzutake | GCF_015039405.1_Cananz1 | train_val |
| Capronia coronata | Capronia_coronata | GCF_000585585.1_Capr_coro_CBS_617_96_V1 | train_val |
| Capronia epimyces | Capronia_epimyces | GCF_000585565.1_Capr_epim_CBS_606_96_V1 | train_val |
| Ceraceosorus guamensis | Ceraceosorus_guamensis | GCF_003144195.1_Cersp1 | train_val |
| Cercospora beticola | Cercospora_beticola | GCF_002742065.1_CB0940_V2 | test |
| Cercospora kikuchii | Cercospora_kikuchii | GCF_019650295.1_Ck_assembly01 | test |
| Chaetomium globosum | Chaetomium_globosum | GCF_000143365.1_ASM14336v1 | train_val |
| Chaetomium thermophilum | Chaetomium_thermophilum | GCF_000221225.1_CTHT_3.0 | train_val |
| Cladophialophora bantiana | Cladophialophora_bantiana | GCF_000835475.1_Clad_bant_CBS_173_52_V1 | test |
| Cladophialophora carrionii | Cladophialophora_carrionii | GCF_000365165.1_Clad_carr_CBS_160_54_V1 | train_val |
| Cladophialophora immunda | Cladophialophora_immunda | GCF_000835495.1_Clad_immu_CBS83496_V1 | test |
| Cladophialophora psammophila | Cladophialophora_psammophila | GCF_000585535.1_Clad_psam_CBS_110553_V1 | test |
| Cladophialophora yegresii | Cladophialophora_yegresii | GCF_000585515.1_Clad_yegr_CBS_114405_V1 | test |
| Clavispora lusitaniae | Clavispora_lusitaniae | GCF_000003835.1_ASM383v1 | train_val |
| Coccidioides immitis | Coccidioides_immitis | GCF_000149335.2_ASM14933v2 | test (used) |
| Coccidioides posadasii | Coccidioides_posadasii | GCF_000151335.2_JCVI-cpa1-1.0 | train_val |
| Colletotrichum aenigma | Colletotrichum_aenigma | GCF_013390185.1_ASM1339018v1 | train_val |
| Colletotrichum fructicola | Colletotrichum_fructicola | GCF_009771025.1_ASM977102v1 | train_val |
| Colletotrichum gloeosporioides | Colletotrichum_gloeosporioides | GCF_011800055.1_NFU_CgLc1_1.0 | test |
| Colletotrichum graminicola | Colletotrichum_graminicola | GCF_000149035.1_C_graminicola_M1_001_V1 | train_val |
| Colletotrichum higginsianum | Colletotrichum_higginsianum | GCF_001672515.1_ASM167251v1 | train_val |
| Colletotrichum karsti | Colletotrichum_karsti | GCF_011947395.1_ASM1194739v2 | train_val |
| Colletotrichum orchidophilum | Colletotrichum_orchidophilum | GCF_001831195.1_CORC01 | test |
| Colletotrichum scovillei | Colletotrichum_scovillei | GCF_011075155.1_ASM1107515v1 | test |
| Colletotrichum siamense | Colletotrichum_siamense | GCF_013390195.1_ASM1339019v1 | train_val |
| Colletotrichum truncatum | Colletotrichum_truncatum | GCF_014235925.1_CTRU02 | test |
| Coniophora puteana | Coniophora_puteana | GCF_000271625.1_Conpu1 | train_val |
| Coniosporium apollinis | Coniosporium_apollinis | GCF_000281105.1_Coni_apol_CBS100218_V1 | train_val |
| Coprinopsis cinerea | Coprinopsis_cinerea | GCF_000182895.1_CC3 | train_val |
| Cordyceps fumosorosea | Cordyceps_fumosorosea | GCF_001636725.1_ISF_1.0 | test |
| Cordyceps militaris | Cordyceps_militaris | GCF_000225605.1_CmilitarisCM01_v01 | train_val |
| Cryphonectria parasitica | Cryphonectria_parasitica | GCF_011745365.1_Crypa2 | test |
| Cryptococcus amylolentus | Cryptococcus_amylolentus | GCF_001720205.1_Cryp_amyl_CBS6039_V3 | train_val |
| Cryptococcus gattii VGI | Cryptococcus_gattii_VGI | GCF_000185945.1_ASM18594v1 | test |
| Cryptococcus neoformans | Cryptococcus_neoformans | GCF_000091045.1_ASM9104v1 | train_val |
| Cryptococcus wingfieldii | Cryptococcus_wingfieldii | GCF_001720155.1_Tsuc_wing_CBS7118_V1 | test |
| Cucurbitaria berberidis | Cucurbitaria_berberidis | GCF_010015615.1_Cucbe1 | train_val |
| Cutaneotrichosporon oleaginosum | Cutaneotrichosporon_oleaginosum | GCF_001027345.1_Triol1 | train_val |
| Cyberlindnera jadinii | Cyberlindnera_jadinii | GCF_001661405.1_Cybja1 | train_val |
| Cyphellophora europaea | Cyphellophora_europaea | GCF_000365145.1_Phia_euro_CBS_101466_V1 | train_val |
| Dacryopinax primogenitus | Dacryopinax_primogenitus | GCF_000292625.1_Dacryopinax_sp._DJM_731_SSP1_v1.0 | train_val |
| Daldinia childiae | Daldinia_childiae | GCF_008694065.1_Dalch_JS-1345 | test |
| Debaryomyces fabryi | Debaryomyces_fabryi | GCF_001447935.2_debFab1.1 | train_val |
| Debaryomyces hansenii | Debaryomyces_hansenii | GCF_000006445.2_ASM644v2 | test (used) |
| Diaporthe batatas | Diaporthe_batatas | GCF_019321695.1_ASM1932169v1 | train_val |
| Diaporthe citri | Diaporthe_citri | GCF_014595645.1_ASM1459564v1 | train_val |
| Dichomitus squalens | Dichomitus_squalens | GCF_000275845.1_Dichomitus_squalens_v1.0 | train_val |
| Didymella exigua | Didymella_exigua | GCF_010094145.1_Didex1 | test |
| Diplodia corticola | Diplodia_corticola | GCF_001883845.1_ASM188384v1 | train_val |
| Dissoconium aciculare | Dissoconium_aciculare | GCF_010015565.1_Disac1 | test |
| Diutina rugosa | Diutina_rugosa | GCF_008704595.1_ASM870459v1 | train_val |
| Dothidotthia symphoricarpi | Dothidotthia_symphoricarpi | GCF_010015815.1_Dotsy1 | train_val |
| Drechmeria coniospora | Drechmeria_coniospora | GCF_001625195.1_ASM162519v1 | train_val |
| Drepanopeziza brunnea | Drepanopeziza_brunnea | GCF_000298775.1_ASM29877v1 | test |
| Emericellopsis atlantica | Emericellopsis_atlantica | GCF_019669845.1_AcreTS7_1 | train_val |
| Encephalitozoon cuniculi | Encephalitozoon_cuniculi | GCF_000091225.1_ASM9122v1 | train_val |
| Encephalitozoon hellem | Encephalitozoon_hellem | GCF_000277815.2_ASM27781v3 | train_val |
| Encephalitozoon intestinalis | Encephalitozoon_intestinalis | GCF_000146465.1_ASM14646v1 | test |

| scientific | ID | version | split |
|---|---|---|---|
| Encephalitozoon romaleae | Encephalitozoon_romaleae | GCF_000280035.1_ASM28003v2 | train_val |
| Endocarpon pusillum | Endocarpon_pusillum | GCF_000464535.1_EPUS | train_val |
| Eremomyces bilateralis | Eremomyces_bilateralis | GCF_010015585.1_Erebi1 | train_val |
| Eremothecium cymbalariae | Eremothecium_cymbalariae | GCF_000235365.1_ASM23536v1 | test |
| Eremothecium gossypii | Eremothecium_gossypii | GCF_000091025.4_ASM9102v4 | train_val |
| Eremothecium sinecaudum | Eremothecium_sinecaudum | GCF_001548555.1_ASM154855v1 | train_val |
| Exophiala aquamarina | Exophiala_aquamarina | GCF_000709125.1_Exop_aqua_CBS_119918_V1 | train_val |
| Exophiala dermatitidis | Exophiala_dermatitidis | GCF_000230625.1_Exop_derm_V1 | test |
| Exophiala mesophila | Exophiala_mesophila | GCF_000836275.1_Exop_meso_CBS40295_V1 | test |
| Exophiala oligosperma | Exophiala_oligosperma | GCF_000835515.1_Exop_olig_CBS72588_V1 | train_val |
| Exophiala spinifera | Exophiala_spinifera | GCF_000836115.1_Exop_spin_CBS89968_V1 | train_val |
| Exophiala xenobiotica | Exophiala_xenobiotica | GCF_000835505.1_Exop_xeno_CBS118157_V1 | train_val |
| Exserohilum turcicum | Exserohilum_turcicum | GCF_000359705.1_Setospaeria_trucica_Et28A_v1.0 | train_val |
| Fibroporia radiculosa | Fibroporia_radiculosa | GCF_000313525.1_ASM31352v1 | train_val |
| Filobasidium floriforme | Filobasidium_floriforme | GCF_021052385.1_Filflo1 | train_val |
| Fomitiporia mediterranea | Fomitiporia_mediterranea | GCF_000271605.1_Fomme1 | train_val |
| Fonsecaea erecta | Fonsecaea_erecta | GCF_001651985.1_ASM165198v1 | train_val |
| Fonsecaea monophora | Fonsecaea_monophora | GCF_001642475.1_ASM164247v1 | train_val |
| Fonsecaea multimorphosa | Fonsecaea_multimorphosa | GCF_000836435.1_Fons_mult_CBS_102226_V1 | test |
| Fonsecaea nubica | Fonsecaea_nubica | GCF_001646965.1_ASM164696v1 | train_val |
| Fonsecaea pedrosoi | Fonsecaea_pedrosoi | GCF_000835455.1_Fons_pedr_CBS_271_37_V1 | train_val |
| Fusarium coffeatum | Fusarium_coffeatum | GCF_003316985.1_ASM331698v1 | train_val |
| Fusarium flagelliforme | Fusarium_flagelliforme | GCF_020744385.1_Fuseq1 | train_val |
| Fusarium fujikuroi | Fusarium_fujikuroi | GCF_900079805.1_Fusarium_fujikuroi_IMI58289_V2 | train_val |
| Fusarium graminearum | Fusarium_graminearum | GCF_000240135.3_ASM24013v3 | test (used) |
| Fusarium mangiferae | Fusarium_mangiferae | GCF_900044065.1_Genome_assembly_version_1 | train_val |
| Fusarium musae | Fusarium_musae | GCF_019915245.1_ASM1991524v1 | test |
| Fusarium odoratissimum | Fusarium_odoratissimum | GCF_000260195.1_FO_II5_V1 | train_val |
| Fusarium oxysporum | Fusarium_oxysporum | GCF_000271745.1_FO_FOSC_3_a_V1 | train_val |
| Fusarium poae | Fusarium_poae | GCF_019609905.1_ASM1960990v1 | test |
| Fusarium proliferatum | Fusarium_proliferatum | GCF_900067095.1_F._proliferatum_ET1_version_1 | train_val |
| Fusarium pseudograminearum | Fusarium_pseudograminearum | GCF_000303195.2_FP7 | train_val |
| Fusarium redolens | Fusarium_redolens | GCF_020744475.1_Fusre1 | train_val |
| Fusarium solani | Fusarium_solani | GCF_020744495.1_Fusso1 | test |
| Fusarium subglutinans | Fusarium_subglutinans | GCF_013396075.1_ASM1339607v1 | test |
| Fusarium tjaetaba | Fusarium_tjaetaba | GCF_013396195.1_ASM1339619v1 | test |
| Fusarium vanettenii | Fusarium_vanettenii | GCF_000151355.1_v2.0 | test |
| Fusarium venenatum | Fusarium_venenatum | GCF_900007375.1_ASM90000737v1 | train_val |
| Fusarium verticillioides | Fusarium_verticillioides | GCF_000149555.1_ASM14955v1 | test |
| Gaeumannomyces tritici | Gaeumannomyces_tritici | GCF_000145635.1_Gae_graminis_V2 | test |
| Geosmithia morbida | Geosmithia_morbida | GCF_012550715.1_ASM1255071v1 | train_val |
| Glarea lozoyensis | Glarea_lozoyensis | GCF_000409485.1_GLAREA | test |
| Gloeophyllum trabeum | Gloeophyllum_trabeum | GCF_000344685.1_Glotr1_1 | test |
| Grosmannia clavigera | Grosmannia_clavigera | GCF_000143105.1_Sanger-454-IlluminaPA_2.0 | train_val |
| Guyanagaster necrorhizus | Guyanagaster_necrorhizus | GCF_019112545.1_Guyne1 | train_val |
| Heterobasidion irregulare | Heterobasidion_irregulare | GCF_000320585.1_Heterobasidion_irregulare_v2.0 | train_val |
| Hirsutella rhossiliensis | Hirsutella_rhossiliensis | GCF_020360975.1_ASM2036097v1 | train_val |
| Histoplasma capsulatum | Histoplasma_capsulatum | GCF_000150115.1_ASM15011v1 | train_val |
| Histoplasma mississippiense nom inval | Histoplasma_mississippiense_nom._inval._ | GCF_000149585.1_ASM14958v1 | train_val |
| Hyaloscypha bicolor | Hyaloscypha_bicolor | GCF_002865645.1_Melbi2 | train_val |
| Hyphopichia burtonii | Hyphopichia_burtonii | GCF_001661395.1_Hypbu1 | train_val |
| Ilyonectria robusta | Ilyonectria_robusta | GCF_021365365.1_Ilyrob1 | test |
| Jaminaea rosea | Jaminaea_rosea | GCF_003144245.1_Jamsp1 | train_val |
| Kalmanozyma brasiliensis | Kalmanozyma_brasiliensis | GCF_000497045.1_PSEUBRA1 | test |
| Kazachstania africana | Kazachstania_africana | GCF_000304475.1_Ka_CBS2517 | test |
| Kazachstania barnettii | Kazachstania_barnettii | GCF_903064755.1_KABA2 | test |
| Kazachstania naganishii | Kazachstania_naganishii | GCF_000348985.1_ASM34898v1 | test |
| Kluyveromyces lactis | Kluyveromyces_lactis | GCF_000002515.2_ASM251v1 | train_val |
| Kluyveromyces marxianus | Kluyveromyces_marxianus | GCF_001417885.1_Kmar_1.0 | test |
| Kockovaella imperatae | Kockovaella_imperatae | GCF_002102565.1_Kocim1 | train_val |
| Komagataella phaffii | Komagataella_phaffii | GCF_000027005.1_ASM2700v1 | train_val |
| Kuraishia capsulata | Kuraishia_capsulata | GCF_000576695.1_AUH_PRJEB4427_v1 | train_val |
| Kwoniella bestiolae | Kwoniella_bestiolae | GCF_000512585.1_Cryp_best_CBS10118_V1 | train_val |
| Kwoniella dejecticola | Kwoniella_dejecticola | GCF_000512565.1_Cryp_deje_CBS10117_V1 | train_val |
| Kwoniella mangrovensis | Kwoniella_mangrovensis | GCF_000507465.1_Kwon_mang_CBS8507_V2 | train_val |
| Kwoniella pini | Kwoniella_pini | GCF_000512605.1_Cryp_pinu_CBS10737_V1 | train_val |
| Kwoniella shandongensis | Kwoniella_shandongensis | GCF_008629635.1_Kwon_shan_CBS_12478_V1 | train_val |
| Laccaria bicolor | Laccaria_bicolor | GCF_000143565.1_V1.0 | test (used) |
| Lachancea lanzarotensis | Lachancea_lanzarotensis | GCF_000938715.1_LALA0 | test |
| Lachancea thermotolerans | Lachancea_thermotolerans | GCF_000142805.1_ASM14280v1 | train_val |
| Lachnellula hyalina | Lachnellula_hyalina | GCF_007821495.1_CFIA_Lhya_EG2017 | test |
| Laetiporus sulphureus | Laetiporus_sulphureus | GCF_001632365.1_Laesu1 | train_val |
| Lasiodiplodia theobromae | Lasiodiplodia_theobromae | GCF_012971845.1_ASM1297184v1 | train_val |
| Lentinula edodes | Lentinula_edodes | GCF_021015755.1_Lenedo1 | train_val |
| Leptosphaeria maculans | Leptosphaeria_maculans | GCF_000230375.1_ASM23037v1 | train_val |
| Letharia columbiana | Letharia_columbiana | GCF_014066305.1_Lecol_v1.0 | train_val |
| Letharia lupina | Letharia_lupina | GCF_014066315.1_Lelup_v1.1 | test |
| Linderina pennispora | Linderina_pennispora | GCF_002104995.1_Linpe1 | test |
| Lindgomyces ingoldianus | Lindgomyces_ingoldianus | GCF_010093535.1_Linin1 | train_val |
| Lobosporangium transversale | Lobosporangium_transversale | GCF_002105155.1_Lobtra1 | test |

| scientific | ID | version | split |
|---|---|---|---|
| Lodderomyces elongisporus | Lodderomyces_elongisporus | GCF_000149685.1_ASM14968v1 | train_val |
| Macroventuria anomochaeta | Macroventuria_anomochaeta | GCF_010093625.1_Macan1 | train_val |
| Malassezia globosa | Malassezia_globosa | GCF_000181695.1_ASM18169v1 | train_val |
| Malassezia pachydermatis | Malassezia_pachydermatis | GCF_001278385.1_MalaPachy | train_val |
| Malassezia restricta | Malassezia_restricta | GCF_003290485.1_ASM329048v1 | test |
| Malassezia sympodialis | Malassezia_sympodialis | GCF_000349305.1_ASM34930v2 | train_val |
| Marasmius oreades | Marasmius_oreades | GCF_018924745.1_UU_Maror_2 | test |
| Meira miltonrushii | Meira_miltonrushii | GCF_003144205.1_Meimi1 | train_val |
| Melampsora larici-populina | Melampsora_larici-populina | GCF_000204055.1_v1.0 | train_val |
| Metarhizium acridum | Metarhizium_acridum | GCF_000187405.1_MetAcr_May2010 | test |
| Metarhizium album | Metarhizium_album | GCF_000804445.1_MAM_1.0_for_version_1_of_the_Me... | train_val |
| Metarhizium brunneum | Metarhizium_brunneum | GCF_000814965.1_MBR_1.0 | train_val |
| Metarhizium robertsii | Metarhizium_robertsii | GCF_000187425.2_MAA_2.0 | test |
| Metschnikowia bicuspidata | Metschnikowia_bicuspidata | GCF_001664035.1_Metbi1 | train_val |
| Meyerozyma guilliermondii | Meyerozyma_guilliermondii | GCF_000149425.1_ASM14942v1 | train_val |
| Microdochium trichocladiopsis | Microdochium_trichocladiopsis | GCF_020744255.1_Mictri1 | test |
| Microsporum canis | Microsporum_canis | GCF_000151145.1_ASM15114v1 | train_val |
| Mitosporidium daphniae | Mitosporidium_daphniae | GCF_000760515.2_UGP1.1 | test |
| Mixia osmundae | Mixia_osmundae | GCF_000708205.1_Mixia_osmundae_v1.0 | train_val |
| Moesziomyces antarcticus | Moesziomyces_antarcticus | GCF_000747765.1_ASM74776v1 | train_val |
| Mollisia scopiformis | Mollisia_scopiformis | GCF_001500285.1_Phisc1 | train_val |
| Morchella importuna | Morchella_importuna | GCF_003444635.1_ASM344463v2 | train_val |
| Morchella sextelata | Morchella_sextelata | GCF_020137385.1_ASM2013738v1 | train_val |
| Mycena indigotica | Mycena_indigotica | GCF_014461135.1_ASM1446113v1 | train_val |
| Mytilinidion resinicola | Mytilinidion_resinicola | GCF_010093595.1_Mytre1 | train_val |
| Nannizzia gypsea | Nannizzia_gypsea | GCF_000150975.2_MS_CBS118893 | test |
| Naumovozyma castellii | Naumovozyma_castellii | GCF_000237345.1_ASM23734v1 | train_val |
| Naumovozyma dairenensis | Naumovozyma_dairenensis | GCF_000227115.2_ASM22711v2 | train_val |
| Nematocida parisii | Nematocida_parisii | GCF_000250985.1_Nema_parisii_ERTm1_V3 | train_val |
| Neohortaea acidophila | Neohortaea_acidophila | GCF_010093505.1_Horac1 | train_val |
| Neurospora crassa | Neurospora_crassa | GCF_000182925.2_NC12 | test (used) |
| Neurospora tetrasperma | Neurospora_tetrasperma | GCF_000213175.1_v2.0 | train_val |
| Nosema ceranae | Nosema_ceranae | GCF_000988165.1_ASM98816v1 | train_val |
| Ogataea angusta | Ogataea_angusta | GCF_019207475.1_ASM1920747v1 | test |
| Ogataea haglerorum | Ogataea_haglerorum | GCF_019207285.1_ASM1920728v1 | train_val |
| Ogataea parapolymorpha | Ogataea_parapolymorpha | GCF_000187245.1_Hansenula_2 | train_val |
| Ogataea philodendri | Ogataea_philodendri | GCF_020536065.1_ASM2053606v1 | test |
| Ogataea polymorpha | Ogataea_polymorpha | GCF_001664045.1_Hanpo2 | train_val |
| Orbilia oligospora | Orbilia_oligospora | GCF_000225545.1_AOL24927_1.0 | test |
| Ordospora colligata | Ordospora_colligata | GCF_000803265.1_ASM80326v1 | test |
| Paecilomyces variotii | Paecilomyces_variotii | GCF_004022145.1_Paevar1 | test |
| Paracoccidioides brasiliensis | Paracoccidioides_brasiliensis | GCF_000150735.1_Paracocci_br_Pb18_V2 | test |
| Paracoccidioides lutzii | Paracoccidioides_lutzii | GCF_000150705.2_Paracocci_br_Pb01_V2 | test |
| Paraphaeosphaeria sporulosa | Paraphaeosphaeria_sporulosa | GCF_001642045.1_Parsp1 | train_val |
| Parastagonospora nodorum | Parastagonospora_nodorum | GCF_000146915.1_ASM14691v2 | train_val |
| Penicilliopsis zonata | Penicilliopsis_zonata | GCF_001890105.1_Aspzo1 | train_val |
| Penicillium arizonense | Penicillium_arizonense | GCF_001773325.1_ASM177332v1 | test |
| Penicillium digitatum | Penicillium_digitatum | GCF_000315645.1_PdigPd1_v1 | train_val |
| Penicillium expansum | Penicillium_expansum | GCF_000769745.1_ASM76974v1 | test |
| Penicillium griseofulvum | Penicillium_griseofulvum | GCF_001561935.1_ASM156193v1 | train_val |
| Penicillium roqueforti | Penicillium_roqueforti | GCF_015533775.1_ASM1553377v1 | train_val |
| Penicillium rubens | Penicillium_rubens | GCF_000226395.1_PenChr_Nov2007 | train_val |
| Penicillium solitum | Penicillium_solitum | GCF_002072235.1_ASM207223v1 | train_val |
| Pestalotiopsis fici | Pestalotiopsis_fici | GCF_000516985.1_PFICI | train_val |
| Phaeoacremonium minimum | Phaeoacremonium_minimum | GCF_000392275.1_UCRPA7V03 | train_val |
| Phanerochaete carnosa | Phanerochaete_carnosa | GCF_000300595.1_Phanerochaete_carnosa_HHB-10118... | test |
| Phialophora attinorum | Phialophora_attinorum | GCF_001299255.1_ASM129925v1 | test |
| Phycomyces blakesleeanus | Phycomyces_blakesleeanus | GCF_001638985.1_Phybl2 | train_val |
| Pichia kudriavzevii | Pichia_kudriavzevii | GCF_003054445.1_ASM305444v1 | train_val |
| Pichia membranifaciens | Pichia_membranifaciens | GCF_001661235.1_Picme2 | train_val |
| Pleurotus ostreatus | Pleurotus_ostreatus | GCF_014466165.1_ASM1446616v1 | train_val |
| Pneumocystis carinii | Pneumocystis_carinii | GCF_001477545.1_Pneu_cari_B80_V3 | test |
| Pneumocystis jirovecii | Pneumocystis_jirovecii | GCF_001477535.1_Pneu_jiro_RU7_V2 | train_val |
| Pneumocystis murina | Pneumocystis_murina | GCF_000349005.2_Pneumo_murina_B123_V4 | train_val |
| Pochonia chlamydosporia | Pochonia_chlamydosporia | GCF_001653235.2_ASM165323v2 | train_val |
| Podospora anserina | Podospora_anserina | GCF_000226545.1_ASM22654v1 | train_val |
| Postia placenta | Postia_placenta | GCF_002117355.1_PosplRSB12_1 | train_val |
| Protomyces lactucae-debilis | Protomyces_lactucae-debilis | GCF_002105105.1_Prola1 | train_val |
| Pseudocercospora fijiensis | Pseudocercospora_fijiensis | GCF_000340215.1_Mycfi2 | test |
| Pseudogymnoascus destructans | Pseudogymnoascus_destructans | GCF_001641265.1_ASM164126v1 | train_val |
| Pseudogymnoascus verrucosus | Pseudogymnoascus_verrucosus | GCF_001662655.1_ASM166265v1 | train_val |
| Pseudomassariella vexata | Pseudomassariella_vexata | GCF_002105095.1_Pseve2 | train_val |
| Pseudomicrostroma glucosiphilum | Pseudomicrostroma_glucosiphilum | GCF_003144135.1_Rhodsp1 | train_val |
| Pseudovirgaria hyperparasitica | Pseudovirgaria_hyperparasitica | GCF_010093815.1_Psehy1 | train_val |
| Pseudozyma flocculosa | Pseudozyma_flocculosa | GCF_000417875.1_Pflocc_1.0 | train_val |
| Pseudozyma hubeiensis | Pseudozyma_hubeiensis | GCF_000403515.1_ASM40351v1 | train_val |
| Puccinia graminis | Puccinia_graminis | GCF_000149925.1_ASM14992v1 | test |
| Punctularia strigosozonata | Punctularia_strigosozonata | GCF_000264995.1_Punctularia_strigosozonata_v1.0 | train_val |
| Purpureocillium lilacinum | Purpureocillium_lilacinum | GCF_001653265.1_ASM165326v1 | test |
| Pyrenophora tritici-repentis | Pyrenophora_tritici-repentis | GCF_000149985.1_ASM14998v1 | train_val |

| scientific | ID | version | split |
|---|---|---|---|
| Pyricularia grisea | Pyricularia_grisea | GCF_004355905.1_ASM435590v1 | train_val |
| Pyricularia oryzae | Pyricularia_oryzae | GCF_000002495.2_MG8 | train_val |
| Pyricularia pennisetigena | Pyricularia_pennisetigena | GCF_004337985.1_ASM433798v1 | train_val |
| Ramularia collo-cygni | Ramularia_collo-cygni | GCF_900074925.1_version_1 | train_val |
| Rasamsonia emersonii | Rasamsonia_emersonii | GCF_000968595.1_ASM96859v1 | test |
| Rhinocladiella mackenziei | Rhinocladiella_mackenziei | GCF_000835555.1_Rhin_mack_CBS_650_93_V1 | train_val |
| Rhizoctonia solani | Rhizoctonia_solani | GCF_016906535.1_ASM1690653v1 | test |
| Rhizophagus irregularis | Rhizophagus_irregularis | GCF_000439145.1_ASM43914v3 | train_val |
| Rhizopus microsporus | Rhizopus_microsporus | GCF_002708625.1_Rhimi1_1 | test |
| Rhodotorula graminis | Rhodotorula_graminis | GCF_001329695.1_Rhoba1_1 | test |
| Rhodotorula toruloides | Rhodotorula_toruloides | GCF_000320785.1_RHOziaDV1.0 | train_val |
| Saccharomyces cerevisiae | Saccharomyces_cerevisiae | GCF_000146045.2_R64 | train_val |
| Saccharomyces eubayanus | Saccharomyces_eubayanus | GCF_001298625.1_SEUB3.0 | test |
| Saccharomyces paradoxus | Saccharomyces_paradoxus | GCF_002079055.1_ASM207905v1 | train_val |
| Saccharomycodes ludwigii | Saccharomycodes_ludwigii | GCF_020623625.1_UHD_SCDLUD_16 | train_val |
| Saitoella complicata | Saitoella_complicata | GCF_001661265.1_Saico1 | train_val |
| Saprochaete ingens | Saprochaete_ingens | GCF_902498895.1_sapIngB | train_val |
| Scedosporium apiospermum | Scedosporium_apiospermum | GCF_000732125.1_ScApio1.0 | train_val |
| Scheffersomyces spartinae | Scheffersomyces_spartinae | GCF_019049425.1_ASM1904942v1 | train_val |
| Scheffersomyces stipitis | Scheffersomyces_stipitis | GCF_000209165.1_ASM20916v1 | train_val |
| Schizophyllum commune | Schizophyllum_commune | GCF_000143185.1_v1.0 | train_val |
| Schizosaccharomyces cryophilus | Schizosaccharomyces_cryophilus | GCF_000004155.1_SCY4 | train_val |
| Schizosaccharomyces japonicus | Schizosaccharomyces_japonicus | GCF_000149845.2_SJ5 | train_val |
| Schizosaccharomyces octosporus | Schizosaccharomyces_octosporus | GCF_000150505.1_SO6 | test |
| Schizosaccharomyces pombe | Schizosaccharomyces_pombe | GCF_000002945.1_ASM294v2 | train_val |
| Sclerotinia sclerotiorum | Sclerotinia_sclerotiorum | GCF_000146945.2_ASM14694v2 | train_val |
| Serpula lacrymans | Serpula_lacrymans | GCF_000218685.1_v1.0 | test |
| Sodiomyces alkalinus | Sodiomyces_alkalinus | GCF_003711515.1_Sodal1 | test |
| Sordaria macrospora | Sordaria_macrospora | GCF_000182805.2_ASM18280v2 | train_val |
| Sparassis crispa | Sparassis_crispa | GCF_003851025.1_SCP_1.1 | train_val |
| Spathaspora passalidarum | Spathaspora_passalidarum | GCF_000223485.1_Spathaspora_passalidarum_v2.0 | train_val |
| Sphaerulina musiva | Sphaerulina_musiva | GCF_000320565.1_Septoria_musiva_SO2202_v1.0 | train_val |
| Spizellomyces punctatus | Spizellomyces_punctatus | GCF_000182565.1_S_punctatus_V1 | train_val |
| Sporisorium graminicola | Sporisorium_graminicola | GCF_005498985.1_PGRAM_IIB_1.0 | train_val |
| Sporothrix brasiliensis | Sporothrix_brasiliensis | GCF_000820605.1_S_brasiliensis_5110_v1 | train_val |
| Sporothrix schenckii | Sporothrix_schenckii | GCF_000961545.1_S_schenckii_v1 | train_val |
| Stereum hirsutum | Stereum_hirsutum | GCF_000264905.1_Stehi1 | train_val |
| Sugiyamaella lignohabitans | Sugiyamaella_lignohabitans | GCF_001640025.1_ASM164002v2 | train_val |
| Suhomyces tanzawaensis | Suhomyces_tanzawaensis | GCF_001661415.1_Canta1 | test |
| Suillus bovinus | Suillus_bovinus | GCF_016758785.1_Suibov1 | train_val |
| Suillus clintonianus | Suillus_clintonianus | GCF_016758775.1_Suicli1 | train_val |
| Suillus discolor | Suillus_discolor | GCF_016758755.1_Suidis1 | train_val |
| Suillus fuscotomentosus | Suillus_fuscotomentosus | GCF_016647785.1_Suifus1 | test |
| Suillus paluster | Suillus_paluster | GCF_016628075.1_Suipal1 | train_val |
| Suillus plorans | Suillus_plorans | GCF_016647745.1_Suiplo1 | train_val |
| Suillus subalutaceus | Suillus_subalutaceus | GCF_016647625.1_Suisu1 | test |
| Suillus subaureus | Suillus_subaureus | GCF_016647635.1_Suisub1 | train_val |
| Synchytrium microbalum | Synchytrium_microbalum | GCF_006535985.1_ASM653598v1 | train_val |
| Talaromyces amestolkiae | Talaromyces_amestolkiae | GCF_001896365.1_ASM189636v1 | train_val |
| Talaromyces atroroseus | Talaromyces_atroroseus | GCF_001907595.1_ASM190759v1 | train_val |
| Talaromyces marneffei | Talaromyces_marneffei | GCF_000001985.1_JCVI-PMFA1-2.0 | train_val |
| Talaromyces proteolyticus | Talaromyces_proteolyticus | GCF_021365285.1_Talpro1 | test |
| Talaromyces rugulosus | Talaromyces_rugulosus | GCF_013368755.1_ASM1336875v1 | train_val |
| Talaromyces stipitatus | Talaromyces_stipitatus | GCF_000003125.1_JCVI-TSTA1-3.0 | train_val |
| Tetrapisispora blattae | Tetrapisispora_blattae | GCF_000315915.1_ASM31591v1 | train_val |
| Tetrapisispora phaffii | Tetrapisispora_phaffii | GCF_000236905.1_ASM23690v1 | test |
| Thermothelomyces thermophilus | Thermothelomyces_thermophilus | GCF_000226095.1_ASM22609v1 | train_val |
| Thermothielavioides terrestris | Thermothielavioides_terrestris | GCF_000226115.1_ASM22611v1 | train_val |
| Thyridium curvatum | Thyridium_curvatum | GCF_004353045.1_ASM435304v1 | train_val |
| Tilletiaria anomala | Tilletiaria_anomala | GCF_000711695.1_Tilletiaria_anomala_UBC_951_v1.0 | train_val |
| Tilletiopsis washingtonensis | Tilletiopsis_washingtonensis | GCF_003144115.1_Tilwa1 | train_val |
| Torulaspora delbrueckii | Torulaspora_delbrueckii | GCF_000243375.1_ASM24337v1 | test |
| Torulaspora globosa | Torulaspora_globosa | GCF_014133895.1_ASM1413389v1 | test |
| Trametes versicolor | Trametes_versicolor | GCF_000271585.1_Trametes_versicolor_v1.0 | train_val |
| Trematosphaeria pertusa | Trematosphaeria_pertusa | GCF_010094035.1_Trepe1 | test |
| Tremella mesenterica | Tremella_mesenterica | GCF_000271645.1_Treme1 | train_val |
| Trichoderma asperellum | Trichoderma_asperellum | GCF_003025105.1_Trias_v._1.0 | test |
| Trichoderma atroviride | Trichoderma_atroviride | GCF_000171015.1_TRIAT_v2.0 | train_val |
| Trichoderma citrinoviride | Trichoderma_citrinoviride | GCF_003025115.1_Trici_v4.0 | train_val |
| Trichoderma gamsii | Trichoderma_gamsii | GCF_001481775.2_TGAM01v2 | train_val |
| Trichoderma harzianum | Trichoderma_harzianum | GCF_003025095.1_Triha_v1.0 | train_val |
| Trichoderma reesei | Trichoderma_reesei | GCF_000167675.1_v2.0 | train_val |
| Trichoderma virens | Trichoderma_virens | GCF_000170995.1_TRIVI_v2.0 | train_val |
| Trichophyton benhamiae | Trichophyton_benhamiae | GCF_000151125.1_ASM15112v2 | train_val |
| Trichophyton rubrum | Trichophyton_rubrum | GCF_000151425.1_ASM15142v1 | test |
| Trichophyton verrucosum | Trichophyton_verrucosum | GCF_000151505.1_ASM15150v1 | test |
| Trichosporon asahii | Trichosporon_asahii | GCF_000293215.1_Trichosporon_asahii_1 | test |
| Truncatella angustata | Truncatella_angustata | GCF_020726525.1_Truan1 | test |
| Tuber melanosporum | Tuber_melanosporum | GCF_000151645.1_ASM15164v1 | train_val |
| Uncinocarpus reesii | Uncinocarpus_reesii | GCF_000003515.1_ASM351v2 | train_val |

| scientific | ID | version | split |
|---|---|---|---|
| Ustilaginoidea virens | Ustilaginoidea_virens | GCF_000687475.1_ASM68747v2 | test |
| Ustilago hordei | Ustilago_hordei | GCF_900519145.1_Uho2_v1 | train_val |
| Ustilago maydis | Ustilago_maydis | GCF_000328475.2_Umaydis521_2.0 | train_val |
| Vanderwaltozyma polyspora | Vanderwaltozyma_polyspora | GCF_000150035.1_ASM15003v1 | train_val |
| Vavraia culicis | Vavraia_culicis | GCF_000192795.1_Vavr_culi_floridensis_V1 | train_val |
| Venustampulla echinocandica | Venustampulla_echinocandica | GCF_003357145.1_ASM335714v1 | train_val |
| Verruconis gallopava | Verruconis_gallopava | GCF_000836295.1_O_gall_CBS43764 | test |
| Verticillium alfalfae | Verticillium_alfalfae | GCF_000150825.1_ASM15082v1 | train_val |
| Verticillium dahliae | Verticillium_dahliae | GCF_000150675.1_ASM15067v2 | train_val |
| Verticillium nonalfalfae | Verticillium_nonalfalfae | GCF_003724135.2_ASM372413v2 | test |
| Vittaforma corneae | Vittaforma_corneae | GCF_000231115.1_Vitt_corn_V1 | train_val |
| Wallemia ichthyophaga | Wallemia_ichthyophaga | GCF_000400465.1_Wallemia_ichthyophaga_version_1.0 | train_val |
| Wallemia mellicola | Wallemia_mellicola | GCF_000263375.1_Wallemia_sebi_v1.0 | train_val |
| Westerdykella ornata | Westerdykella_ornata | GCF_010094085.1_Wesor1 | test |
| Wickerhamiella sorbophila | Wickerhamiella_sorbophila | GCF_002251995.1_ASM225199v2 | train_val |
| Wickerhamomyces anomalus | Wickerhamomyces_anomalus | GCF_001661255.1_Wican1 | train_val |
| Wickerhamomyces ciferrii | Wickerhamomyces_ciferrii | GCF_000313485.1_ASM31348v1 | train_val |
| Xylona heveae | Xylona_heveae | GCF_001619985.1_Xylona_heveae_TC161_v1.0 | train_val |
| Yamadazyma tenuis | Yamadazyma_tenuis | GCF_000223465.1_Candida_tenuis_v1.0 | train_val |
| Yarrowia lipolytica | Yarrowia_lipolytica | GCF_000002525.2_ASM252v1 | train_val |
| Zasmidium cellare | Zasmidium_cellare | GCF_010093935.1_Zasce1 | train_val |
| Zygosaccharomyces rouxii | Zygosaccharomyces_rouxii | GCF_000026365.1_ASM2636v1 | train_val |
| Zygotorulaspora mrakii | Zygotorulaspora_mrakii | GCF_013402915.1_ASM1340291v1 | test |
| Zymoseptoria tritici | Zymoseptoria_tritici | GCF_000219625.1_MYCGR_v2.0 | train_val |

##### S2.1.2 Plants

Plant training and validation genomes were acquired from Phytozome13 on June 7th 2021.

Plant test genomes, were acquired from RefSeq on July 14th 2022, and were selected as follows.

- exclude any species present in the training validation set
- include genomes with matching or close AUGUSTUS models
- include several more that were expected to have acceptable quality genomes based upon crop impact and researcher experience

Thus, these genomes were selected for comparability with other tools, as well as to have acceptable quality for analyses that rely on the reference (the ablation analyses).

Selection was performed prior to downloading, thus other RefSeq genomes meeting e.g. just the first criteria above are not listed, in contrast to the other phylogenetic groups.

Table S2: All plant genomes, set assignment and versions

| scientific | ID | version | split |
|---|---|---|---|
| Ananas comosus | Acomosus | v3 | train_val |
| Amaranthus hypochondriacus | Ahypochondriacus | v2.1 | train_val |
| Arabidopsis lyrata | Alyrata | v2.1 | train_val |
| Asparagus officinalis | Aofficinalis | V1.1 | train_val |
| Arabidopsis thaliana | Athaliana | TAIR10 | train_val |
| Amborella trichopoda | Atrichopoda | v1.0 | train_val |
| Brachypodium distachyon | Bdistachyon | v3.1 | train_val |
| Brachypodium hybridum | Bhybridum | v1.1 | train_val |
| Brassica oleracea | Boleraceacapitata | v1.0 | train_val |
| Brassica rapa | BrapaFPsc | v1.3 | train_val |
| Beta vulgaris | Bvulgaris | EL10_1.0 | train_val |
| Cicer arietinum | Carietinum | v1.0 | train_val |
| Citrus clementina | Cclementina | v1.0 | train_val |
| Capsella grandiflora | Cgrandiflora | v1.1 | train_val |
| Cinnamomum kanehirae | Ckanehirae | v3 | train_val |
| Carica papaya | Cpapaya | ASGPBv0.4 | train_val |
| Chenopodium quinoa | Cquinoa | v1.0 | train_val |
| Chlamydomonas reinhardtii | Creinhardtii | v5.6 | train_val |
| Capsella rubella | Crubella | v1.1 | train_val |
| Cucumis sativus | Csativus | v1.0 | train_val |
| Citrus sinensis | Csinensis | v1.1 | train_val |
| Coccomyxa subellipsoidea C-169 | CsubellipsoideaC169 | v2.0 | train_val |
| Chromochloris zofingiensis | Czofingiensis | v5.2.3.2 | train_val |
| Dioscorea alata | Dalata | v2.1 | train_val |
| Daucus carota | Dcarota | v2.0 | train_val |
| Dunaliella salina | Dsalina | v1.0 | train_val |
| Eucalyptus grandis | Egrandis | v2.0 | train_val |
| Eutrema salsugineum | Esalsugineum | v1.0 | train_val |
| Fragaria vesca | Fvesca | v4.0.a2 | train_val |
| Glycine max | Gmax | Wm82.a4.v1 | train_val |
| Gossypium raimondii | Graimondii | v2.1 | train_val |
| Glycine soja | Gsoja | v1.1 | train_val |
| Helianthus annuus | Hannuus | r1.2 | train_val |
| Hordeum vulgare | Hvulgare | r1 | train_val |
| Kalanchoe fedtschenkoi | Kfedtschenkoi | v1.1 | train_val |
| Lupinus albus | Lalbus | v1 | train_val |
| Lotus japonicus | Ljaponicus | Lj1.0v1 | train_val |
| Lactuca sativa | Lsativa | v5 | train_val |
| Linum usitatissimum | Lusitatissimum | v1.0 | train_val |
| Musa acuminata | Macuminata | v1 | train_val |
| Malus domestica | Mdomestica | v1.1 | train_val |
| Manihot esculenta | Mesculenta | v8.1 | train_val |
| Mimulus guttatus | Mguttatus | v2.0 | train_val |
| Marchantia polymorpha | Mpolymorpha | v3.1 | train_val |
| Micromonas pusilla | MpusillaCCMP1545 | v3.0 | train_val |
| Micromonas sp. RCC299 | MspRCC299 | v3.0 | train_val |
| Medicago truncatula | Mtruncatula | Mt4.0v1 | train_val |
| Nymphaea colorata | Ncolorata | v1.2 | train_val |
| Olea europaea | Oeuropaea | v1.0 | train_val |
| Ostreococcus lucimarinus | Olucimarinus | v2.0 | train_val |
| Oryza sativa | Osativa | v7.0 | train_val |
| Oropetium thomaeum | Othomaeum | v1.0 | train_val |
| Phaseolus acutifolius | Pacutifolius | v1.0 | train_val |
| Panicum hallii | Phallii | v3.2 | train_val |
| Physcomitrella patens | Ppatens | v3.3 | train_val |
| Prunus persica | Ppersica | v2.1 | train_val |
| Populus trichocarpa | Ptrichocarpa | v4.1 | train_val |
| Poncirus trifoliata | Ptrifoliata | v1.3.1 | train_val |
| Porphyra umbilicalis | Pumbilicalis | v1.5 | train_val |
| Panicum virgatum | Pvirgatum | v5.1 | train_val |
| Ricinus communis | Rcommunis | v0.1 | train_val |
| Sorghum bicolor | Sbicolor | v3.1.1 | train_val |
| Setaria italica | Sitalica | v2.2 | train_val |
| Solanum lycopersicum | Slycopersicum | ITAG3.2 | train_val |
| Selaginella moellendorffii | Smoellendorffii | v1.0 | train_val |
| Schrenkiella parvula | Sparvula | v2.2 | train_val |
| Spirodela polyrhiza | Spolyrhiza | v2 | train_val |
| Salix purpurea | Spurpurea | v1.0 | train_val |
| Solanum tuberosum | Stuberosum | v4.03 | train_val |
| Triticum aestivum | Taestivum | v2.2 | train_val |
| Theobroma cacao | Tcacao | v2.1 | train_val |
| Trifolium pratense | Tpratense | v2 | train_val |
| Volvox carteri | Vcarteri | v2.1 | train_val |
| Vigna unguiculata | Vunguiculata | v1.2 | train_val |
| Vitis vinifera | Vvinifera | v2.1 | train_val |
| Zostera marina | Zmarina | v3.1 | train_val |
| Zea mays | Zmays | RefGen_V4 | train_val |
| Arachis hypogaea | Arachis_hypogaea | GCF_003086295.2_arahy.Tifrunner.gnm1.KYV3 | test (used) |
| Brassica napus | Brassica_napus | GCF_020379485.1_Da-Ae | test (used) |
| Cannabis sativa | Cannabis_sativa | GCF_900626175.2_cs10 | test (used) |

| scientific | ID | version | split |
|---|---|---|---|
| Coffea arabica | Coffea_arabica | GCF_003713225.1_Cara_1.0 | test (used) |
| Hibiscus syriacus | Hibiscus_syriacus | GCF_006381635.1_ASM638163v2 | test (used) |
| Nicotiana attenuata | Nicotiana_attenuata | GCF_001879085.1_NIATTr2 | test (used) |
| Oryza brachyantha | Oryza_brachyantha | GCF_000231095.2_ObraRS2 | test (used) |
| Papaver somniferum | Papaver_somniferum | GCF_003573695.1_ASM357369v1 | test (used) |
| Phoenix dactylifera | Phoenix_dactylifera | GCF_009389715.1_palm_55x_up_171113_PBpolish2nd_... | test (used) |
| Setaria viridis | Setaria_viridis | GCF_005286985.1_Setaria_viridis_v2.0 | test (used) |
| Solanum pennellii | Solanum_pennellii | GCF_001406875.1_SPENNV200 | test (used) |
| Triticum dicoccoides | Triticum_dicoccoides | GCF_002162155.2_WEW_v2.1 | test (used) |
| Vitis riparia | Vitis_riparia | GCF_004353265.1_EGFV_Vit.rip_1.0 | test (used) |

##### S2.1.3 Vertebrates

Vertebrate training, validation, and test genomes were acquired from RefSeq on May 6th, 2022; and are the combined RefSeq sets of 'vertebrate_mammalian' and 'vertebrate_other'.

Species previously used in Stiehler et al. (2020) were assigned to the training and validation set, after that test genomes were assigned randomly *within* genomes that had a matching or close AUGUSTUS trained model available and *within* those that did not.

All species assigned to test that had a close AUGUSTUS model were actually used as test species in this manuscript, additional used test species were selected arbitrarily (every Nth).

Table S3: All vertebrate genomes, set assignment and versions

| scientific | ID | version | split |
|---|---|---|---|
| Acinonyx jubatus | Acinonyx_jubatus | GCF_003709585.1_Aci_jub_2 | train_val |
| Ailuropoda melanoleuca | Ailuropoda_melanoleuca | GCF_002007445.1_ASM200744v2 | train_val |
| Aotus nancymaae | Aotus_nancymaae | GCF_000952055.2_Anan_2.0 | train_val |
| Artibeus jamaicensis | Artibeus_jamaicensis | GCF_014825515.1_WHU_Ajam_v2 | test |
| Arvicanthis niloticus | Arvicanthis_niloticus | GCF_011762505.1_mArvNil1.pat.X | test |
| Arvicola amphibius | Arvicola_amphibius | GCF_903992535.2_mArvAmp1.2 | train_val |
| Balaenoptera acutorostrata | Balaenoptera_acutorostrata | GCF_000493695.1_BalAcu1.0 | test |
| Balaenoptera musculus | Balaenoptera_musculus | GCF_009873245.2_mBalMus1.pri.v3 | test (used) |
| Bison bison | Bison_bison | GCF_000754665.1_Bison_UMD1.0 | train_val |
| Bos indicus | Bos_indicus | GCF_000247795.1_Bos_indicus_1.0 | train_val |
| Bos indicus x Bos taurus | Bos_indicus_x_Bos_taurus | GCF_003369695.1_UOA_Brahman_1 | test |
| Bos mutus | Bos_mutus | GCF_000298355.1_BosGru_v2.0 | train_val |
| Bos taurus | Bos_taurus | GCF_002263795.1_ARS-UCD1.2 | train_val |
| Bubalus bubalis | Bubalus_bubalis | GCF_019923935.1_NDDB_SH_1 | train_val |
| Callithrix jacchus | Callithrix_jacchus | GCF_009663435.1_Callithrix_jacchus_cj1700_1.1 | train_val |
| Callorhinus ursinus | Callorhinus_ursinus | GCF_003265705.1_ASM326570v1 | train_val |
| Camelus bactrianus | Camelus_bactrianus | GCF_000767855.1_Ca_bactrianus_MBC_1.0 | test |
| Camelus dromedarius | Camelus_dromedarius | GCF_000803125.2_CamDro3 | train_val |
| Camelus ferus | Camelus_ferus | GCF_009834535.1_BCGSAC_Cfer_1.0 | test |
| Canis lupus dingo | Canis_lupus_dingo | GCF_012295265.1_UNSW_AlpineDingo_1.0 | train_val |
| Canis lupus familiaris | Canis_lupus_familiaris | GCF_014441545.1_ROS_Cfam_1.0 | train_val |
| Capra hircus | Capra_hircus | GCF_001704415.1_ARS1 | train_val |
| Carlito syrichta | Carlito_syrichta | GCF_000164805.1_Tarsius_syrichta-2.0.1 | train_val |
| Castor canadensis | Castor_canadensis | GCF_001984765.1_C.can_genome_v1.0 | train_val |
| Cavia porcellus | Cavia_porcellus | GCF_000151735.1_Cavpor3.0 | train_val |
| Cebus imitator | Cebus_imitator | GCF_001604975.1_Cebus_imitator-1.0 | train_val |
| Ceratotherium simum | Ceratotherium_simum | GCF_000283155.1_CerSimSim1.0 | train_val |
| Cercocebus atys | Cercocebus_atys | GCF_000955945.1_Caty_1.0 | train_val |
| Cervus canadensis | Cervus_canadensis | GCF_019320065.1_ASM1932006v1 | test |
| Cervus elaphus | Cervus_elaphus | GCF_910594005.1_mCerEla1.1 | train_val |
| Chinchilla lanigera | Chinchilla_lanigera | GCF_000276665.1_ChiLan1.0 | train_val |
| Chlorocebus sabaeus | Chlorocebus_sabaeus | GCF_015252025.1_Vero_WHO_p1.0 | train_val |
| Choloepus didactylus | Choloepus_didactylus | GCF_015220235.1_mChoDid1.pri | test |
| Chrysochloris asiatica | Chrysochloris_asiatica | GCF_000296735.1_ChrAsi1.0 | train_val |
| Colobus angolensis | Colobus_angolensis | GCF_000951035.1_Cang.pa_1.0 | train_val |
| Condylura cristata | Condylura_cristata | GCF_000260355.1_ConCri1.0 | test |
| Cricetulus griseus | Cricetulus_griseus | GCF_000223135.1_CriGri_1.0 | train_val |
| Dasypus novemcinctus | Dasypus_novemcinctus | GCF_000208655.1_Dasnov3.0 | train_val |
| Delphinapterus leucas | Delphinapterus_leucas | GCF_002288925.2_ASM228892v3 | train_val |
| Desmodus rotundus | Desmodus_rotundus | GCF_002940915.1_ASM294091v2 | test (used) |
| Dipodomys ordii | Dipodomys_ordii | GCF_000151885.1_Dord_2.0 | train_val |
| Dipodomys spectabilis | Dipodomys_spectabilis | GCF_019054845.1_ASM1905484v1 | test |
| Dromiciops gliroides | Dromiciops_gliroides | GCF_019393635.1_mDroGli1.pri | train_val |
| Echinops telfairi | Echinops_telfairi | GCF_000313985.2_ASM31398v2 | train_val |
| Elephantulus edwardii | Elephantulus_edwardii | GCF_000299155.1_EleEdw1.0 | test |
| Enhydra lutris | Enhydra_lutris | GCF_002288905.1_ASM228890v2 | test |
| Eptesicus fuscus | Eptesicus_fuscus | GCF_000308155.1_EptFus1.0 | test |
| Equus asinus | Equus_asinus | GCF_016077325.2_ASM1607732v2 | train_val |
| Equus caballus | Equus_caballus | GCF_002863925.1_EquCab3.0 | train_val |
| Equus przewalskii | Equus_przewalskii | GCF_000696695.1_Burgud | test |
| Erinaceus europaeus | Erinaceus_europaeus | GCF_000296755.1_EriEur2.0 | train_val |
| Eumetopias jubatus | Eumetopias_jubatus | GCF_004028035.1_ASM402803v1 | test |
| Felis catus | Felis_catus | GCF_018350175.1_F.catus_Fca126_mat1.0 | train_val |
| Fukomys damarensis | Fukomys_damarensis | GCF_012274545.1_DMR_v1.0_HiC | train_val |
| Galeopterus variegatus | Galeopterus_variegatus | GCF_000696425.1_G_variegatus-3.0.2 | test |
| Globicephala melas | Globicephala_melas | GCF_006547405.1_ASM654740v1 | test |
| Gorilla gorilla | Gorilla_gorilla | GCF_008122165.1_Kamilah_GGO_v0 | train_val |
| Gracilinanus agilis | Gracilinanus_agilis | GCF_016433145.1_AgileGrace | test |
| Grammomys surdaster | Grammomys_surdaster | GCF_004785775.1_NIH_TR_1.0 | test |
| Halichoerus grypus | Halichoerus_grypus | GCF_012393455.1_Tufts_HGry_1.1 | test |
| Heterocephalus glaber | Heterocephalus_glaber | GCF_000247695.1_HetGla_female_1.0 | train_val |
| Hipposideros armiger | Hipposideros_armiger | GCF_001890085.1_ASM189008v1 | train_val |
| Homo sapiens | Homo_sapiens | GCF_000001405.39_GRCh38.p13 | train_val |
| Hyaena hyaena | Hyaena_hyaena | GCF_003009895.1_ASM300989v1 | train_val |
| Hylobates moloch | Hylobates_moloch | GCF_009828535.2_HMol_V2 | train_val |
| Ictidomys tridecemlineatus | Ictidomys_tridecemlineatus | GCF_016881025.1_HiC_Itri_2 | train_val |
| Jaculus jaculus | Jaculus_jaculus | GCF_020740685.1_mJacJac1.mat.Y.cur | train_val |
| Lagenorhynchus obliquidens | Lagenorhynchus_obliquidens | GCF_003676395.1_ASM367639v1 | test |
| Lemur catta | Lemur_catta | GCF_020740605.2_mLemCat1.pri | train_val |
| Leopardus geoffroyi | Leopardus_geoffroyi | GCF_018350155.1_O.geoffroyi_Oge1_pat1.0 | test |
| Leptonychotes weddellii | Leptonychotes_weddellii | GCF_000349705.1_LepWed1.0 | train_val |
| Lipotes vexillifer | Lipotes_vexillifer | GCF_000442215.1_Lipotes_vexillifer_v1 | train_val |
| Lontra canadensis | Lontra_canadensis | GCF_010015895.1_GSC_riverotter_1.0 | train_val |
| Loxodonta africana | Loxodonta_africana | GCF_000001905.1_Loxafr3.0 | train_val |
| Lynx canadensis | Lynx_canadensis | GCF_007474595.2_mLynCan4.pri.v2 | train_val |
| Macaca fascicularis | Macaca_fascicularis | GCF_012559485.2_MFA1912RKSv2 | train_val |
| Macaca mulatta | Macaca_mulatta | GCF_003339765.1_Mmul_10 | train_val |
| Macaca nemestrina | Macaca_nemestrina | GCF_000956065.1_Mnem_1.0 | train_val |
| Mandrillus leucophaeus | Mandrillus_leucophaeus | GCF_000951045.1_Mleu.le_1.0 | train_val |
| Manis javanica | Manis_javanica | GCF_014570535.1_YNU_ManJav_2.0 | test |

| scientific | ID | version | split |
|---|---|---|---|
| Manis pentadactyla | Manis_pentadactyla | GCF_014570555.1_YNU_ManPten_2.0 | test |
| Marmota flaviventris | Marmota_flaviventris | GCF_003676075.2_GSC_YBM_2.0 | test |
| Marmota marmota | Marmota_marmota | GCF_001458135.1_marMar2.1 | train_val |
| Marmota monax | Marmota_monax | GCF_021218885.1_Marmota_monax_Labrador192_V1.0 | test |
| Mastomys coucha | Mastomys_coucha | GCF_008632895.1_UCSF_Mcou_1 | test |
| Meles meles | Meles_meles | GCF_922984935.1_mMelMel3.1_paternal_haplotype | train_val |
| Meriones unguiculatus | Meriones_unguiculatus | GCF_002204375.1_MunDraft-v1.0 | train_val |
| Mesocricetus auratus | Mesocricetus_auratus | GCF_017639785.1_BCM_Maur_2.0 | train_val |
| Microcebus murinus | Microcebus_murinus | GCF_000165445.2_Mmur_3.0 | train_val |
| Microtus ochrogaster | Microtus_ochrogaster | GCF_000317375.1_MicOch1.0 | train_val |
| Microtus oregoni | Microtus_oregoni | GCF_018167655.1_Mior012 | train_val |
| Miniopterus natalensis | Miniopterus_natalensis | GCF_001595765.1_Mnat.v1 | test |
| Mirounga angustirostris | Mirounga_angustirostris | GCF_021288785.1_ASM2128878v2 | test |
| Mirounga leonina | Mirounga_leonina | GCF_011800145.1_KU_Mleo_1.0 | train_val |
| Molossus molossus | Molossus_molossus | GCF_014108415.1_mMolMol1.p | test |
| Monodelphis domestica | Monodelphis_domestica | GCF_000002295.2_MonDom5 | train_val |
| Monodon monoceros | Monodon_monoceros | GCF_005190385.1_NGI_Narwhal_1 | test |
| Mus caroli | Mus_caroli | GCF_900094665.1_CAROLI_EIJ_v1.1 | train_val |
| Mus musculus | Mus_musculus | GCF_000001635.27_GRCm39 | train_val |
| Mus pahari | Mus_pahari | GCF_900095145.1_PAHARI_EIJ_v1.1 | train_val |
| Mustela erminea | Mustela_erminea | GCF_009829155.1_mMusErm1.Pri | test |
| Mustela putorius | Mustela_putorius | GCF_011764305.1_ASM1176430v1.1 | train_val |
| Myotis brandtii | Myotis_brandtii | GCF_000412655.1_ASM41265v1 | train_val |
| Myotis davidii | Myotis_davidii | GCF_000327345.1_ASM32734v1 | test (used) |
| Myotis lucifugus | Myotis_lucifugus | GCF_000147115.1_Myoluc2.0 | train_val |
| Myotis myotis | Myotis_myotis | GCF_014108235.1_mMyoMyo1.p | train_val |
| Nannospalax galili | Nannospalax_galili | GCF_000622305.1_S.galili_v1.0 | train_val |
| Neogale vison | Neogale_vison | GCF_020171115.1_ASM_NN_V1 | train_val |
| Neomonachus schauinslandi | Neomonachus_schauinslandi | GCF_002201575.2_ASM220157v2 | train_val |
| Neophocaena asiaeorientalis | Neophocaena_asiaeorientalis | GCF_003031525.2_Neophocaena_asiaeorientalis_V1 | test |
| Nomascus leucogenys | Nomascus_leucogenys | GCF_006542625.1_Asia_NLE_v1 | train_val |
| Ochotona curzoniae | Ochotona_curzoniae | GCF_017591425.1_NIBS_Ocur_1.0 | train_val |
| Ochotona princeps | Ochotona_princeps | GCF_014633375.1_OchPri4.0 | train_val |
| Octodon degus | Octodon_degus | GCF_000260255.1_OctDeg1.0 | train_val |
| Odobenus rosmarus | Odobenus_rosmarus | GCF_000321225.1_Oros_1.0 | train_val |
| Odocoileus virginianus | Odocoileus_virginianus | GCF_002102435.1_Ovir.te_1.0 | test |
| Onychomys torridus | Onychomys_torridus | GCF_903995425.1_mOncTor1.1 | test |
| Orcinus orca | Orcinus_orca | GCF_000331955.2_Oorc_1.1 | train_val |
| Ornithorhynchus anatinus | Ornithorhynchus_anatinus | GCF_004115215.2_mOrnAna1.pri.v4 | train_val |
| Orycteropus afer | Orycteropus_afer | GCF_000298275.1_OryAfe1.0 | test |
| Oryctolagus cuniculus | Oryctolagus_cuniculus | GCF_000003625.3_OryCun2.0 | train_val |
| Oryx dammah | Oryx_dammah | GCF_014754425.2_SCBI_Odam_1.1 | test |
| Otolemur garnettii | Otolemur_garnettii | GCF_000181295.1_OtoGar3 | train_val |
| Ovis aries | Ovis_aries | GCF_016772045.1_ARS-UI_Ramb_v2.0 | train_val |
| Pan paniscus | Pan_paniscus | GCF_013052645.1_Mhudiblu_PPA_v0 | train_val |
| Panthera leo | Panthera_leo | GCF_018350215.1_P.leo_Ple1_pat1.1 | train_val |
| Panthera pardus | Panthera_pardus | GCF_001857705.1_PanPar1.0 | train_val |
| Panthera tigris | Panthera_tigris | GCF_018350195.1_P.tigris_Pti1_mat1.1 | train_val |
| Pan troglodytes | Pan_troglodytes | GCF_002880755.1_Clint_PTRv2 | train_val |
| Papio anubis | Papio_anubis | GCF_008728515.1_Panubis1.0 | train_val |
| Peromyscus leucopus | Peromyscus_leucopus | GCF_004664715.2_UCI_PerLeu_2.1 | test |
| Peromyscus maniculatus | Peromyscus_maniculatus | GCF_003704035.1_HU_Pman_2.1.3 | train_val |
| Phascolarctos cinereus | Phascolarctos_cinereus | GCF_002099425.1_phaCin_unsw_v4.1 | train_val |
| Phoca vitulina | Phoca_vitulina | GCF_004348235.1_GSC_HSeal_1.0 | train_val |
| Phocoena sinus | Phocoena_sinus | GCF_008692025.1_mPhoSin1.pri | train_val |
| Phyllostomus discolor | Phyllostomus_discolor | GCF_004126475.2_mPhyDis1.pri.v3 | train_val |
| Phyllostomus hastatus | Phyllostomus_hastatus | GCF_019186645.2_TTU_PhHast_1.1 | train_val |
| Physeter catodon | Physeter_catodon | GCF_002837175.2_ASM283717v2 | test |
| Piliocolobus tephrosceles | Piliocolobus_tephrosceles | GCF_002776525.3_ASM277652v3 | train_val |
| Pipistrellus kuhlii | Pipistrellus_kuhlii | GCF_014108245.1_mPipKuh1.p | train_val |
| Pongo abelii | Pongo_abelii | GCF_002880775.1_Susie_PABv2 | train_val |
| Prionailurus bengalensis | Prionailurus_bengalensis | GCF_016509475.1_Fcat_Pben_1.1_paternal_pri | test |
| Propithecus coquereli | Propithecus_coquereli | GCF_000956105.1_Pcoq_1.0 | train_val |
| Pteropus alecto | Pteropus_alecto | GCF_000325575.1_ASM32557v1 | train_val |
| Pteropus giganteus | Pteropus_giganteus | GCF_902729225.1_Ma_sr-lr_union100 | test |
| Pteropus vampyrus | Pteropus_vampyrus | GCF_000151845.1_Pvam_2.0 | train_val |
| Puma concolor | Puma_concolor | GCF_003327715.1_PumCon1.0 | train_val |
| Puma yagouaroundi | Puma_yagouaroundi | GCF_014898765.1_PumYag | train_val |
| Rattus norvegicus | Rattus_norvegicus | GCF_015227675.2_mRatBN7.2 | train_val |
| Rattus rattus | Rattus_rattus | GCF_011064425.1_Rrattus_CSIRO_v1 | train_val |
| Rhinolophus ferrumequinum | Rhinolophus_ferrumequinum | GCF_004115265.1_mRhiFer1_v1.p | test |
| Rhinopithecus bieti | Rhinopithecus_bieti | GCF_001698545.1_ASM169854v1 | train_val |
| Rhinopithecus roxellana | Rhinopithecus_roxellana | GCF_007565055.1_ASM756505v1 | train_val |
| Rousettus aegyptiacus | Rousettus_aegyptiacus | GCF_014176215.1_mRouAeg1.p | test |
| Saimiri boliviensis | Saimiri_boliviensis | GCF_016699345.1_BCM_Sbol_2.0 | train_val |
| Sapajus apella | Sapajus_apella | GCF_009761245.1_GSC_monkey_1.0 | test |
| Sarcophilus harrisii | Sarcophilus_harrisii | GCF_902635505.1_mSarHar1.11 | train_val |
| Sorex araneus | Sorex_araneus | GCF_000181275.1_SorAra2.0 | train_val |
| Sturnira hondurensis | Sturnira_hondurensis | GCF_014824575.2_WHU_Shon_v2.1 | train_val |
| Suricata suricatta | Suricata_suricatta | GCF_006229205.1_meerkat_22Aug2017_6uvM2_HiC | test |

| scientific | ID | version | split |
|---|---|---|---|
| Sus scrofa | Sus_scrofa | GCF_000003025.6_Sscrofa11.1 | train_val |
| Tachyglossus aculeatus | Tachyglossus_aculeatus | GCF_015852505.1_mTacAcu1.pri | train_val |
| Talpa occidentalis | Talpa_occidentalis | GCF_014898055.1_MPIMG_talOcc4 | train_val |
| Theropithecus gelada | Theropithecus_gelada | GCF_003255815.1_Tgel_1.0 | train_val |
| Trachypithecus francoisi | Trachypithecus_francoisi | GCF_009764315.1_Tfra_2.0 | test |
| Trichechus manatus | Trichechus_manatus | GCF_000243295.1_TriManLat1.0 | test |
| Trichosurus vulpecula | Trichosurus_vulpecula | GCF_011100635.1_mTriVul1.pri | test |
| Tupaia chinensis | Tupaia_chinensis | GCF_000334495.1_TupChi_1.0 | train_val |
| Tursiops truncatus | Tursiops_truncatus | GCF_011762595.1_mTurTru1.mat.Y | train_val |
| Urocitellus parryii | Urocitellus_parryii | GCF_003426925.1_ASM342692v1 | train_val |
| Ursus americanus | Ursus_americanus | GCF_020975775.1_gsc_jax_bbear_1.0 | train_val |
| Ursus arctos | Ursus_arctos | GCF_003584765.2_ASM358476v2 | train_val |
| Ursus maritimus | Ursus_maritimus | GCF_017311325.1_ASM1731132v1 | train_val |
| Vicugna pacos | Vicugna_pacos | GCF_000164845.3_VicPac3.1 | train_val |
| Vombatus ursinus | Vombatus_ursinus | GCF_900497805.2_bare-nosed_wombat_genome_assembly | train_val |
| Vulpes lagopus | Vulpes_lagopus | GCF_018345385.1_ASM1834538v1 | test |
| Vulpes vulpes | Vulpes_vulpes | GCF_003160815.1_VulVul2.2 | train_val |
| Zalophus californianus | Zalophus_californianus | GCF_009762305.2_mZalCal1.pri.v2 | test |
| Acanthisitta chloris | Acanthisitta_chloris | GCF_000695815.1_ASM69581v1 | test |
| Acanthochromis polyacanthus | Acanthochromis_polyacanthus | GCF_002109545.1_ASM210954v1 | train_val |
| Acanthopagrus latus | Acanthopagrus_latus | GCF_904848185.1_fAcaLat1.1 | train_val |
| Acipenser ruthenus | Acipenser_ruthenus | GCF_010645085.1_ASM1064508v1 | test |
| Alligator mississippiensis | Alligator_mississippiensis | GCF_000281125.3_ASM28112v4 | train_val |
| Alligator sinensis | Alligator_sinensis | GCF_000455745.1_ASM45574v1 | test |
| Alosa sapidissima | Alosa_sapidissima | GCF_018492685.1_fAloSap1.pri | train_val |
| Amblyraja radiata | Amblyraja_radiata | GCF_010909765.2_sAmbRad1.1.pri | test |
| Amphiprion ocellaris | Amphiprion_ocellaris | GCF_002776465.1_AmpOce1.0 | train_val |
| Anabas testudineus | Anabas_testudineus | GCF_900324465.2_fAnaTes1.2 | train_val |
| Anarrhichthys ocellatus | Anarrhichthys_ocellatus | GCF_004355925.1_GSC_Weel_1.0 | train_val |
| Anas platyrhynchos | Anas_platyrhynchos | GCF_015476345.1_ZJU1.0 | train_val |
| Anguilla anguilla | Anguilla_anguilla | GCF_013347855.1_fAngAng1.pri | test |
| Anolis carolinensis | Anolis_carolinensis | GCF_000090745.1_AnoCar2.0 | train_val |
| Anser cygnoides | Anser_cygnoides | GCF_000971095.1_AnsCyg_PRJNA183603_v1.0 | train_val |
| Antrostomus carolinensis | Antrostomus_carolinensis | GCF_000700745.1_ASM70074v1 | train_val |
| Apaloderma vittatum | Apaloderma_vittatum | GCF_000703405.1_ASM70340v1 | test |
| Aptenodytes forsteri | Aptenodytes_forsteri | GCF_000699145.1_ASM69914v1 | test |
| Apteryx mantelli | Apteryx_mantelli | GCF_001039765.1_AptMant0 | test |
| Apteryx rowi | Apteryx_rowi | GCF_003343035.1_aptRow1 | train_val |
| Aquila chrysaetos | Aquila_chrysaetos | GCF_900496995.4_bAquChr1.4 | train_val |
| Archocentrus centrarchus | Archocentrus_centrarchus | GCF_007364275.1_fArcCen1 | train_val |
| Astatotilapia calliptera | Astatotilapia_calliptera | GCF_900246225.1_fAstCal1.2 | train_val |
| Astyanax mexicanus | Astyanax_mexicanus | GCF_000372685.2_Astyanax_mexicanus-2.0 | train_val |
| Athene cunicularia | Athene_cunicularia | GCF_003259725.1_athCun1 | train_val |
| Austrofundulus limnaeus | Austrofundulus_limnaeus | GCF_001266775.1_Austrofundulus_limnaeus-1.0 | train_val |
| Aythya fuligula | Aythya_fuligula | GCF_009819795.1_bAytFul2.pri | test |
| Balearica regulorum | Balearica_regulorum | GCF_000709895.1_ASM70989v1 | test |
| Betta splendens | Betta_splendens | GCF_900634795.3_fBetSpl5.3 | train_val |
| Boleophthalmus pectinirostris | Boleophthalmus_pectinirostris | GCF_000788275.1_BP.fa | train_val |
| Buceros rhinoceros | Buceros_rhinoceros | GCF_000710305.1_ASM71030v1 | test |
| Bufo bufo | Bufo_bufo | GCF_905171765.1_aBufBuf1.1 | test |
| Bufo gargarizans | Bufo_gargarizans | GCF_014858855.1_ASM1485885v1 | test |
| Calidris pugnax | Calidris_pugnax | GCF_001431845.1_ASM143184v1 | train_val |
| Callorhinchus milii | Callorhinchus_milii | GCF_018977255.1_IMCB_Cmil_1.0 | train_val |
| Calypte anna | Calypte_anna | GCF_003957555.1_bCalAnn1_v1.p | train_val |
| Camarhynchus parvulus | Camarhynchus_parvulus | GCF_901933205.1_STF_HiC | train_val |
| Carassius auratus | Carassius_auratus | GCF_003368295.1_ASM336829v1 | train_val |
| Carcharodon carcharias | Carcharodon_carcharias | GCF_017639515.1_sCarCar2.pri | test |
| Cariama cristata | Cariama_cristata | GCF_000690535.1_ASM69053v1 | train_val |
| Catharus ustulatus | Catharus_ustulatus | GCF_009819885.2_bCatUst1.pri.v2 | train_val |
| Centrocercus urophasianus | Centrocercus_urophasianus | GCF_019232065.1_USGS_Curo_1.0 | train_val |
| Chaetura pelagica | Chaetura_pelagica | GCF_000747805.1_ChaPel_1.0 | train_val |
| Chanos chanos | Chanos_chanos | GCF_902362185.1_fChaCha1.1 | test |
| Charadrius vociferus | Charadrius_vociferus | GCF_000708025.1_ASM70802v2 | train_val |
| Cheilinus undulatus | Cheilinus_undulatus | GCF_018320785.1_ASM1832078v1 | train_val |
| Chelmon rostratus | Chelmon_rostratus | GCF_017976325.1_fCheRos1.pri | test |
| Chelonia mydas | Chelonia_mydas | GCF_015237465.2_rCheMyd1.pri.v2 | test |
| Chelonoidis abingdonii | Chelonoidis_abingdonii | GCF_003597395.1_ASM359739v1 | train_val |
| Chiloscyllium plagiosum | Chiloscyllium_plagiosum | GCF_004010195.1_ASM401019v2 | train_val |
| Chiroxiphia lanceolata | Chiroxiphia_lanceolata | GCF_009829145.1_bChiLan1.pri | test (used) |
| Chlamydotis macqueenii | Chlamydotis_macqueenii | GCF_000695195.1_ASM69519v1 | train_val |
| Chrysemys picta | Chrysemys_picta | GCF_000241765.4_Chrysemys_picta_BioNano-3.0.4 | train_val |
| Clupea harengus | Clupea_harengus | GCF_900700415.2_Ch_v2.0.2 | train_val |
| Colius striatus | Colius_striatus | GCF_000690715.1_ASM69071v1 | test |
| Colossoma macropomum | Colossoma_macropomum | GCF_904425465.1_Colossoma_macropomum | test |
| Columba livia | Columba_livia | GCF_000337935.1_Cliv_1.0 | train_val |
| Corapipo altera | Corapipo_altera | GCF_003945725.1_ASM394572v1 | test |
| Coregonus clupeaformis | Coregonus_clupeaformis | GCF_020615455.1_ASM2061545v1 | test |
| Corvus brachyrhynchos | Corvus_brachyrhynchos | GCF_000691975.1_ASM69197v1 | test |
| Corvus cornix | Corvus_cornix | GCF_000738735.5_ASM73873v5 | test |
| Corvus kubaryi | Corvus_kubaryi | GCF_017639235.1_C.kubaryi_AGA036_p1.0 | test |

| scientific | ID | version | split |
|---|---|---|---|
| Corvus moneduloides | Corvus_moneduloides | GCF_009650955.1_bCorMon1.pri | train_val |
| Cottoperca gobio | Cottoperca_gobio | GCF_900634415.1_fCotGob3.1 | train_val |
| Coturnix japonica | Coturnix_japonica | GCF_001577835.2_Coturnix_japonica_2.1 | train_val |
| Crocodylus porosus | Crocodylus_porosus | GCF_001723895.1_CroPor_comp1 | train_val |
| Crotalus tigris | Crotalus_tigris | GCF_016545835.1_ASM1654583v1 | test |
| Cuculus canorus | Cuculus_canorus | GCF_000709325.1_ASM70932v1 | test |
| Cyanistes caeruleus | Cyanistes_caeruleus | GCF_002901205.1_cyaCae2 | train_val |
| Cyclopterus lumpus | Cyclopterus_lumpus | GCF_009769545.1_fCycLum1.pri | train_val |
| Cygnus atratus | Cygnus_atratus | GCF_013377495.1_Cygnus_atratus_primary_v1.0 | train_val |
| Cygnus olor | Cygnus_olor | GCF_009769625.2_bCygOlo1.pri.v2 | train_val |
| Cynoglossus semilaevis | Cynoglossus_semilaevis | GCF_000523025.1_Cse_v1.0 | train_val |
| Cyprinodon tularosa | Cyprinodon_tularosa | GCF_016077235.1_ASM1607723v1 | train_val |
| Cyprinodon variegatus | Cyprinodon_variegatus | GCF_000732505.1_C_variegatus-1.0 | train_val |
| Cyprinus carpio | Cyprinus_carpio | GCF_018340385.1_ASM1834038v1 | train_val |
| Danio rerio | Danio_rerio | GCF_000002035.6_GRCz11 | train_val |
| Denticeps clupeoides | Denticeps_clupeoides | GCF_900700375.1_fDenClu1.1 | train_val |
| Dermochelys coriacea | Dermochelys_coriacea | GCF_009764565.3_rDerCor1.pri.v4 | test |
| Dromaius novaehollandiae | Dromaius_novaehollandiae | GCF_003342905.1_droNov1 | train_val |
| Dryobates pubescens | Dryobates_pubescens | GCF_000699005.1_ASM69900v1 | test |
| Echeneis naucrates | Echeneis_naucrates | GCF_900963305.1_fEcheNa1.1 | train_val |
| Egretta garzetta | Egretta_garzetta | GCF_000687185.1_ASM68718v1 | test |
| Electrophorus electricus | Electrophorus_electricus | GCF_013358815.1_fEleEle1.pri | train_val |
| Empidonax traillii | Empidonax_traillii | GCF_003031625.1_ASM303162v1 | test |
| Epinephelus lanceolatus | Epinephelus_lanceolatus | GCF_005281545.1_ASM528154v1 | test |
| Erpetoichthys calabaricus | Erpetoichthys_calabaricus | GCF_900747795.1_fErpCal1.1 | train_val |
| Esox lucius | Esox_lucius | GCF_011004845.1_fEsoLuc1.pri | train_val |
| Etheostoma cragini | Etheostoma_cragini | GCF_013103735.1_CSU_Ecrag_1.0 | test |
| Etheostoma spectabile | Etheostoma_spectabile | GCF_008692095.1_UIUC_Espe_1.0 | train_val |
| Eurypyga helias | Eurypyga_helias | GCF_000690775.1_ASM69077v1 | train_val |
| Falco cherrug | Falco_cherrug | GCF_000337975.1_F_cherrug_v1.0 | test |
| Falco naumanni | Falco_naumanni | GCF_017639655.2_bFalNau1.pat | test (used) |
| Falco peregrinus | Falco_peregrinus | GCF_000337955.1_F_peregrinus_v1.0 | train_val |
| Falco rusticolus | Falco_rusticolus | GCF_015220075.1_bFalRus1.pri | test |
| Ficedula albicollis | Ficedula_albicollis | GCF_000247815.1_FicAlb1.5 | train_val |
| Fulmarus glacialis | Fulmarus_glacialis | GCF_000690835.1_ASM69083v1 | train_val |
| Fundulus heteroclitus | Fundulus_heteroclitus | GCF_011125445.2_MU-UCD_Fhet_4.1 | train_val |
| Gadus morhua | Gadus_morhua | GCF_902167405.1_gadMor3.0 | train_val |
| Gallus gallus | Gallus_gallus | GCF_016699485.2_bGalGal1.mat.broiler.GRCg7b | train_val |
| Gambusia affinis | Gambusia_affinis | GCF_019740435.1_SWU_Gaff_1.0 | train_val |
| Gasterosteus aculeatus | Gasterosteus_aculeatus | GCF_016920845.1_GAculeatus_UGA_version5 | train_val |
| Gavialis gangeticus | Gavialis_gangeticus | GCF_001723915.1_GavGan_comp1 | train_val |
| Gavia stellata | Gavia_stellata | GCF_000690875.1_ASM69087v1 | test |
| Gekko japonicus | Gekko_japonicus | GCF_001447785.1_Gekko_japonicus_V1.1 | test |
| Geospiza fortis | Geospiza_fortis | GCF_000277835.1_GeoFor_1.0 | train_val |
| Geotrypetes seraphini | Geotrypetes_seraphini | GCF_902459505.1_aGeoSer1.1 | train_val |
| Gopherus evgoodei | Gopherus_evgoodei | GCF_007399415.2_rGopEvg1_v1.p | train_val |
| Gouania willdenowi | Gouania_willdenowi | GCF_900634775.1_fGouWil2.1 | train_val |
| Gymnodraco acuticeps | Gymnodraco_acuticeps | GCF_902827175.1_fGymAcu1.1 | test |
| Haliaeetus albicilla | Haliaeetus_albicilla | GCF_000691405.1_ASM69140v1 | train_val |
| Haliaeetus leucocephalus | Haliaeetus_leucocephalus | GCF_000737465.1_Haliaeetus_leucocephalus-4.0 | test |
| Haplochromis burtoni | Haplochromis_burtoni | GCF_018398535.1_NCSU_Asbu1 | train_val |
| Hippocampus comes | Hippocampus_comes | GCF_001891065.1_H_comes_QL1_v1 | train_val |
| Hippoglossus hippoglossus | Hippoglossus_hippoglossus | GCF_009819705.1_fHipHip1.pri | test |
| Hippoglossus stenolepis | Hippoglossus_stenolepis | GCF_013339905.1_IPHC_HiSten_1.0 | train_val |
| Hirundo rustica | Hirundo_rustica | GCF_015227805.1_bHirRus1.pri.v2 | train_val |
| Ictalurus punctatus | Ictalurus_punctatus | GCF_001660625.1_IpCoco_1.2 | train_val |
| Kryptolebias marmoratus | Kryptolebias_marmoratus | GCF_001649575.2_ASM164957v2 | train_val |
| Labrus bergylta | Labrus_bergylta | GCF_900080235.1_BallGen_V1 | train_val |
| Lacerta agilis | Lacerta_agilis | GCF_009819535.1_rLacAgi1.pri | train_val |
| Lagopus leucura | Lagopus_leucura | GCF_019238085.1_USGS_WTPT01 | test (used) |
| Larimichthys crocea | Larimichthys_crocea | GCF_000972845.2_L_crocea_2.0 | train_val |
| Lates calcarifer | Lates_calcarifer | GCF_001640805.1_ASM164080v1 | train_val |
| Latimeria chalumnae | Latimeria_chalumnae | GCF_000225785.1_LatCha1 | train_val |
| Lepidothrix coronata | Lepidothrix_coronata | GCF_001604755.1_Lepidothrix_coronata-1.0 | train_val |
| Lepisosteus oculatus | Lepisosteus_oculatus | GCF_000242695.1_LepOcu1 | train_val |
| Leptosomus discolor | Leptosomus_discolor | GCF_000691785.1_ASM69178v1 | train_val |
| Lonchura striata | Lonchura_striata | GCF_005870125.1_lonStrDom2 | train_val |
| Manacus vitellinus | Manacus_vitellinus | GCF_001715985.3_ASM171598v3 | train_val |
| Mastacembelus armatus | Mastacembelus_armatus | GCF_900324485.2_fMasArm1.2 | train_val |
| Mauremys mutica | Mauremys_mutica | GCF_020497125.1_ASM2049712v1 | train_val |
| Mauremys reevesii | Mauremys_reevesii | GCF_016161935.1_ASM1616193v1 | train_val |
| Maylandia zebra | Maylandia_zebra | GCF_000238955.4_M_zebra_UMD2a | train_val |
| Megalops cyprinoides | Megalops_cyprinoides | GCF_013368585.1_fMegCyp1.pri | train_val |
| Melanotaenia boesemani | Melanotaenia_boesemani | GCF_017639745.1_fMelBoe1.pri | train_val |
| Meleagris gallopavo | Meleagris_gallopavo | GCF_000146605.3_Turkey_5.1 | train_val |
| Melopsittacus undulatus | Melopsittacus_undulatus | GCF_012275295.1_bMelUnd1.mat.Z | train_val |
| Merops nubicus | Merops_nubicus | GCF_000691845.1_ASM69184v1 | test |
| Mesitornis unicolor | Mesitornis_unicolor | GCF_000695765.1_ASM69576v1 | train_val |
| Microcaecilia unicolor | Microcaecilia_unicolor | GCF_901765095.1_aMicUni1.1 | train_val |
| Micropterus dolomieu | Micropterus_dolomieu | GCF_021292245.1_ASM2129224v1 | train_val |

| scientific | ID | version | split |
|---|---|---|---|
| Micropterus salmoides | Micropterus_salmoides | GCF_014851395.1_ASM1485139v1 | train_val |
| Molothrus ater | Molothrus_ater | GCF_012460135.1_BPBGC_Mater_1.0 | train_val |
| Monopterus albus | Monopterus_albus | GCF_001952655.1_M_albus_1.0 | train_val |
| Morone saxatilis | Morone_saxatilis | GCF_004916995.1_NCSU_SB_2.0 | train_val |
| Motacilla alba | Motacilla_alba | GCF_015832195.1_Motacilla_alba_V1.0_pri | train_val |
| Myripristis murdjan | Myripristis_murdjan | GCF_902150065.1_fMyrMur1.1 | train_val |
| Nanorana parkeri | Nanorana_parkeri | GCF_000935625.1_ASM93562v1 | test |
| Nematolebias whitei | Nematolebias_whitei | GCF_014905685.2_NemWhi1 | train_val |
| Neolamprologus brichardi | Neolamprologus_brichardi | GCF_000239395.1_NeoBri1.0 | train_val |
| Neopelma chrysocephalum | Neopelma_chrysocephalum | GCF_003984885.1_ASM398488v2 | test |
| Nestor notabilis | Nestor_notabilis | GCF_000696875.1_ASM69687v1 | train_val |
| Nipponia nippon | Nipponia_nippon | GCF_000708225.1_ASM70822v1 | train_val |
| Notechis scutatus | Notechis_scutatus | GCF_900518725.1_TS10Xv2-PRI | train_val |
| Nothobranchius furzeri | Nothobranchius_furzeri | GCF_001465895.1_Nfu_20140520 | train_val |
| Nothoprocta perdicaria | Nothoprocta_perdicaria | GCF_003342845.1_notPer1 | train_val |
| Notolabrus celidotus | Notolabrus_celidotus | GCF_009762535.1_fNotCel1.pri | test |
| Notothenia coriiceps | Notothenia_coriiceps | GCF_000735185.1_NC01 | test |
| Numida meleagris | Numida_meleagris | GCF_002078875.1_NumMel1.0 | train_val |
| Oncorhynchus gorbuscha | Oncorhynchus_gorbuscha | GCF_021184085.1_OgorEven_v1.0 | test |
| Oncorhynchus keta | Oncorhynchus_keta | GCF_012931545.1_Oket_V1 | train_val |
| Oncorhynchus kisutch | Oncorhynchus_kisutch | GCF_002021735.2_Okis_V2 | test |
| Oncorhynchus mykiss | Oncorhynchus_mykiss | GCF_013265735.2_USDA_OmykA_1.1 | test |
| Oncorhynchus nerka | Oncorhynchus_nerka | GCF_006149115.1_Oner_1.0 | train_val |
| Oncorhynchus tshawytscha | Oncorhynchus_tshawytscha | GCF_018296145.1_Otsh_v2.0 | test |
| Onychostruthus taczanowskii | Onychostruthus_taczanowskii | GCF_017590055.1_ASM1759005v1 | test |
| Opisthocomus hoazin | Opisthocomus_hoazin | GCF_000692075.1_ASM69207v1 | test (used) |
| Oreochromis aureus | Oreochromis_aureus | GCF_013358895.1_ZZ_aureus | test |
| Oreochromis niloticus | Oreochromis_niloticus | GCF_001858045.2_O_niloticus_UMD_NMBU | train_val |
| Oryzias latipes | Oryzias_latipes | GCF_002234675.1_ASM223467v1 | train_val |
| Oryzias melastigma | Oryzias_melastigma | GCF_002922805.2_ASM292280v2 | train_val |
| Oxyura jamaicensis | Oxyura_jamaicensis | GCF_011077185.1_BPBGC_Ojam_1.0 | train_val |
| Pangasianodon hypophthalmus | Pangasianodon_hypophthalmus | GCF_009078355.1_GENO_Phyp_1.0 | train_val |
| Pantherophis guttatus | Pantherophis_guttatus | GCF_001185365.1_UNIGE_PanGut_3.0 | train_val |
| Paralichthys olivaceus | Paralichthys_olivaceus | GCF_001970005.1_Flounder_ref_guided_V1.0 | test |
| Parambassis ranga | Parambassis_ranga | GCF_900634625.1_fParRan2.1 | train_val |
| Paramormyrops kingsleyae | Paramormyrops_kingsleyae | GCF_002872115.1_PKINGS_0.1 | train_val |
| Parus major | Parus_major | GCF_001522545.3_Parus_major1.1 | train_val |
| Passer montanus | Passer_montanus | GCF_014805655.1_ASM1480565v1 | train_val |
| Pelecanus crispus | Pelecanus_crispus | GCF_000687375.1_ASM68737v1 | train_val |
| Pelodiscus sinensis | Pelodiscus_sinensis | GCF_000230535.1_PelSin_1.0 | train_val |
| Perca flavescens | Perca_flavescens | GCF_004354835.1_PFLA_1.0 | train_val |
| Perca fluviatilis | Perca_fluviatilis | GCF_010015445.1_GENO_Pfluv_1.0 | train_val |
| Periophthalmus magnuspinnatus | Periophthalmus_magnuspinnatus | GCF_009829125.1_fPerMag1.pri | train_val |
| Petromyzon marinus | Petromyzon_marinus | GCF_010993605.1_kPetMar1.pri | train_val |
| Phaethon lepturus | Phaethon_lepturus | GCF_000687285.1_ASM68728v1 | train_val |
| Phalacrocorax carbo | Phalacrocorax_carbo | GCF_000708925.1_ASM70892v1 | train_val |
| Phasianus colchicus | Phasianus_colchicus | GCF_004143745.1_ASM414374v1 | test |
| Pimephales promelas | Pimephales_promelas | GCF_016745375.1_EPA_FHM_2.0 | test |
| Pipra filicauda | Pipra_filicauda | GCF_003945595.2_ASM394559v2 | train_val |
| Plectropomus leopardus | Plectropomus_leopardus | GCF_008729295.1_YSFRI_Pleo_2.0 | test |
| Podarcis muralis | Podarcis_muralis | GCF_004329235.1_PodMur_1.0 | test |
| Poecilia formosa | Poecilia_formosa | GCF_000485575.1_Poecilia_formosa-5.1.2 | train_val |
| Poecilia latipinna | Poecilia_latipinna | GCF_001443285.1_P_latipinna-1.0 | train_val |
| Poecilia mexicana | Poecilia_mexicana | GCF_001443325.1_P_mexicana-1.0 | train_val |
| Poecilia reticulata | Poecilia_reticulata | GCF_000633615.1_Guppy_female_1.0_MT | train_val |
| Pogona vitticeps | Pogona_vitticeps | GCF_900067755.1_pvi1.1 | train_val |
| Polyodon spathula | Polyodon_spathula | GCF_017654505.1_ASM1765450v1 | test |
| Polypterus senegalus | Polypterus_senegalus | GCF_016835505.1_ASM1683550v1 | train_val |
| Protobothrops mucrosquamatus | Protobothrops_mucrosquamatus | GCF_001527695.2_P.Mucros_1.0 | train_val |
| Protopterus annectens | Protopterus_annectens | GCF_019279795.1_PAN1.0 | train_val |
| Pseudochaenichthys georgianus | Pseudochaenichthys_georgianus | GCF_902827115.1_fPseGeo1.1 | train_val |
| Pseudonaja textilis | Pseudonaja_textilis | GCF_900518735.1_EBS10Xv2-PRI | test (used) |
| Pseudopodoces humilis | Pseudopodoces_humilis | GCF_000331425.1_PseHum1.0 | train_val |
| Pterocles gutturalis | Pterocles_gutturalis | GCF_000699245.1_ASM69924v1 | train_val |
| Pundamilia nyererei | Pundamilia_nyererei | GCF_000239375.1_PunNye1.0 | train_val |
| Pungitius pungitius | Pungitius_pungitius | GCF_902500615.1_NSP_V7 | test |
| Puntigrus tetrazona | Puntigrus_tetrazona | GCF_018831695.1_ASM1883169v1 | train_val |
| Pygocentrus nattereri | Pygocentrus_nattereri | GCF_015220715.1_fPygNat1.pri | train_val |
| Pygoscelis adeliae | Pygoscelis_adeliae | GCF_000699105.1_ASM69910v1 | train_val |
| Pyrgilauda ruficollis | Pyrgilauda_ruficollis | GCF_017590135.1_ASM1759013v1 | train_val |
| Python bivittatus | Python_bivittatus | GCF_000186305.1_Python_molurus_bivittatus-5.0.2 | test |
| Rana temporaria | Rana_temporaria | GCF_905171775.1_aRanTem1.1 | train_val |
| Rhinatrema bivittatum | Rhinatrema_bivittatum | GCF_901001135.1_aRhiBiv1.1 | test |
| Rhincodon typus | Rhincodon_typus | GCF_001642345.1_ASM164234v2 | test (used) |
| Salarias fasciatus | Salarias_fasciatus | GCF_902148845.1_fSalaFa1.1 | train_val |
| Salmo salar | Salmo_salar | GCF_905237065.1_Ssal_v3.1 | train_val |
| Salmo trutta | Salmo_trutta | GCF_901001165.1_fSalTru1.1 | train_val |
| Salvelinus alpinus | Salvelinus_alpinus | GCF_002910315.2_ASM291031v2 | test |
| Salvelinus namaycush | Salvelinus_namaycush | GCF_016432855.1_SaNama_1.0 | train_val |
| Salvelinus sp IW2-2015 | Salvelinus_sp._IW2-2015 | GCF_002910315.2_ASM291031v2 | train_val |

| scientific | ID | version | split |
|---|---|---|---|
| Sander lucioperca | Sander_lucioperca | GCF_008315115.2_SLUC_FBN_1.2 | train_val |
| Scatophagus argus | Scatophagus_argus | GCF_020382885.2_fScaArg1.pri | test |
| Sceloporus undulatus | Sceloporus_undulatus | GCF_019175285.1_SceUnd_v1.1 | train_val |
| Scleropages formosus | Scleropages_formosus | GCF_900964775.1_fSclFor1.1 | train_val |
| Scophthalmus maximus | Scophthalmus_maximus | GCF_013347765.1_ASM1334776v1 | train_val |
| Scyliorhinus canicula | Scyliorhinus_canicula | GCF_902713615.1_sScyCan1.1 | train_val |
| Sebastes umbrosus | Sebastes_umbrosus | GCF_015220745.1_fSebUmb1.pri | train_val |
| Serinus canaria | Serinus_canaria | GCF_007115625.1_cibio_Scana_2019 | train_val |
| Seriola dumerili | Seriola_dumerili | GCF_002260705.1_Sdu_1.0 | train_val |
| Seriola lalandi | Seriola_lalandi | GCF_002814215.1_Sedor1 | train_val |
| Silurus meridionalis | Silurus_meridionalis | GCF_014805685.1_ASM1480568v1 | test |
| Simochromis diagramma | Simochromis_diagramma | GCF_900408965.1_fSimDia1.1 | train_val |
| Siniperca chuatsi | Siniperca_chuatsi | GCF_020085105.1_ASM2008510v1 | train_val |
| Sinocyclocheilus anshuiensis | Sinocyclocheilus_anshuiensis | GCF_001515605.1_SAMN03320099.WGS_v1.1 | test |
| Sinocyclocheilus grahami | Sinocyclocheilus_grahami | GCF_001515645.1_SAMN03320097.WGS_v1.1 | train_val |
| Sinocyclocheilus rhinocerous | Sinocyclocheilus_rhinocerous | GCF_001515625.1_SAMN03320098_v1.1 | train_val |
| Solea senegalensis | Solea_senegalensis | GCF_019176455.1_IFAPA_SoseM_1 | train_val |
| Sparus aurata | Sparus_aurata | GCF_900880675.1_fSpaAur1.1 | test (used) |
| Sphaeramia orbicularis | Sphaeramia_orbicularis | GCF_902148855.1_fSphaOr1.1 | test |
| Stegastes partitus | Stegastes_partitus | GCF_000690725.1_Stegastes_partitus-1.0.2 | train_val |
| Strigops habroptila | Strigops_habroptila | GCF_004027225.2_bStrHab1.2.pri | test |
| Struthio camelus | Struthio_camelus | GCF_000698965.1_ASM69896v1 | train_val |
| Sturnus vulgaris | Sturnus_vulgaris | GCF_001447265.1_Sturnus_vulgaris-1.0 | train_val |
| Syngnathus acus | Syngnathus_acus | GCF_901709675.1_fSynAcu1.2 | test |
| Tachysurus fulvidraco | Tachysurus_fulvidraco | GCF_003724035.1_ASM372403v1 | train_val |
| Taeniopygia guttata | Taeniopygia_guttata | GCF_003957565.2_bTaeGut1.4.pri | train_val |
| Takifugu rubripes | Takifugu_rubripes | GCF_901000725.2_fTakRub1.2 | train_val |
| Tauraco erythrolophus | Tauraco_erythrolophus | GCF_000709365.1_ASM70936v1 | train_val |
| Terrapene carolina | Terrapene_carolina | GCF_002925995.2_T_m_triunguis-2.0 | train_val |
| Thalassophryne amazonica | Thalassophryne_amazonica | GCF_902500255.1_fThaAma1.1 | train_val |
| Thamnophis elegans | Thamnophis_elegans | GCF_009769535.1_rThaEle1.pri | test |
| Thamnophis sirtalis | Thamnophis_sirtalis | GCF_001077635.1_Thamnophis_sirtalis-6.0 | test |
| Thunnus albacares | Thunnus_albacares | GCF_914725855.1_fThuAlb1.1 | train_val |
| Thunnus maccoyii | Thunnus_maccoyii | GCF_910596095.1_fThuMac1.1 | train_val |
| Tinamus guttatus | Tinamus_guttatus | GCF_000705375.1_ASM70537v2 | train_val |
| Toxotes jaculatrix | Toxotes_jaculatrix | GCF_017976425.1_fToxJac2.pri | train_val |
| Trachemys scripta | Trachemys_scripta | GCF_013100865.1_CAS_Tse_1.0 | test |
| Trematomus bernacchii | Trematomus_bernacchii | GCF_902827165.1_fTreBer1.1 | train_val |
| Tyto alba | Tyto_alba | GCF_018691265.1_T.alba_DEE_v4.0 | test |
| Varanus komodoensis | Varanus_komodoensis | GCF_004798865.1_ASM479886v1 | train_val |
| Xenopus laevis | Xenopus_laevis | GCF_017654675.1_Xenopus_laevis_v10.1 | train_val |
| Xenopus tropicalis | Xenopus_tropicalis | GCF_000004195.4_UCB_Xtro_10.0 | train_val |
| Xiphias gladius | Xiphias_gladius | GCF_016859285.1_ASM1685928v1 | test (used) |
| Xiphophorus couchianus | Xiphophorus_couchianus | GCF_001444195.1_X_couchianus-1.0 | train_val |
| Xiphophorus hellerii | Xiphophorus_hellerii | GCF_003331165.1_Xiphophorus_hellerii-4.1 | test |
| Xiphophorus maculatus | Xiphophorus_maculatus | GCF_002775205.1_X_maculatus-5.0-male | train_val |
| Zonotrichia albicollis | Zonotrichia_albicollis | GCF_000385455.1_Zonotrichia_albicollis-1.0.1 | train_val |
| Zootoca vivipara | Zootoca_vivipara | GCF_011800845.1_UG_Zviv_1 | test |

##### S2.1.4 Invertebrates

Invertebrate training, validation and test genomes were acquired from RefSeq on May 6th, 2022.

Test genomes were assigned randomly *within* genomes that had a matching or close AUGUSTUS trained model available and *within* those that did not.

All species assigned to test that had a close AUGUSTUS model were actually used as test species in this manuscript, additional used test species were selected arbitrarily (every Nth).

Table S4: All invertebrate genomes, set assignment and versions

| scientific | ID | version | split |
|---|---|---|---|
| Acanthaster planci | Acanthaster_planci | GCF_001949145.1_OKI-Apl_1.0 | train_val |
| Acromyrmex echinatior | Acromyrmex_echinatior | GCF_000204515.1_Aech_3.9 | train_val |
| Acropora digitifera | Acropora_digitifera | GCF_000222465.1_Adig_1.1 | test |
| Acropora millepora | Acropora_millepora | GCF_013753865.1_Amil_v2.1 | train_val |
| Actinia tenebrosa | Actinia_tenebrosa | GCF_009602425.1_ASM960242v1 | train_val |
| Acyrthosiphon pisum | Acyrthosiphon_pisum | GCF_005508785.1_pea_aphid_22Mar2018_4r6ur | train_val |
| Aedes aegypti | Aedes_aegypti | GCF_002204515.2_AaegL5.0 | train_val |
| Aedes albopictus | Aedes_albopictus | GCF_006496715.1_Aalbo_primary.1 | test |
| Aethina tumida | Aethina_tumida | GCF_001937115.1_Atum_1.0 | train_val |
| Agrilus planipennis | Agrilus_planipennis | GCF_000699045.2_Apla_2.0 | train_val |
| Amphibalanus amphitrite | Amphibalanus_amphitrite | GCF_019059575.1_NRLGWU_Aamphi_draft | train_val |
| Amphimedon queenslandica | Amphimedon_queenslandica | GCF_000090795.1_v1.0 | train_val |
| Amyelois transitella | Amyelois_transitella | GCF_001186105.1_ASM118610v1 | train_val |
| Anneissia japonica | Anneissia_japonica | GCF_011630105.1_ASM1163010v1 | test |
| Anopheles albimanus | Anopheles_albimanus | GCF_013758885.1_VT_AalbS3_pri_1.0 | train_val |
| Anopheles arabiensis | Anopheles_arabiensis | GCF_016920715.1_AaraD3 | train_val |
| Anopheles coluzzii | Anopheles_coluzzii | GCF_016920705.1_AcolMOP1 | train_val |
| Anopheles gambiae | Anopheles_gambiae | GCF_000005575.2_AgamP3 | test |
| Anopheles merus | Anopheles_merus | GCF_017562075.2_AmerM5.1 | train_val |
| Anopheles stephensi | Anopheles_stephensi | GCF_013141755.1_UCI_ANSTEP_V1.0 | train_val |
| Anoplophora glabripennis | Anoplophora_glabripennis | GCF_000390285.2_Agla_2.0 | train_val |
| Aphidius gifuensis | Aphidius_gifuensis | GCF_014905175.1_ASM1490517v1 | train_val |
| Aphis gossypii | Aphis_gossypii | GCF_004010815.1_ASM401081v1 | train_val |
| Apis cerana | Apis_cerana | GCF_001442555.1_ACSNU-2.0 | train_val |
| Apis dorsata | Apis_dorsata | GCF_000469605.1_Apis_dorsata_1.3 | test (used) |
| Apis florea | Apis_florea | GCF_000184785.3_Aflo_1.1 | train_val |
| Apis laboriosa | Apis_laboriosa | GCF_014066325.1_ASM1406632v1 | train_val |
| Apis mellifera | Apis_mellifera | GCF_003254395.2_Amel_HAv3.1 | train_val |
| Aplysia californica | Aplysia_californica | GCF_000002075.1_AplCal3.0 | train_val |
| Aricia agestis | Aricia_agestis | GCF_905147365.1_ilAriAges1.1 | test |
| Asterias rubens | Asterias_rubens | GCF_902459465.1_eAstRub1.3 | train_val |
| Athalia rosae | Athalia_rosae | GCF_000344095.2_Aros_2.0 | test |
| Atta cephalotes | Atta_cephalotes | GCF_000143395.1_Attacep1.0 | test (used) |
| Atta colombica | Atta_colombica | GCF_001594045.1_Acol1.0 | train_val |
| Bactrocera dorsalis | Bactrocera_dorsalis | GCF_000789215.1_ASM78921v2 | test |
| Bactrocera latifrons | Bactrocera_latifrons | GCF_001853355.1_ASM185335v1 | test |
| Bactrocera oleae | Bactrocera_oleae | GCF_001188975.3_MU_Boleae_v2 | test |
| Bactrocera tryoni | Bactrocera_tryoni | GCF_016617805.1_CSIRO_BtryS06_freeze2 | train_val |
| Belonocnema kinseyi | Belonocnema_kinseyi | GCF_010883055.1_B_treatae_v1 | train_val |
| Belonocnema treatae | Belonocnema_treatae | GCF_010883055.1_B_treatae_v1 | train_val |
| Bemisia tabaci | Bemisia_tabaci | GCF_001854935.1_ASM185493v1 | train_val |
| Bicyclus anynana | Bicyclus_anynana | GCF_900239965.1_Bicyclus_anynana_v1.2 | train_val |
| Biomphalaria glabrata | Biomphalaria_glabrata | GCF_000457365.1_ASM45736v1 | train_val |
| Bombus bifarius | Bombus_bifarius | GCF_011952205.1_Bbif_JDL3187 | train_val |
| Bombus impatiens | Bombus_impatiens | GCF_000188095.3_BIMP_2.2 | test (used) |
| Bombus pyrosoma | Bombus_pyrosoma | GCF_014825855.1_ASM1482585v1 | train_val |
| Bombus terrestris | Bombus_terrestris | GCF_000214255.1_Bter_1.0 | train_val |
| Bombus vancouverensis | Bombus_vancouverensis | GCF_011952275.1_Bvanc_JDL1245 | train_val |
| Bombus vosnesenskii | Bombus_vosnesenskii | GCF_011952255.1_Bvos_JDL3184-5_v1.1 | test |
| Bombyx mandarina | Bombyx_mandarina | GCF_003987935.1_ASM398793v1 | train_val |
| Bombyx mori | Bombyx_mori | GCF_014905235.1_Bmori_2016v1.0 | train_val |
| Bradysia coprophila | Bradysia_coprophila | GCF_014529535.1_BU_Bcop_v1 | train_val |
| Branchiostoma belcheri | Branchiostoma_belcheri | GCF_001625305.1_Haploidv18h27 | test |
| Branchiostoma floridae | Branchiostoma_floridae | GCF_000003815.2_Bfl_VNyyK | test |
| Brugia malayi | Brugia_malayi | GCF_000002995.4_B_malayi-4.0 | train_val |
| busco downloads | busco_downloads | NaN | test |
| Caenorhabditis briggsae | Caenorhabditis_briggsae | GCF_000004555.2_CB4 | train_val |
| Caenorhabditis elegans | Caenorhabditis_elegans | GCF_000002985.6_WBcel235 | train_val |
| Caenorhabditis remanei | Caenorhabditis_remanei | GCF_000149515.1_ASM14951v1 | test (used) |
| Camponotus floridanus | Camponotus_floridanus | GCF_003227725.1_Cflo_v7.5 | train_val |
| Capsaspora owczarzaki | Capsaspora_owczarzaki | GCF_000151315.2_C_owczarzaki_V2 | train_val |
| Centruroides sculpturatus | Centruroides_sculpturatus | GCF_000671375.1_Cexi_2.0 | train_val |
| Cephus cinctus | Cephus_cinctus | GCF_000341935.1_Ccin1 | test |
| Ceratina calcarata | Ceratina_calcarata | GCF_001652005.1_ASM165200v1 | test |
| Ceratitis capitata | Ceratitis_capitata | GCF_000347755.3_Ccap_2.1 | train_val |
| Ceratosolen solmsi | Ceratosolen_solmsi | GCF_000503995.1_CerSol_1.0 | test |
| Chelonus insularis | Chelonus_insularis | GCF_013357705.1_ASM1335770v1 | train_val |
| Chrysoperla carnea | Chrysoperla_carnea | GCF_905475395.1_inChrCarn1.1 | test |
| Cimex lectularius | Cimex_lectularius | GCF_000648675.2_Clec_2.1 | train_val |
| Ciona intestinalis | Ciona_intestinalis | GCF_000224145.3_KH | train_val |
| Coccinella septempunctata | Coccinella_septempunctata | GCF_907165205.1_icCocSept1.1 | train_val |
| Colias croceus | Colias_croceus | GCF_905220415.1_ilColCroc2.1 | test |
| Colletes gigas | Colletes_gigas | GCF_013123115.1_ASM1312311v1 | train_val |
| Contarinia nasturtii | Contarinia_nasturtii | GCF_009176525.2_AAFC_CNas_1.1 | train_val |
| Copidosoma floridanum | Copidosoma_floridanum | GCF_000648655.2_Cflo_2.0 | test |
| Cotesia glomerata | Cotesia_glomerata | GCF_020080835.1_MPM_Cglom_v2.3 | train_val |
| Crassostrea gigas | Crassostrea_gigas | GCF_902806645.1_cgigas_uk_roslin_v1 | train_val |
| Crassostrea virginica | Crassostrea_virginica | GCF_002022765.2_C_virginica-3.0 | test |
| Cryptotermes secundus | Cryptotermes_secundus | GCF_002891405.2_Csec_1.0 | train_val |
| Ctenocephalides felis | Ctenocephalides_felis | GCF_003426905.1_ASM342690v1 | test (used) |

| scientific | ID | version | split |
|---|---|---|---|
| Culex pipiens | Culex_pipiens | GCF_016801865.1_TS_Cpip_V1 | test (used) |
| Culex quinquefasciatus | Culex_quinquefasciatus | GCF_015732765.1_VPISU_Cqui_1.0_pri_paternal | test |
| Cyphomyrmex costatus | Cyphomyrmex_costatus | GCF_001594065.1_Ccosl1.0 | test |
| Danaus plexippus | Danaus_plexippus | GCF_009731565.1_Dplex_v4 | test |
| Daphnia magna | Daphnia_magna | GCF_020631705.1_ASM2063170v1.1 | train_val |
| Daphnia pulex | Daphnia_pulex | GCF_021134715.1_ASM2113471v1 | test |
| Daphnia pulicaria | Daphnia_pulicaria | GCF_021234035.1_SC_F0-13Bv2 | train_val |
| Dendroctonus ponderosae | Dendroctonus_ponderosae | GCF_000355655.1_DendPond_male_1.0 | train_val |
| Dendronephthya gigantea | Dendronephthya_gigantea | GCF_004324835.1_DenGig_1.0 | train_val |
| Dermacentor silvarum | Dermacentor_silvarum | GCF_013339745.1_ASM1333974v1 | train_val |
| Dermatophagoides pteronyssinus | Dermatophagoides_pteronyssinus | GCF_001901225.1_ASM190122v2 | train_val |
| Diabrotica virgifera | Diabrotica_virgifera | GCF_003013835.1_Dvir_v2.0 | train_val |
| Diachasma alloeum | Diachasma_alloeum | GCF_001412515.2_Dall2.0 | train_val |
| Diaphorina citri | Diaphorina_citri | GCF_000475195.1_Diaci_psyllid_genome_assembly_v... | train_val |
| Dinoponera quadriceps | Dinoponera_quadriceps | GCF_001313825.1_ASM131382v1 | test |
| Diprion similis | Diprion_similis | GCF_021155765.1_iyDipSimi1.1 | test |
| Diuraphis noxia | Diuraphis_noxia | GCF_001186385.1_Dnoxia_1.0 | train_val |
| Drosophila albomicans | Drosophila_albomicans | GCF_009650485.1_drosAlbom15112-1751.03v1 | test (used) |
| Drosophila ananassae | Drosophila_ananassae | GCF_017639315.1_ASM1763931v2 | train_val |
| Drosophila arizonae | Drosophila_arizonae | GCF_001654025.1_ASM165402v1 | test |
| Drosophila biarmipes | Drosophila_biarmipes | GCF_018148935.1_ASM1814893v1 | train_val |
| Drosophila bipectinata | Drosophila_bipectinata | GCF_018153845.1_ASM1815384v1 | train_val |
| Drosophila busckii | Drosophila_busckii | GCF_011750605.1_ASM1175060v1 | train_val |
| Drosophila elegans | Drosophila_elegans | GCF_018152505.1_ASM1815250v1 | train_val |
| Drosophila erecta | Drosophila_erecta | GCF_003286155.1_DereRS2 | train_val |
| Drosophila eugracilis | Drosophila_eugracilis | GCF_018153835.1_ASM1815383v1 | train_val |
| Drosophila ficusphila | Drosophila_ficusphila | GCF_018152265.1_ASM1815226v1 | test |
| Drosophila grimshawi | Drosophila_grimshawi | GCF_018153295.1_ASM1815329v1 | train_val |
| Drosophila guanche | Drosophila_guanche | GCF_900245975.1_DGUA_6 | test |
| Drosophila hydei | Drosophila_hydei | GCF_003285905.1_DhydRS2 | train_val |
| Drosophila innubila | Drosophila_innubila | GCF_004354385.1_UK_Dinn_1.0 | train_val |
| Drosophila kikkawai | Drosophila_kikkawai | GCF_018152535.1_ASM1815253v1 | train_val |
| Drosophila mauritiana | Drosophila_mauritiana | GCF_004382145.1_ASM438214v1 | train_val |
| Drosophila melanogaster | Drosophila_melanogaster | GCF_000001215.4_Release_6_plus_ISO1_MT | train_val |
| Drosophila miranda | Drosophila_miranda | GCF_003369915.1_D.miranda_PacBio2.1 | train_val |
| Drosophila mojavensis | Drosophila_mojavensis | GCF_018153725.1_ASM1815372v1 | train_val |
| Drosophila navojoa | Drosophila_navojoa | GCF_001654015.2_UFRJ_Dnav_4.2 | train_val |
| Drosophila novamexicana | Drosophila_novamexicana | GCF_003285875.2_DnovRS2.1 | train_val |
| Drosophila obscura | Drosophila_obscura | GCF_018151105.1_ASM1815110v1 | test |
| Drosophila persimilis | Drosophila_persimilis | GCF_003286085.1_DperRS2 | train_val |
| Drosophila pseudoobscura | Drosophila_pseudoobscura | GCF_009870125.1_UCI_Dpse_MV25 | train_val |
| Drosophila rhopaloa | Drosophila_rhopaloa | GCF_018152115.1_ASM1815211v1 | train_val |
| Drosophila santomea | Drosophila_santomea | GCF_016746245.2_Prin_Dsan_1.1 | train_val |
| Drosophila sechellia | Drosophila_sechellia | GCF_004382195.1_ASM438219v1 | test |
| Drosophila serrata | Drosophila_serrata | GCF_002093755.1_Dser1.0 | train_val |
| Drosophila simulans | Drosophila_simulans | GCF_016746395.2_Prin_Dsim_3.1 | train_val |
| Drosophila subobscura | Drosophila_subobscura | GCF_008121235.1_UCBerk_Dsub_1.0 | train_val |
| Drosophila subpulchrella | Drosophila_subpulchrella | GCF_014743375.2_RU_Dsub_v1.1 | test |
| Drosophila suzukii | Drosophila_suzukii | GCF_013340165.1_LBDM_Dsuz_2.1.pri | train_val |
| Drosophila takahashii | Drosophila_takahashii | GCF_018152695.1_ASM1815269v1 | test |
| Drosophila teissieri | Drosophila_teissieri | GCF_016746235.2_Prin_Dtei_1.1 | train_val |
| Drosophila virilis | Drosophila_virilis | GCF_003285735.1_DvirRS2 | test (used) |
| Drosophila willistoni | Drosophila_willistoni | GCF_000005925.1_dwil_caf1 | test |
| Drosophila yakuba | Drosophila_yakuba | GCF_016746365.2_Prin_Dyak_Tai18E2_2.1 | test |
| Dufourea novaeangliae | Dufourea_novaeangliae | GCF_001272555.1_ASM127255v1 | test |
| Echinococcus granulosus | Echinococcus_granulosus | GCF_000524195.1_ASM52419v1 | test |
| Eufriesea mexicana | Eufriesea_mexicana | GCF_001483705.1_ASM148370v1 | test |
| Eurytemora affinis | Eurytemora_affinis | GCF_000591075.1_Eaff_2.0 | test |
| Exaiptasia diaphana | Exaiptasia_diaphana | GCF_001417965.1_Aiptasia_genome_1.1 | train_val |
| Folsomia candida | Folsomia_candida | GCF_002217175.1_ASM221717v1 | train_val |
| Fonticula alba | Fonticula_alba | GCF_000388065.1_Font_alba_ATCC_38817_V2 | train_val |
| Fopius arisanus | Fopius_arisanus | GCF_000806365.1_ASM80636v1 | train_val |
| Formica exsecta | Formica_exsecta | GCF_003651465.1_ASM365146v1 | test |
| Frankliniella occidentalis | Frankliniella_occidentalis | GCF_000697945.2_Focc_2.1 | test (used) |
| Frieseomelitta varia | Frieseomelitta_varia | GCF_011392965.1_Fvar_1.2 | test |
| Galendromus occidentalis | Galendromus_occidentalis | GCF_000255335.1_Mocc_1.0 | train_val |
| Galleria mellonella | Galleria_mellonella | GCF_003640425.2_ASM364042v2 | train_val |
| Gigantopelta aegis | Gigantopelta_aegis | GCF_016097555.1_Gae_host_genome | train_val |
| Glossina fuscipes | Glossina_fuscipes | GCF_014805625.1_Yale_Gfus_2 | train_val |
| Habropoda laboriosa | Habropoda_laboriosa | GCF_001263275.1_ASM126327v1 | train_val |
| Haliotis rubra | Haliotis_rubra | GCF_003918875.1_ASM391887v1 | train_val |
| Haliotis rufescens | Haliotis_rufescens | GCF_003343065.1_H.ruf_v1.0 | test |
| Halyomorpha halys | Halyomorpha_halys | GCF_000696795.2_Hhal_2.0 | train_val |
| Harmonia axyridis | Harmonia_axyridis | GCF_914767665.1_icHarAxyr1.1 | test |
| Harpegnathos saltator | Harpegnathos_saltator | GCF_003227715.1_Hsal_v8.5 | train_val |
| Helicoverpa armigera | Helicoverpa_armigera | GCF_002156985.1_Harm_1.0 | test |
| Helobdella robusta | Helobdella_robusta | GCF_000326865.1_Helobdella_robusta_v1.0 | test |
| Hermetia illucens | Hermetia_illucens | GCF_905115235.1_iHerIll2.2.curated.20191125 | train_val |
| Homalodisca vitripennis | Homalodisca_vitripennis | GCF_021130785.1_UT_GWSS_2.1 | train_val |
| Homarus americanus | Homarus_americanus | GCF_018991925.1_GMGI_Hamer_2.0 | train_val |

| scientific | ID | version | split |
|---|---|---|---|
| Hyalella azteca | Hyalella_azteca | GCF_000764305.1_Hazt_2.0 | test |
| Hydra vulgaris | Hydra_vulgaris | GCF_000004095.1_Hydra_RP_1.0 | test |
| Hyposmocoma kahamanoa | Hyposmocoma_kahamanoa | GCF_003589595.1_ASM358959v1 | test (used) |
| Ischnura elegans | Ischnura_elegans | GCF_921293095.1_ioIscEleg1.1 | train_val |
| Ixodes scapularis | Ixodes_scapularis | GCF_016920785.2_ASM1692078v2 | train_val |
| Lepeophtheirus salmonis | Lepeophtheirus_salmonis | GCF_016086655.3_UVic_Lsal_1.2 | train_val |
| Leptinotarsa decemlineata | Leptinotarsa_decemlineata | GCF_000500325.1_Ldec_2.0 | train_val |
| Leptopilina heterotoma | Leptopilina_heterotoma | GCF_015476425.1_ASM1547642v1 | train_val |
| Limulus polyphemus | Limulus_polyphemus | GCF_000517525.1_Limulus_polyphemus-2.1.2 | test |
| Linepithema humile | Linepithema_humile | GCF_000217595.1_Lhum_UMD_V04 | train_val |
| Lingula anatina | Lingula_anatina | GCF_001039355.2_LinAna2.0 | train_val |
| Loa loa | Loa_loa | GCF_000183805.2_Loa_loa_V3.1 | train_val |
| Lottia gigantea | Lottia_gigantea | GCF_000327385.1_Helro1 | train_val |
| Lucilia cuprina | Lucilia_cuprina | GCF_000699065.1_Lcup_2.0 | test |
| Lucilia sericata | Lucilia_sericata | GCF_015586225.1_ASM1558622v1 | train_val |
| Lytechinus variegatus | Lytechinus_variegatus | GCF_018143015.1_Lvar_3.0 | test |
| Manduca sexta | Manduca_sexta | GCF_014839805.1_JHU_Msex_v1.0 | train_val |
| Maniola hyperantus | Maniola_hyperantus | GCF_902806685.1_iAphHyp1.1 | test |
| Maniola jurtina | Maniola_jurtina | GCF_905333055.1_ilManJurt1.1 | train_val |
| Megachile rotundata | Megachile_rotundata | GCF_000220905.1_MROT_1.0 | train_val |
| Megalopta genalis | Megalopta_genalis | GCF_011865705.1_USU_MGEN_1.2 | train_val |
| Melanaphis sacchari | Melanaphis_sacchari | GCF_002803265.2_SCAv2.0 | train_val |
| Melitaea cinxia | Melitaea_cinxia | GCF_905220565.1_ilMelCinx1.1 | train_val |
| Mercenaria mercenaria | Mercenaria_mercenaria | GCF_014805675.1_ASM1480567v1.1 | train_val |
| Microplitis demolitor | Microplitis_demolitor | GCF_000572035.2_Mdem2 | train_val |
| Mizuhopecten yessoensis | Mizuhopecten_yessoensis | GCF_002113885.1_ASM211388v2 | test |
| Monomorium pharaonis | Monomorium_pharaonis | GCF_013373865.1_ASM1337386v2 | test |
| Monosiga brevicollis | Monosiga_brevicollis | GCF_000002865.3_V1.0 | train_val |
| Musca domestica | Musca_domestica | GCF_000371365.1_Musca_domestica-2.0.2 | train_val |
| Myzus persicae | Myzus_persicae | GCF_001856785.1_MPER_G0061.0 | train_val |
| Nasonia vitripennis | Nasonia_vitripennis | GCF_009193385.2_Nvit_psr_1.1 | train_val |
| Necator americanus | Necator_americanus | GCF_000507365.1_N_americanus_v1 | train_val |
| Nematostella vectensis | Nematostella_vectensis | GCF_000209225.1_ASM20922v1 | train_val |
| Neodiprion fabricii | Neodiprion_fabricii | GCF_021155785.1_iyNeoFabr1.1 | train_val |
| Neodiprion lecontei | Neodiprion_lecontei | GCF_021901455.1_iyNeoLeco1.1 | train_val |
| Neodiprion pinetum | Neodiprion_pinetum | GCF_021155775.1_iyNeoPine1.1 | train_val |
| Neodiprion virginiana | Neodiprion_virginiana | GCF_021901495.1_iyNeoVirg1.1 | train_val |
| Nicrophorus vespilloides | Nicrophorus_vespilloides | GCF_001412225.1_Nicve_v1.0 | train_val |
| Nilaparvata lugens | Nilaparvata_lugens | GCF_014356525.1_ASM1435652v1 | train_val |
| Nomia melanderi | Nomia_melanderi | GCF_003710045.1_USU_Nmel_1.2 | train_val |
| Nylanderia fulva | Nylanderia_fulva | GCF_005281655.1_TAMU_Nfulva_1.0 | train_val |
| Octopus bimaculoides | Octopus_bimaculoides | GCF_001194135.1_Octopus_bimaculoides_v2_0 | train_val |
| Octopus sinensis | Octopus_sinensis | GCF_006345805.1_ASM634580v1 | train_val |
| Octopus vulgaris | Octopus_vulgaris | GCF_006345805.1_ASM634580v1 | train_val |
| Odontomachus brunneus | Odontomachus_brunneus | GCF_010583005.1_Obru_v1 | train_val |
| Onthophagus taurus | Onthophagus_taurus | GCF_000648695.1_Otau_2.0 | test |
| Ooceraea biroi | Ooceraea_biroi | GCF_003672135.1_Obir_v5.4 | train_val |
| Opisthorchis viverrini | Opisthorchis_viverrini | GCF_000715545.1_OpiViv1.0 | test (used) |
| Orbicella faveolata | Orbicella_faveolata | GCF_002042975.1_ofav_dov_v1 | train_val |
| Orussus abietinus | Orussus_abietinus | GCF_000612105.2_Oabi_2.0 | train_val |
| Osmia bicornis | Osmia_bicornis | GCF_907164935.1_iOsmBic2.1 | train_val |
| Osmia lignaria | Osmia_lignaria | GCF_012274295.1_USDA_OLig_1.0 | train_val |
| Ostrinia furnacalis | Ostrinia_furnacalis | GCF_004193835.1_ASM419383v1 | train_val |
| Papilio machaon | Papilio_machaon | GCF_912999745.1_ilPapMach1.1 | train_val |
| Papilio polytes | Papilio_polytes | GCF_000836215.1_Ppol_1.0 | train_val |
| Papilio xuthus | Papilio_xuthus | GCF_000836235.1_Pxut_1.0 | train_val |
| Pararge aegeria | Pararge_aegeria | GCF_905163445.1_ilParAegt1.1 | train_val |
| Parasteatoda tepidariorum | Parasteatoda_tepidariorum | GCF_000365465.3_Ptep_3.0 | train_val |
| Patiria miniata | Patiria_miniata | GCF_015706575.1_ASM1570657v1 | train_val |
| Pecten maximus | Pecten_maximus | GCF_902652985.1_xPecMax1.1 | train_val |
| Pediculus humanus | Pediculus_humanus | GCF_000006295.1_JCVI_LOUSE_1.0 | test |
| Penaeus japonicus | Penaeus_japonicus | GCF_017312705.1_Mj_TUMSAT_v1.0 | train_val |
| Penaeus monodon | Penaeus_monodon | GCF_015228065.1_NSTDA_Pmon_1 | train_val |
| Penaeus vannamei | Penaeus_vannamei | GCF_003789085.1_ASM378908v1 | test |
| Photinus pyralis | Photinus_pyralis | GCF_008802855.1_Ppyr1.3 | train_val |
| Pieris brassicae | Pieris_brassicae | GCF_905147105.1_ilPieBrab1.1 | train_val |
| Pieris rapae | Pieris_rapae | GCF_905147795.1_ilPieRapa1.1 | train_val |
| Plutella xylostella | Plutella_xylostella | GCF_905116875.1_Haplomerged_assembly | train_val |
| Pocillopora damicornis | Pocillopora_damicornis | GCF_003704095.1_ASM370409v1 | train_val |
| Pogonomyrmex barbatus | Pogonomyrmex_barbatus | GCF_000187915.1_Pbar_UMD_V03 | train_val |
| Polistes canadensis | Polistes_canadensis | GCF_001313835.1_ASM131383v1 | train_val |
| Polistes dominula | Polistes_dominula | GCF_001465965.1_Pdom_r1.2 | test |
| Polistes fuscatus | Polistes_fuscatus | GCF_010416935.1_CU_Pfus_HIC | test |
| Pollicipes pollicipes | Pollicipes_pollicipes | GCF_011947565.2_Ppol_2 | train_val |
| Pomacea canaliculata | Pomacea_canaliculata | GCF_003073045.1_ASM307304v1 | train_val |
| Portunus trituberculatus | Portunus_trituberculatus | GCF_017591435.1_ASM1759143v1 | train_val |
| Priapulus caudatus | Priapulus_caudatus | GCF_000485595.1_Priapulus_caudatus-5.0.1 | test |
| Procambarus clarkii | Procambarus_clarkii | GCF_020424385.1_ASM2042438v2 | test |
| Pseudomyrmex gracilis | Pseudomyrmex_gracilis | GCF_002006095.1_ASM200609v1 | train_val |
| Rhagoletis pomonella | Rhagoletis_pomonella | GCF_013731165.1_Rhpom_1.0 | test |

| scientific | ID | version | split |
|---|---|---|---|
| Rhagoletis zephyria | Rhagoletis_zephyria | GCF_001687245.1_Rhagoletis_zephyria_1.0 | train_val |
| Rhipicephalus microplus | Rhipicephalus_microplus | GCF_013339725.1_ASM1333972v1 | train_val |
| Rhipicephalus sanguineus | Rhipicephalus_sanguineus | GCF_013339695.1_ASM1333969v1 | train_val |
| Rhopalosiphum maidis | Rhopalosiphum_maidis | GCF_003676215.2_ASM367621v3 | test (used) |
| Saccoglossus kowalevskii | Saccoglossus_kowalevskii | GCF_000003605.2_Skow_1.1 | train_val |
| Salpingoeca rosetta | Salpingoeca_rosetta | GCF_000188695.1_Proterospongia_sp_ATCC50818 | train_val |
| Scaptodrosophila lebanonensis | Scaptodrosophila_lebanonensis | GCF_003285725.1_SlebRS2 | train_val |
| Schistosoma haematobium | Schistosoma_haematobium | GCF_000699445.2_SchHae_2.0 | train_val |
| Schistosoma mansoni | Schistosoma_mansoni | GCF_000237925.1_ASM23792v2 | test (used) |
| Sipha flava | Sipha_flava | GCF_003268045.1_YSA_version1 | test |
| Sitophilus oryzae | Sitophilus_oryzae | GCF_002938485.1_Soryzae_2.0 | train_val |
| Solenopsis invicta | Solenopsis_invicta | GCF_016802725.1_UNIL_Sinv_3.0 | train_val |
| Sphaeroforma arctica | Sphaeroforma_arctica | GCF_001186125.1_Spha_arctica_JP610_V1 | train_val |
| Spodoptera frugiperda | Spodoptera_frugiperda | GCF_011064685.1_ZJU_Sfru_1.0 | train_val |
| Spodoptera litura | Spodoptera_litura | GCF_002706865.1_ASM270686v1 | train_val |
| Stegodyphus dumicola | Stegodyphus_dumicola | GCF_010614865.1_ASM1061486v1 | train_val |
| Stomoxys calcitrans | Stomoxys_calcitrans | GCF_001015335.1_Stomoxys_calcitrans-1.0.1 | train_val |
| Strongylocentrotus purpuratus | Strongylocentrotus_purpuratus | GCF_000002235.5_Spur_5.0 | train_val |
| Strongyloides ratti | Strongyloides_ratti | GCF_001040885.1_S_ratti_ED321 | test |
| Styela clava | Styela_clava | GCF_013122585.1_ASM1312258v2 | test |
| Stylophora pistillata | Stylophora_pistillata | GCF_002571385.1_Stylophora_pistillata_v1 | train_val |
| Teleopsis dalmanni | Teleopsis_dalmanni | GCF_002237135.1_ASM223713v2 | test |
| Temnothorax curvispinosus | Temnothorax_curvispinosus | GCF_003070985.1_ASM307098v1 | test |
| Tetranychus urticae | Tetranychus_urticae | GCF_000239435.1_ASM23943v1 | train_val |
| Thrips palmi | Thrips_palmi | GCF_012932325.1_TpBJ-2018v1 | train_val |
| Trachymyrmex cornetzi | Trachymyrmex_cornetzi | GCF_001594075.1_Tcor1.0 | test (used) |
| Trachymyrmex septentrionalis | Trachymyrmex_septentrionalis | GCF_001594115.1_Tsep1.0 | train_val |
| Trachymyrmex zeteki | Trachymyrmex_zeteki | GCF_001594055.1_Tzet1.0 | test |
| Tribolium castaneum | Tribolium_castaneum | GCF_000002335.3_Tcas5.2 | test (used) |
| Tribolium madens | Tribolium_madens | GCF_015345945.1_Tmad_KSU_1.1 | train_val |
| Trichinella spiralis | Trichinella_spiralis | GCF_000181795.1_Trichinella_spiralis-3.7.1 | train_val |
| Trichogramma pretiosum | Trichogramma_pretiosum | GCF_000599845.2_Tpre_2.0 | test |
| Trichoplax adhaerens | Trichoplax_adhaerens | GCF_000150275.1_v1.0 | train_val |
| Trichoplusia ni | Trichoplusia_ni | GCF_003590095.1_tn1 | test |
| Vanessa tameamea | Vanessa_tameamea | GCF_002938995.1_ASM293899v1 | test |
| Varroa destructor | Varroa_destructor | GCF_002443255.1_Vdes_3.0 | train_val |
| Varroa jacobsoni | Varroa_jacobsoni | GCF_002532875.1_vjacob_1.0 | train_val |
| Venturia canescens | Venturia_canescens | GCF_019457755.1_ASM1945775v1 | train_val |
| Vespa crabro | Vespa_crabro | GCF_910589235.1_iyVesCrab1.2 | train_val |
| Vespa mandarinia | Vespa_mandarinia | GCF_014083535.2_V.mandarinia_Nanaimo_p1.0 | train_val |
| Vespula pensylvanica | Vespula_pensylvanica | GCF_014466175.1_ASM1446617v1 | train_val |
| Vollenhovia emeryi | Vollenhovia_emeryi | GCF_000949405.1_V.emery_V1.0 | train_val |
| Wasmannia auropunctata | Wasmannia_auropunctata | GCF_000956235.1_wasmannia.A_1.0 | test |
| Xenia sp Carnegie-2017 | Xenia_sp._Carnegie-2017 | GCF_021976095.1_XeniaSp_v1 | train_val |
| Zerene cesonia | Zerene_cesonia | GCF_012273895.1_Zerene_cesonia_1.1 | train_val |
| Zeugodacus cucurbitae | Zeugodacus_cucurbitae | GCF_000806345.1_ASM80634v1 | train_val |
| Zootermopsis nevadensis | Zootermopsis_nevadensis | GCF_000696155.1_ZooNev1.0 | train_val |

#### S2.2 Training species sets

Table S5: Fungi species in datasets used for training

| name | seed_005_059 | seed_006_059 | seed_005_033 | Seed_006_033 |
|---|---|---|---|---|
| comment | automated selection | automated selection | automated selection | automated selection |
| trainers | Agaricus_bisporus | Aaosphaeria_arxii | Agaricus_bisporus | Aaosphaeria_arxii |
| | Alternaria_arborescens | Alternaria_arborescens | Alternaria_arborescens | Alternaria_arborescens |
| | Alternaria_burnsii | Alternaria_atra | Amorphotheca_resinae | Alternaria_atra |
| | Amorphotheca_resinae | Alternaria_burnsii | Apiotrichum_porosum | Alternaria_rosae |
| | Apiotrichum_porosum | Alternaria_rosae | Ascoidea_rubescens | Apiotrichum_porosum |
| | Aspergillus_alliaceus | Apiotrichum_porosum | Aspergillus_aculeatus | Ascoidea_rubescens |
| | Aspergillus_campestris | Aspergillus_aculeatinus | Aspergillus_alliaceus | Aspergillus_aculeatinus |
| | Aspergillus_chevalieri | Aspergillus_alliaceus | Aspergillus_brunneoviolaceus | Aspergillus_aculeatus |
| | Aspergillus_clavatus | Aspergillus_homomorphus | Aspergillus_campestris | Aspergillus_alliaceus |
| | Aspergillus_homomorphus | Aspergillus_melleus | Aspergillus_candidus | Aspergillus_brunneoviolaceus |
| | Aspergillus_japonicus | Aspergillus_nomiae | Aspergillus_chevalieri | Aspergillus_candidus |
| | Aspergillus_luchuensis | Aspergillus_tanneri | Aspergillus_clavatus | Aspergillus_flavus |
| | Aspergillus_nomiae | Aspergillus_tubingensis | Aspergillus_flavus | Aspergillus_homomorphus |
| | Aspergillus_tanneri | Aspergillus_udagawae | Aspergillus_homomorphus | Aspergillus_melleus |
| | Aspergillus_tubingensis | Aspergillus_uvarum | Aspergillus_japonicus | Aspergillus_nomiae |
| | Aspergillus_udagawae | Aspergillus_viridinutans | Aspergillus_luchuensis | Aspergillus_oryzae |
| | Aspergillus_uvarum | Aureobasidium_pullulans | Aspergillus_nomiae | Aspergillus_tanneri |
| | Aspergillus_viridinutans | Babjeviella_inositovora | Aspergillus_oryzae | Aspergillus_tubingensis |
| | Aspergillus_welwitschiae | Bacidia_gigantensis | Aspergillus_tanneri | Aspergillus_udagawae |
| | Aureobasidium_pullulans | Batrachochytrium_dendrobatidis | Aspergillus_tubingensis | Aspergillus_viridinutans |
| | Batrachochytrium_dendrobatidis | Bipolaris_victoriae | Aspergillus_udagawae | Aureobasidium_namibiae |
| | Bipolaris_sorokiniana | Blastomyces_dermatitidis | Aspergillus_viridinutans | Aureobasidium_pullulans |
| | Bipolaris_victoriae | Blastomyces_gilchristii | Aspergillus_welwitschiae | Babjeviella_inositovora |
| | Blastomyces_dermatitidis | Boeremia_exigua | Aureobasidium_namibiae | Bacidia_gigantensis |
| | Boeremia_exigua | Botrytis_cinerea | Aureobasidium_pullulans | Batrachochytrium_dendrobatidis |
| | Botrytis_cinerea | Botrytis_porri | Batrachochytrium_dendrobatidis | Blastomyces_dermatitidis |
| | Botrytis_fragariae | _Candida_auris | Bipolaris_sorokiniana | Blastomyces_gilchristii |
| | Botrytis_porri | Candida_dubliniensis | Blastomyces_dermatitidis | Boeremia_exigua |
| | _Candida_auris | Candida_parapsilosis | Boeremia_exigua | Botrytis_cinerea |
| | _Candida_haemuloni | Candida_tropicalis | Botrytis_cinerea | Brettanomyces_nanus |
| | Candida_parapsilosis | Colletotrichum_higginsianum | Botrytis_fragariae | Candida_albicans |
| | Candida_tropicalis | Cordyceps_militaris | Brettanomyces_nanus | Candida_dubliniensis |
| | Capronia_epimyces | Cryptococcus_amylolentus | Candida_albicans | Capronia_coronata |
| | Ceraceosorus_guamensis | Cryptococcus_neoformans | _Candida_haemuloni | Colletotrichum_fructicola |
| | Chaetomium_globosum | Cutaneotrichosporon_oleaginosum | Capronia_coronata | Coniophora_puteana |
| | Coccidioides_posadasii | Cyberlindnera_jadinii | Capronia_epimyces | Cordyceps_militaris |
| | Colletotrichum_higginsianum | Debaryomyces_fabryi | Ceraceosorus_guamensis | Cryptococcus_amylolentus |
| | Coniosporium_apollinis | Dichomitus_squalens | Chaetomium_globosum | Cryptococcus_neoformans |
| | Cordyceps_militaris | Diutina_rugosa | Coccidioides_posadasii | Cutaneotrichosporon_oleaginosum |
| | Cucurbitaria_berberidis | Dothidotthia_symphoricarpi | Colletotrichum_fructicola | Cyberlindnera_jadinii |
| | Cutaneotrichosporon_oleaginosum | Drechmeria_coniospora | Coniophora_puteana | Dacryopinax_primogenitus |
| | Cyberlindnera_jadinii | Encephalitozoon_cuniculi | Coniosporium_apollinis | Debaryomyces_fabryi |
| | Debaryomyces_fabryi | Encephalitozoon_romaleae | Cordyceps_militaris | Dichomitus_squalens |
| | Diaporthe_citri | Endocarpon_pusillum | Cucurbitaria_berberidis | Dothidotthia_symphoricarpi |
| | Dichomitus_squalens | Exophiala_aquamarina | Cutaneotrichosporon_oleaginosum | Encephalitozoon_romaleae |
| | Diutina_rugosa | Exophiala_oligosperma | Cyberlindnera_jadinii | Endocarpon_pusillum |
| | Drechmeria_coniospora | Exserohilum_turcicum | Dacryopinax_primogenitus | Exophiala_oligosperma |
| | Emericellopsis_atlantica | Fibroporia_radiculosa | Debaryomyces_fabryi | Exophiala_xenobiotica |
| | Encephalitozoon_cuniculi | Filobasidium_floriforme | Diaporthe_citri | Exserohilum_turcicum |
| | Endocarpon_pusillum | Fomitiporia_mediterranea | Dichomitus_squalens | Fibroporia_radiculosa |
| | Eremomyces_bilateralis | Fonsecaea_erecta | Emericellopsis_atlantica | Fonsecaea_erecta |
| | Exophiala_aquamarina | Fonsecaea_pedrosoi | Endocarpon_pusillum | Fonsecaea_nubica |
| | Exophiala_oligosperma | Fusarium_flagelliforme | Eremomyces_bilateralis | Fonsecaea_pedrosoi |
| | Exserohilum_turcicum | Fusarium_fujikuroi | Exophiala_oligosperma | Fusarium_odoratissimum |
| | Filobasidium_floriforme | Fusarium_mangiferae | Exophiala_xenobiotica | Fusarium_oxysporum |
| | Fomitiporia_mediterranea | Fusarium_odoratissimum | Exserohilum_turcicum | Fusarium_proliferatum |
| | Fonsecaea_pedrosoi | Fusarium_oxysporum | Fonsecaea_nubica | Fusarium_pseudograminearum |
| | Fusarium_flagelliforme | Fusarium_proliferatum | Fonsecaea_pedrosoi | Geosmithia_morbida |
| | Fusarium_fujikuroi | Fusarium_pseudograminearum | Fusarium_odoratissimum | Heterobasidion_irregulare |
| | Fusarium_mangiferae | Geosmithia_morbida | Fusarium_proliferatum | Histoplasma_capsulatum |
| | Fusarium_odoratissimum | Grosmannia_clavigera | Fusarium_pseudograminearum | Hyphopichia_burtonii |
| | Fusarium_proliferatum | Heterobasidion_irregulare | Geosmithia_morbida | Komagataella_phaffii |
| | Fusarium_pseudograminearum | Histoplasma_capsulatum | Heterobasidion_irregulare | Kwoniella_bestiolae |
| | Geosmithia_morbida | Hyaloscypha_bicolor | Kluyveromyces_lactis | Kwoniella_mangrovensis |
| | Grosmannia_clavigera | Hyphopichia_burtonii | Kwoniella_bestiolae | Kwoniella_shandongensis |
| | Heterobasidion_irregulare | Komagataella_phaffii | Kwoniella_mangrovensis | Lachancea_thermotolerans |
| | Hyaloscypha_bicolor | Kwoniella_shandongensis | Kwoniella_pini | Lasiodiplodia_theobromae |
| | Kluyveromyces_lactis | Laetiporus_sulphureus | Lachancea_thermotolerans | Leptosphaeria_maculans |

| name | seed_005_059 | seed_006_059 | seed_005_033 | Seed_006_033 |
|---|---|---|---|---|
| trainers (cont.) | Kwoniella_pini | Leptosphaeria_maculans | Lasiodiplodia_theobromae | Lindgomyces_ingoldianus |
| | Laetiporus_sulphureus | Letharia_columbiana | Lentinula_edodes | Malassezia_pachydermatis |
| | Lentinula_edodes | Malassezia_sympodialis | Lindgomyces_ingoldianus | Meira_miltonrushii |
| | Letharia_columbiana | Melampsora_larici-populina | Malassezia_pachydermatis | Metarhizium_album |
| | Malassezia_sympodialis | Metschnikowia_bicuspidata | Meira_miltonrushii | Metschnikowia_bicuspidata |
| | Melampsora_larici-populina | Microsporum_canis | Metarhizium_album | Microsporum_canis |
| | Metarhizium_brunneum | Mixia_osmundae | Metarhizium_brunneum | Mollisia_scopiformis |
| | Microsporum_canis | Mollisia_scopiformis | Microsporum_canis | Morchella_importuna |
| | Mixia_osmundae | Morchella_importuna | Morchella_importuna | Morchella_sextelata |
| | Morchella_importuna | Morchella_sextelata | Morchella_sextelata | Mytilinidion_resinicola |
| | Morchella_sextelata | Mytilinidion_resinicola | Mytilinidion_resinicola | Nematocida_parisii |
| | Mytilinidion_resinicola | Neurospora_tetrasperma | Nematocida_parisii | Neurospora_tetrasperma |
| | Neurospora_tetrasperma | Penicillium_digitatum | Neurospora_tetrasperma | Ogataea_haglerorum |
| | Nosema_ceranae | Pestalotiopsis_fici | Nosema_ceranae | Parastagonospora_nodorum |
| | Ogataea_polymorpha | Pichia_kudriavzevii | Ogataea_haglerorum | Penicillium_digitatum |
| | Pestalotiopsis_fici | Pleurotus_ostreatus | Ogataea_polymorpha | Penicillium_rubens |
| | Pneumocystis_murina | Pneumocystis_jirovecii | Parastagonospora_nodorum | Pestalotiopsis_fici |
| | Pochonia_chlamydosporia | Pneumocystis_murina | Penicillium_rubens | Pichia_kudriavzevii |
| | Podospora_anserina | Pseudogymnoascus_verrucosus | Pestalotiopsis_fici | Pleurotus_ostreatus |
| | Postia_placenta | Pseudomassariella_vexata | Pochonia_chlamydosporia | Pneumocystis_jirovecii |
| | Pseudogymnoascus_destructans | Pseudomicrostroma_glucosiphilum | Podospora_anserina | Protomyces_lactucae-debilis |
| | Pseudogymnoascus_verrucosus | Pseudozyma_hubeiensis | Postia_placenta | Pseudogymnoascus_verrucosus |
| | Pseudomassariella_vexata | Punctularia_strigosozonata | Protomyces_lactucae-debilis | Pseudomassariella_vexata |
| | Pseudozyma_hubeiensis | Pyrenophora_tritici-repentis | Pseudogymnoascus_destructans | Pseudomicrostroma_glucosiphilum |
| | Punctularia_strigosozonata | Rhizophagus_irregularis | Pseudogymnoascus_verrucosus | Pseudozyma_flocculosa |
| | Pyrenophora_tritici-repentis | Saccharomyces_cerevisiae | Pseudomassariella_vexata | Pseudozyma_hubeiensis |
| | Rhinocladiella_mackenziei | Saccharomyces_paradoxus | Pseudozyma_flocculosa | Punctularia_strigosozonata |
| | Rhizophagus_irregularis | Saprochaete_ingens | Pseudozyma_hubeiensis | Pyricularia_oryzae |
| | Saccharomyces_cerevisiae | Scheffersomyces_stipitis | Punctularia_strigosozonata | Pyricularia_pennisetigena |
| | Saccharomyces_paradoxus | Schizosaccharomyces_cryophilus | Pyricularia_oryzae | Rhodotorula_toruloides |
| | Saprochaete_ingens | Schizosaccharomyces_japonicus | Pyricularia_pennisetigena | Saccharomyces_cerevisiae |
| | Scheffersomyces_stipitis | Schizosaccharomyces_pombe | Rhinocladiella_mackenziei | Saccharomycodes_ludwigii |
| | Schizosaccharomyces_cryophilus | Sclerotinia_sclerotiorum | Rhodotorula_toruloides | Saitoella_complicata |
| | Schizosaccharomyces_japonicus | Sordaria_macrospora | Saccharomyces_cerevisiae | Sordaria_macrospora |
| | Schizosaccharomyces_pombe | Sphaerulina_musiva | Saccharomycodes_ludwigii | Spizellomyces_punctatus |
| | Sclerotinia_sclerotiorum | Spizellomyces_punctatus | Saitoella_complicata | Sporisorium_graminicola |
| | Sordaria_macrospora | Sporisorium_graminicola | Sordaria_macrospora | Sporothrix_brasiliensis |
| | Sparassis_crispa | Sugiyamaella_lignohabitans | Sparassis_crispa | Stereum_hirsutum |
| | Spathaspora_passalidarum | Suillus_discolor | Spathaspora_passalidarum | Sugiyamaella_lignohabitans |
| | Sphaerulina_musiva | Suillus_plorans | Spizellomyces_punctatus | Suillus_discolor |
| | Spizellomyces_punctatus | Suillus_subaureus | Sporothrix_brasiliensis | Suillus_plorans |
| | Sugiyamaella_lignohabitans | Synchytrium_microbalum | Stereum_hirsutum | Suillus_subaureus |
| | Suillus_clintonianus | Talaromyces_amestolkiae | Sugiyamaella_lignohabitans | Synchytrium_microbalum |
| | Suillus_paluster | Talaromyces_rugulosus | Suillus_clintonianus | Talaromyces_amestolkiae |
| | Suillus_plorans | Tetrapisispora_blattae | Suillus_paluster | Talaromyces_rugulosus |
| | Talaromyces_rugulosus | Thermothelomyces_thermophilus | Suillus_plorans | Tetrapisispora_blattae |
| | Thermothelomyces_thermophilus | Thermothielavioides_terrestris | Talaromyces_rugulosus | Thermothelomyces_thermophilus |
| | Thermothielavioides_terrestris | Tilletiopsis_washingtonensis | Thermothelomyces_thermophilus | Thermothielavioides_terrestris |
| | Tilletiaria_anomala | Tremella_mesenterica | Thermothielavioides_terrestris | Tilletiopsis_washingtonensis |
| | Tremella_mesenterica | Trichoderma_gamsii | Tilletiaria_anomala | Trametes_versicolor |
| | Trichoderma_reesei | Tuber_melanosporum | Trametes_versicolor | Trichoderma_citrinoviride |
| | Tuber_melanosporum | Uncinocarpus_reesii | Trichoderma_citrinoviride | Trichoderma_gamsii |
| | Uncinocarpus_reesii | Ustilago_hordei | Trichoderma_reesei | Venustampulla_echinocandica |
| | Ustilago_hordei | Venustampulla_echinocandica | Vavraia_culicis | Verticillium_alfalfae |
| | Vavraia_culicis | Wallemia_ichthyophaga | Venustampulla_echinocandica | Wallemia_ichthyophaga |
| | Venustampulla_echinocandica | Wickerhamiella_sorbophila | Verticillium_alfalfae | Wickerhamiella_sorbophila |
| | Wickerhamomyces_anomalus | Wickerhamomyces_anomalus | Wickerhamomyces_anomalus | Wickerhamomyces_anomalus |
| | Xylona_heveae | Xylona_heveae | Yarrowia_lipolytica | Yarrowia_lipolytica |
| | Yarrowia_lipolytica | Yarrowia_lipolytica | Zygosaccharomyces_rouxii | Zygosaccharomyces_rouxii |
| | Zygosaccharomyces_rouxii | Zygosaccharomyces_rouxii | Zymoseptoria_tritici | Zymoseptoria_tritici |

Table S6: Plant species in datasets used for training

| name | Fullmoon_211117_17 | ori9 | seed_002_016 | seed_009_017 | seed_002_017 | seed_009_016 |
|---|---|---|---|---|---|---|
| comment | manual selection | manual selection | automated selection | automated selection | automated selection | automated selection |
| trainers | Creinhardtii<br>Dalata<br>Esalsugineum<br>Graimondii<br>Mguttatus<br>Mpolymorpha<br>MspRCC299<br>Ncolorata<br>Phallii<br>Ppatens<br>Ppersica<br>Ptrifoliata<br>Sitalica<br>Athaliana<br>Bdistachyon<br>Ptrichocarpa<br>Vunguiculata | Creinhardtii<br>Mguttatus<br>Mpolymorpha<br>Sitalica<br>Athaliana<br>Bdistachyon<br>Ptrichocarpa<br>Zmays<br>Gmax | Ahypochondriacus<br>Alyrata<br>Boleraceacapitata<br>Carietinum<br>Cclementina<br>Crubella<br>Csativus<br>Csinensis<br>CsubellipsoideaC169<br>Czofingiensis<br>Dalata<br>Dsalina<br>Esalsugineum<br>Graimondii<br>Lusitatissimum<br>Mesculenta<br>Mguttatus<br>MpusillaCCMP1545<br>Ncolorata<br>Osativa<br>Pacutifolius<br>Phallii<br>Ppatens<br>Ptrichocarpa<br>Ptrifoliata<br>Pvirgatum<br>Slycopersicum<br>Stuberosum<br>Vcarteri<br>Vunguiculata<br>Zmarina<br>Zmays | Acomosus<br>Athaliana<br>Bdistachyon<br>Carietinum<br>Cgrandiflora<br>Creinhardtii<br>Crubella<br>Esalsugineum<br>Fvesca<br>Gmax<br>Ljaponicus<br>Lusitatissimum<br>Macuminata<br>Mesculenta<br>Mguttatus<br>Mpolymorpha<br>MpusillaCCMP1545<br>Mtruncatula<br>Ncolorata<br>Olucimarinus<br>Osativa<br>Othomaeum<br>Pacutifolius<br>Ppersica<br>Pumbilicalis<br>Sbicolor<br>Slycopersicum<br>Sparvula<br>Taestivum<br>Tpratense<br>Vcarteri<br>Zmays | Acomosus<br>Athaliana<br>Boleraceacapitata<br>Carietinum<br>Cgrandiflora<br>Creinhardtii<br>Crubella<br>Csativus<br>Dalata<br>Esalsugineum<br>Fvesca<br>Graimondii<br>Ljaponicus<br>Lusitatissimum<br>Macuminata<br>Mesculenta<br>Mguttatus<br>Mpolymorpha<br>MpusillaCCMP1545<br>Ncolorata<br>Osativa<br>Othomaeum<br>Pacutifolius<br>Ppatens<br>Ptrichocarpa<br>Pvirgatum<br>Slycopersicum<br>Stuberosum<br>Tpratense<br>Vcarteri<br>Zmarina<br>Zmays | Ahypochondriacus<br>Alyrata<br>Bdistachyon<br>Carietinum<br>Cclementina<br>Crubella<br>Csinensis<br>CsubellipsoideaC169<br>Czofingiensis<br>Dsalina<br>Esalsugineum<br>Gmax<br>Lusitatissimum<br>Mesculenta<br>Mguttatus<br>MpusillaCCMP1545<br>Mtruncatula<br>Ncolorata<br>Olucimarinus<br>Osativa<br>Pacutifolius<br>Phallii<br>Ppersica<br>Ptrifoliata<br>Pumbilicalis<br>Sbicolor<br>Slycopersicum<br>Sparvula<br>Taestivum<br>Vcarteri<br>Vunguiculata<br>Zmays |
| skipped validators | Crubella<br>Dsalina<br>Gmax<br>Hannuus<br>Mdomestica<br>Sbicolor<br>Zmays | Crubella<br>Dalata<br>Dsalina<br>Esalsugineum<br>Graimondii<br>Hannuus<br>Mdomestica<br>MspRCC299<br>Ncolorata<br>Phallii<br>Ppatens<br>Ppersica<br>Ptrifoliata<br>Sbicolor<br>Vunguiculata | | | | |

Table S7: Vertebrate species in datasets used for training

| name | seed_020_036 | seed_020_043_ori0_up0_xtr1 | ori6 | seed_004_043 | seed_020_043 |
|---|---|---|---|---|---|
| comment | automated selection | automated selection + tuning | manual selection | automated selection | automated selection |
| trainers | Antrostomus_carolinensis | Alligator_mississippiensis | Anabas_testudineus | Alligator_mississippiensis | Alligator_mississippiensis |
| | Athene_cunicularia | Antrostomus_carolinensis | Drosophila_melanogaster | Calidris_pugnax | Antrostomus_carolinensis |
| | Calidris_pugnax | Calidris_pugnax | Gallus_gallus | Camelus_dromedarius | Calidris_pugnax |
| | Canis_lupus_dingo | Camelus_dromedarius | Mus_musculus | Canis_lupus_dingo | Camelus_dromedarius |
| | Castor_canadensis | Castor_canadensis | Oryzias_latipes | Castor_canadensis | Canis_lupus_dingo |
| | Chelonoidis_abingdonii | Cavia_porcellus | Theropithecus_gelada | Cervus_elaphus | Castor_canadensis |
| | Chinchilla_lanigera | Chelonoidis_abingdonii | | Chlamydotis_macqueenii | Chelonoidis_abingdonii |
| | Chrysochloris_asiatica | Gouania_willdenowi | | Clupea_harengus | Chinchilla_lanigera |
| | Cottoperca_gobio | Lacerta_agilis | | Dasypus_novemcinctus | Gouania_willdenowi |
| | Gouania_willdenowi | Leptosomus_discolor | | Etheostoma_spectabile | Lacerta_agilis |
| | Lates_calcarifer | Macaca_fascicularis | | Lacerta_agilis | Leptonychotes_weddellii |
| | Leptonychotes_weddellii | Mustela_putorius | | Latimeria_chalumnae | Leptosomus_discolor |
| | Megalops_cyprinoides | Myripristis_murdjan | | Leptosomus_discolor | Macaca_fascicularis |
| | Meles_meles | Neogale_vison | | Macaca_fascicularis | Mustela_putorius |
| | Mustela_putorius | Neomonachus_schauinslandi | | Monopterus_albus | Myripristis_murdjan |
| | Neogale_vison | Nothoprocta_perdicaria | | Myripristis_murdjan | Neogale_vison |
| | Nothoprocta_perdicaria | Odobenus_rosmarus | | Neogale_vison | Odobenus_rosmarus |
| | Ochotona_princeps | Oxyura_jamaicensis | | Oncorhynchus_keta | Oxyura_jamaicensis |
| | Odobenus_rosmarus | Panthera_leo | | Oxyura_jamaicensis | Panthera_leo |
| | Panthera_leo | Panthera_tigris | | Panthera_tigris | Panthera_tigris |
| | Pantherophis_guttatus | Pipra_filicauda | | Paramormyrops_kingsleyae | Pantherophis_guttatus |
| | Pan_troglodytes | Protobothrops_mucrosquamatus | | Petromyzon_marinus | Paramormyrops_kingsleyae |
| | Parambassis_ranga | Pygocentrus_nattereri | | Pygocentrus_nattereri | Pipra_filicauda |
| | Piliocolobus_tephrosceles | Sander_lucioperca | | Serinus_canaria | Pygocentrus_nattereri |
| | Pipra_filicauda | Scleropages_formosus | | Sorex_araneus | Sander_lucioperca |
| | Pygoscelis_adeliae | Serinus_canaria | | Struthio_camelus | Serinus_canaria |
| | Rattus_norvegicus | Struthio_camelus | | Tachyglossus_aculeatus | Sorex_araneus |
| | Sander_lucioperca | Tachyglossus_aculeatus | | Talpa_occidentalis | Struthio_camelus |
| | Tachyglossus_aculeatus | Tachysurus_fulvidraco | | Theropithecus_gelada | Tachyglossus_aculeatus |
| | Tachysurus_fulvidraco | Takifugu_rubripes | | Tinamus_guttatus | Tachysurus_fulvidraco |
| | Takifugu_rubripes | Talpa_occidentalis | | Trematomus_bernacchii | Takifugu_rubripes |
| | Thunnus_albacares | | | Varanus_komodoensis | Tinamus_guttatus |
| validation note | | | all non-trainers from vertebrate and invertebrate | | |

Table S8: Invertebrate species in datasets used for training

| name | seed_015_045 | seed_001_045 | seed_022_048_booster_001_039_ori1_up1_xtr0 | seed_001_039 | seed_015_039 |
|---|---|---|---|---|---|
| comment | automated selection | automated selection | automated selection + tuning | automated selection | automated selection |
| trainers | Actinia_tenebrosa<br>Amphimedon_queenslandica<br>Anopheles_coluzzii<br>Anoplophora_glabripennis<br>Caenorhabditis_elegans<br>Drosophila_grimshawi<br>Drosophila_miranda<br>Drosophila_teissieri<br>Haliotis_rubra<br>Homalodisca_vitripennis<br>Ischnura_elegans<br>Loa_loa<br>Nasonia_vitripennis<br>Necator_americanus<br>Neodiprion_fabricii<br>Neodiprion_virginiana<br>Nicrophorus_vespilloides<br>Nomia_melanderi<br>Orussus_abietinus<br>Pogonomyrmex_barbatus<br>Pomacea_canaliculata<br>Portunus_trituberculatus<br>Rhagoletis_zephyria<br>Salpingoeca_rosetta<br>Schistosoma_haematobium<br>Sphaeroforma_arctica<br>Spodoptera_frugiperda<br>Strongylocentrotus_purpuratus<br>Tetranychus_urticae<br>Varroa_jacobsoni<br>Xenia_sp._Carnegie-2017<br>Zerene_cesonia | Acyrthosiphon_pisum<br>Anopheles_albimanus<br>Anopheles_coluzzii<br>Anoplophora_glabripennis<br>Aphis_gossypii<br>Bemisia_tabaci<br>Bicyclus_anynana<br>Chelonus_insularis<br>Drosophila_elegans<br>Drosophila_grimshawi<br>Drosophila_rhopaloa<br>Drosophila_simulans<br>Drosophila_teissieri<br>Fonticula_alba<br>Haliotis_rubra<br>Homalodisca_vitripennis<br>Ischnura_elegans<br>Loa_loa<br>Lottia_gigantea<br>Lucilia_sericata<br>Nasonia_vitripennis<br>Neodiprion_virginiana<br>Nicrophorus_vespilloides<br>Pogonomyrmex_barbatus<br>Pseudomyrmex_gracilis<br>Rhagoletis_zephyria<br>Salpingoeca_rosetta<br>Schistosoma_haematobium<br>Spodoptera_frugiperda<br>Strongylocentrotus_purpuratus<br>Zerene_cesonia<br>Zootermopsis_nevadensis | Acromyrmex_echinatior<br>Actinia_tenebrosa<br>Acyrthosiphon_pisum<br>Anopheles_albimanus<br>Anopheles_coluzzii<br>Anopheles_stephensi<br>Aphis_gossypii<br>Apis_mellifera<br>Bemisia_tabaci<br>Bicyclus_anynana<br>Bombyx_mori<br>Bradysia_coprophila<br>Brugia_malayi<br>Caenorhabditis_elegans<br>Ceratitis_capitata<br>Chelonus_insularis<br>Ciona_intestinalis<br>Cryptotermes_secundus<br>Daphnia_pulicaria<br>Dendroctonus_ponderosae<br>Dermacentor_silvarum<br>Diachasma_alloeum<br>Diuraphis_noxia<br>Drosophila_biarmipes<br>Drosophila_busckii<br>Drosophila_elegans<br>Drosophila_kikkawai<br>Drosophila_melanogaster<br>Drosophila_navojoa<br>Drosophila_rhopaloa<br>Drosophila_santomea<br>Drosophila_serrata<br>Drosophila_simulans<br>Drosophila_teissieri<br>Exaiptasia_diaphana<br>Fonticula_alba<br>Gigantopelta_aegis<br>Haliotis_rubra<br>Hermetia_illucens<br>Ixodes_scapularis<br>Leptopilina_heterotoma<br>Lottia_gigantea<br>Lucilia_sericata<br>Megalopta_genalis<br>Melanaphis_sacchari<br>Neodiprion_pinetum<br>Nylanderia_fulva<br>Octopus_bimaculoides<br>Odontomachus_brunneus<br>Ooceraea_biroi<br>Papilio_machaon<br>Parasteatoda_tepidariorum<br>Penaeus_japonicus<br>Pieris_rapae<br>Plutella_xylostella<br>Pogonomyrmex_barbatus<br>Pomacea_canaliculata<br>Pseudomyrmex_gracilis<br>Salpingoeca_rosetta<br>Thrips_palmi<br>Trichoplax_adhaerens<br>Varroa_destructor<br>Vespa_crabro<br>Vespula_pensylvanica<br>Zootermopsis_nevadensis | Acromyrmex_echinatior<br>Acyrthosiphon_pisum<br>Anopheles_albimanus<br>Aphis_gossypii<br>Bemisia_tabaci<br>Bicyclus_anynana<br>Bombyx_mori<br>Chelonus_insularis<br>Cryptotermes_secundus<br>Diachasma_alloeum<br>Drosophila_biarmipes<br>Drosophila_elegans<br>Drosophila_kikkawai<br>Drosophila_rhopaloa<br>Drosophila_santomea<br>Drosophila_simulans<br>Exaiptasia_diaphana<br>Fonticula_alba<br>Gigantopelta_aegis<br>Haliotis_rubra<br>Lottia_gigantea<br>Lucilia_sericata<br>Megalopta_genalis<br>Neodiprion_pinetum<br>Odontomachus_brunneus<br>Penaeus_japonicus<br>Plutella_xylostella<br>Pogonomyrmex_barbatus<br>Pseudomyrmex_gracilis<br>Salpingoeca_rosetta<br>Trichoplax_adhaerens<br>Zootermopsis_nevadensis | Acromyrmex_echinatior<br>Actinia_tenebrosa<br>Amphimedon_queenslandica<br>Bombyx_mori<br>Caenorhabditis_elegans<br>Cryptotermes_secundus<br>Diachasma_alloeum<br>Drosophila_biarmipes<br>Drosophila_kikkawai<br>Drosophila_miranda<br>Drosophila_santomea<br>Exaiptasia_diaphana<br>Gigantopelta_aegis<br>Haliotis_rubra<br>Megalopta_genalis<br>Necator_americanus<br>Neodiprion_fabricii<br>Neodiprion_pinetum<br>Nomia_melanderi<br>Odontomachus_brunneus<br>Orussus_abietinus<br>Penaeus_japonicus<br>Plutella_xylostella<br>Pogonomyrmex_barbatus<br>Pomacea_canaliculata<br>Portunus_trituberculatus<br>Salpingoeca_rosetta<br>Sphaeroforma_arctica<br>Tetranychus_urticae<br>Trichoplax_adhaerens<br>Varroa_jacobsoni<br>Xenia_sp._Carnegie-2017 |

#### S2.3 Training Parameters

Table S9: Fungi Released Model Parameters

| parameter | fungi_v0.3_a_0100 | fungi_v0.3_a_0200 | fungi_v0.3_a_0300 | fungi_v0.3_a_0400 |
|---|---|---|---|---|
| data_dir | seed_005_059 | seed_006_059 | seed_005_033 | seed_006_033 |
| data_helixer_commit | ac7ba57 | ac7ba57 | ac7ba57 | ac7ba57 |
| data_geenuff_commit | 46418ef | 46418ef | 46418ef | 46418ef |
| training_helixer_commit | ac7ba57 | ac7ba57 | ac7ba57 | ac7ba57 |
| pool_size | 9 | 9 | 9 | 9 |
| batch_size | 140 | 140 | 140 | 140 |
| val_test_batch_size | 280 | 280 | 280 | 280 |
| class_weights | [0.7,1.6,1.2,1.2] | [0.7,1.6,1.2,1.2] | [0.7,1.6,1.2,1.2] | [0.7,1.6,1.2,1.2] |
| transition_weights | [1,12,3,1,12,3] | [1,12,3,1,12,3] | [1,12,3,1,12,3] | [1,12,3,1,12,3] |
| save_every_check | True | True | True | True |
| predict_phase | True | True | True | True |
| lstm_layers | 3 | 3 | 3 | 3 |
| cnn_layers | 4 | 4 | 4 | 4 |
| units | 128 | 128 | 128 | 128 |
| filter_depth | 96 | 96 | 96 | 96 |
| kernel_size | 9 | 9 | 9 | 9 |
| check_every_nth_batch | 1000 | 1000 | 1000 | 1000 |
| patience | 6 | 6 | 6 | 6 |

Table S10: Plant Released Model Parameters

| parameter | land_plant_v0.3_a_0080 | land_plant_v0.3_a_0090 | land_plant_v0.3_a_0100 | land_plant_v0.3_a_0200 |
|---|---|---|---|---|
| data_dir | seed_009_017 | seed_002_017 | seed_009_017 | seed_009_016 |
| data_helixer_commit | ac7ba57 | ac7ba57 | 02e5e94 | 02e5e94 |
| data_geenuff_commit | 46418ef | 46418ef | 46418ef | 46418ef |
| training_helixer_commit | ac7ba57 | ac7ba57 | 02e5e94 | 02e5e94 |
| kernel_size | 12 | 10 | 10 | 10 |
| learning_rate | 0.0003 | 0.0003 | NaN | NaN |
| pool_size | 9 | 9 | 9 | 9 |
| batch_size | 140 | 140 | 50 | 50 |
| val_test_batch_size | 280 | 280 | 100 | 100 |
| class_weights | [0.7,1.6,1.2,1.2] | [0.7,1.6,1.2,1.2] | [0.7,1.6,1.2,1.2] | [0.7,1.6,1.2,1.2] |
| transition_weights | [1,12,3,1,12,3] | [1,12,3,1,12,3] | [1,12,3,1,12,3] | [1,12,3,1,12,3] |
| save_every_check | True | True | True | True |
| predict_phase | True | True | True | True |
| lstm_layers | 3 | 3 | 3 | 3 |
| cnn_layers | 4 | 4 | 4 | 4 |
| units | 128 | 128 | 128 | 128 |
| filter_depth | 96 | 96 | 96 | 96 |
| check_every_nth_batch | 1000 | 1000 | 5000 | 5000 |
| patience | 9 | 9 | 7 | 7 |
| save_every_epoch | NaN | NaN | NaN | NaN |

| parameter | land_plant_v0.3_a_0300 | land_plant_v0.3_a_0400 | land_plant_v0.3_m_0100 | land_plant_v0.3_m_0200 |
|---|---|---|---|---|
| data_dir | seed_002_016 | seed_002_017 | fullmoon_211117_17 | ori9 |
| data_helixer_commit | 02e5e94 | 02e5e94 | bb840b4 | bb840b4 |
| data_geenuff_commit | 46418ef | 46418ef | 1f6cffb | 1f6cffb |
| training_helixer_commit | 02e5e94 | 02e5e94 | bb840b4 | bb840b4 |
| kernel_size | 10 | 10 | 10 | 10 |
| learning_rate | NaN | NaN | NaN | NaN |
| pool_size | 9 | 9 | 9 | 9 |
| batch_size | 50 | 50 | 50 | 50 |
| val_test_batch_size | 100 | 100 | 100 | 100 |
| class_weights | [0.7,1.6,1.2,1.2] | [0.7,1.6,1.2,1.2] | [0.7,1.6,1.2,1.2] | [0.7,1.6,1.2,1.2] |
| transition_weights | [1,12,3,1,12,3] | [1,12,3,1,12,3] | [1,12,3,1,12,3] | [1,12,3,1,12,3] |
| save_every_check | True | True | NaN | NaN |
| predict_phase | True | True | True | True |
| lstm_layers | 3 | 3 | 3 | 3 |
| cnn_layers | 4 | 4 | 4 | 4 |
| units | 128 | 128 | 128 | 128 |
| filter_depth | 96 | 96 | 96 | 96 |
| check_every_nth_batch | 5000 | 5000 | NaN | NaN |
| patience | 7 | 7 | NaN | NaN |
| save_every_epoch | NaN | NaN | True | True |

Table S11: Vertebrate Released Model Parameters

| parameter | vertebrate_v0.3_a_0200 | vertebrate_v0.3_a_0300 | vertebrate_v0.3_a_0400 |
|---|---|---|---|
| data_dir | seed_020_043 | seed_020_036 | seed_004_043 |
| data_helixer_commit | ac7ba57 | ac7ba57 | ac7ba57 |
| data_geenuff_commit | 46418ef | 46418ef | 46418ef |
| training_helixer_commit | ac7ba57 | ac7ba57 | ac7ba57 |
| pool_size | 9 | 9 | 9 |
| batch_size | 140 | 140 | 140 |
| val_test_batch_size | 280 | 280 | 280 |
| class_weights | [0.7,1.6,1.2,1.2] | [0.7,1.6,1.2,1.2] | [0.7,1.6,1.2,1.2] |
| transition_weights | [1,12,3,1,12,3] | [1,12,3,1,12,3] | [1,12,3,1,12,3] |
| save_every_check | True | True | True |
| predict_phase | True | True | True |
| lstm_layers | 3 | 3 | 3 |
| cnn_layers | 4 | 4 | 4 |
| units | 128 | 128 | 128 |
| filter_depth | 96 | 96 | 96 |
| kernel_size | 9 | 9 | 9 |
| check_every_nth_batch | 1000 | 1000 | 1000 |
| patience | 6 | 6 | 6 |
| manual_checkpoint_selection | NaN | NaN | NaN |

| parameter | vertebrate_v0.3_m0100 | vertebrate_v0.3_m_0080 | vertebrate_v0.3_m_0090 |
|---|---|---|---|
| data_dir | ori6 | seed_020_043_ori0_up0_xtr1 | seed_020_043_ori0_up0_xtr1 |
| data_helixer_commit | 02e5e94 | ac7ba57 | ac7ba57 |
| data_geenuff_commit | 46418ef | 46418ef | 46418ef |
| training_helixer_commit | 02e5e94 | 82bf1ca | 82bf1ca |
| pool_size | 9 | 9 | 9 |
| batch_size | 240 | 120 | 120 |
| val_test_batch_size | 480 | 240 | 240 |
| class_weights | [0.7,1.6,1.2,1.2] | [0.7,1.6,1.2,1.2] | [0.7,1.6,1.2,1.2] |
| transition_weights | [1,12,3,1,12,3] | [1,12,3,1,12,3] | [1,12,3,1,12,3] |
| save_every_check | True | True | True |
| predict_phase | True | True | True |
| lstm_layers | 3 | 4 | 4 |
| cnn_layers | 4 | 4 | 4 |
| units | 128 | 128 | 128 |
| filter_depth | 96 | 128 | 128 |
| kernel_size | 10 | 10 | 10 |
| check_every_nth_batch | 1000 | 1300 | 1300 |
| patience | 7 | 9 | 9 |
| manual_checkpoint_selection | NaN | model_e2_b003899.h5 -> model_e0_b002599.h5 | model_e2_b003899.h5 -> model_e0_b003899.h5 |

Table S12: Invertebrate Released Model Parameters

| parameter | invertebrate_v0.3_a_0300 | invertebrate_v0.3_a_0400 | invertebrate_v0.3_a_0500 |
|---|---|---|---|
| data_dir | seed_001_045 | seed_001_039 | seed_015_045 |
| data_helixer_commit | ac7ba57 | ac7ba57 | ac7ba57 |
| data_geenuff_commit | 46418ef | 46418ef | 46418ef |
| training_helixer_commit | ac7ba57 | ac7ba57 | ac7ba57 |
| pool_size | 9 | 9 | 9 |
| batch_size | 140 | 140 | 140 |
| val_test_batch_size | 280 | 280 | 280 |
| class_weights | [0.7,1.6,1.2,1.2] | [0.7,1.6,1.2,1.2] | [0.7,1.6,1.2,1.2] |
| transition_weights | [1,12,3,1,12,3] | [1,12,3,1,12,3] | [1,12,3,1,12,3] |
| save_every_check | True | True | True |
| predict_phase | True | True | True |
| lstm_layers | 3 | 3 | 3 |
| cnn_layers | 4 | 4 | 4 |
| units | 128 | 128 | 128 |
| filter_depth | 96 | 96 | 96 |
| kernel_size | 9 | 9 | 9 |
| check_every_nth_batch | 1000 | 1000 | 1000 |
| patience | 6 | 6 | 6 |
| manual_checkpoint_selection | NaN | NaN | NaN |

| parameter | invertebrate_v0.3_a_0600 | invertebrate_v0.3_m_0100 | invertebrate_v0.3_m_0200 |
|---|---|---|---|
| data_dir | seed_015_039 | seed_022_048_booster_001_039_ori1_up1_xtr0 | seed_022_048_booster_001_039_ori1_up1_xtr0 |
| data_helixer_commit | ac7ba57 | ac7ba57 | ac7ba57 |
| data_geenuff_commit | 46418ef | 46418ef | 46418ef |
| training_helixer_commit | ac7ba57 | 82bf1ca | 82bf1ca |
| pool_size | 9 | 9 | 9 |
| batch_size | 140 | 120 | 120 |
| val_test_batch_size | 280 | 240 | 240 |
| class_weights | [0.7,1.6,1.2,1.2] | [0.7,1.6,1.2,1.2] | [0.7,1.6,1.2,1.2] |
| transition_weights | [1,12,3,1,12,3] | [1,12,3,1,12,3] | [1,12,3,1,12,3] |
| save_every_check | True | True | True |
| predict_phase | True | True | True |
| lstm_layers | 3 | 4 | 4 |
| cnn_layers | 4 | 4 | 4 |
| units | 128 | 128 | 128 |
| filter_depth | 96 | 128 | 128 |
| kernel_size | 9 | 10 | 10 |
| check_every_nth_batch | 1000 | 1300 | 1300 |
| patience | 6 | 9 | 9 |
| manual_checkpoint_selection | NaN | model_e3_b006499.h5 | model_e3_b014299.h5 |

#### S2.4 Additional Information

Table S13: Non default and result affecting parameters used for various bioinformatics tools unless otherwise specified.

| tool | parameter | value |
|---|---|---|
| HelixerPost | | |
| helixer_post_bin | windowSize | 100 |
| helixer_post_bin | edgeThresh | 0.1 |
| helixer_post_bin | peakThresh | 0.8 |
| helixer_post_bin | minCodingLength | 60 |
| AUGUSTUS | | |
| augustus | softmasking | 1 |
| augustus | noInFrameStop | "true" |
| augustus | stopCodonExcludedFromCDS | "false" |
| augustus | gff3 | on |
| augustus | UTR | on (if available) |
| GenemarkES | | |
| gmes_petap.pl | ES | |
| gmes_petap.pl | fungus (for fungi only) | |
| BUSCO | | |
| busco | mode | prot |
| busco | lineage | metazoa_odb10 (vertebrates and invertebrates) |
| busco | lineage | viridiplantae_odb10 (land_plants) |
| busco | lineage | fungi_odb10 (fungi) |
| Mercator4 | | |
| mercator4 | sequence type | protein |
| mercator4 | include prot-scriber | TRUE |
| mercator4 | include swissprot | TRUE |

Table S14: Configuration and metadata used for the automated training species selection process.

| Unnamed: 0 | fungi | plants | vertebrates | invertebrates |
|---|---|---|---|---|
| total species | 298 | 77 | 314 | 201 |
| seeds per split | 32 | 12 | 24 | 24 |
| training species (per split) | 64 | 16 | 16 | 16 |

Table S15: AUGUSTUS models (where available) used for test species

| Species | AUGUSTUS closest | Helixer lineage |
|---|---|---|
| Rhincodon_typus | rhincodon | vertebrate |
| Schistosoma_mansoni | (schistosoma2) | invertebrate |
| Apis_dorsata | adorsata | invertebrate |
| Bombus_impatiens | bombus_impatiens1 | invertebrate |
| Tribolium_castaneum | tribolium2012 | invertebrate |
| Culex_pipiens | culex | invertebrate |
| Neurospora_crassa | (neurospora) | fungi |
| Laccaria_bicolor | laccaria_bicolor | fungi |
| Fusarium_graminearum | (fusarium) | fungi |
| Debaryomyces_hansenii | debaryomyces_hansenii | fungi |
| Coccidioides_immitis | coccidioides_immitis | fungi |
| Aspergillus_terreus | aspergillus_terreus | fungi |
| Nicotiana_attenuata | coyote_tobacco | land_plant |
| Triticum_dicoccoides | wheat | land_plant |
| Solanum_pennellii | tomato | land_plant |
| Oryza_brachyantha | rice | land_plant |

Table S16: Additional length parameters used for Helixer inference on test species

| Unnamed: 0 | subsequence length | overlap offset | overlap core length |
|---|---|---|---|
| fungi | 21384 | 10692 | 16038 |
| land_plant | 106920 | 53460 | 80190 |
| invertebrate | 213840 | 106920 | 160380 |
| vertebrate | 213840 | 106920 | 160380 |

Table S17: Parameters changed in training ablation models as compared to best land_plant model (also shown, v0.3_a_0080)

| name | architecture | predict phase | transition weights |
|---|---|---|---|
| land_plant_v0.3_a_0080 | HybridModel | True | [1, 12, 3, 1, 12, 3] |
| transition_weights_medium | HybridModel | True | [1, 8, 2, 1, 8, 2] |
| transition_weights_low | HybridModel | True | [1, 4, 1.5, 1, 4, 1.5] |
| transition_weights_none | HybridModel | True | [1, 1, 1, 1, 1, 1] |
| phase_none | HybridModel | False | [1, 12, 3, 1, 12, 3] |
| LSTM | LSTMModel | True | [1, 12, 3, 1, 12, 3] |

Table S18: The overlapping parameters during inference in the ablation analysis differ slightly from other analyses, but were constant for all models and species.

| parameter | value |
|---|---|
| subsequence length | 106920 |
| overlap | TRUE |
| overlap offset | 13365 |
| core length | 53460 |

Table S19: Performance affecting parameters set during benchmarking. Machines described in methods. Note that setting compression lzf, and the temporary dir to a path on an SSD, improve performance where the installation and hardware, respectively, allow. The highest batch size that fits in GPU memory should be fastest. No multiprocess is set for interpretability of the resulting numbers and a more fair comparison between tools.

| machine | compression | batch size | temporary dir | no multiprocess |
|---|---|---|---|---|
| A | lzf | 64 | <on SSD> | True |
| B | lzf | 64 | <on SSD> | True |
| C | lzf | 58 | <on SSD> | True |

#### S2.5 Tabular Results

Table S20: Subgenic F1 for test species.

| species | helixer | helixer_post | genemark | augustus | group |
|---|---|---|---|---|---|
| Neurospora_crassa | 0.9462 | 0.9517 | 0.9422 | 0.8854 | fungi |
| Coccidioides_immitis | 0.9293 | 0.9313 | 0.9230 | 0.8879 | fungi |
| Fusarium_graminearum | 0.9540 | 0.9536 | 0.9446 | 0.9181 | fungi |
| Debaryomyces_hansenii | 0.9937 | 0.9937 | 0.9880 | 0.9630 | fungi |
| Aspergillus_terreus | 0.9617 | 0.9609 | 0.9379 | 0.9296 | fungi |
| Laccaria_bicolor | 0.8562 | 0.8543 | 0.8268 | 0.8020 | fungi |
| Papaver_somniferum | 0.5209 | 0.5589 | 0.3230 | NaN | plant |
| Arachis_hypogaea | 0.7914 | 0.8031 | 0.2831 | NaN | plant |
| Brassica_napus | 0.8051 | 0.8158 | 0.5507 | NaN | plant |
| Oryza_brachyantha | 0.9038 | 0.9107 | 0.2792 | 0.7133 | plant |
| Nicotiana_attenuata | 0.7789 | 0.8046 | 0.2081 | 0.5104 | plant |
| Solanum_pennellii | 0.8648 | 0.8754 | 0.4171 | 0.5402 | plant |
| Setaria_viridis | 0.8544 | 0.8642 | 0.2477 | NaN | plant |
| Vitis_riparia | 0.8989 | 0.9005 | 0.5207 | NaN | plant |
| Triticum_dicoccoides | 0.5165 | 0.5898 | NaN | 0.3869 | plant |
| Phoenix_dactylifera | 0.8836 | 0.8873 | 0.2258 | NaN | plant |
| Coffea_arabica | 0.8129 | 0.8203 | 0.4924 | NaN | plant |
| Cannabis_sativa | 0.8207 | 0.8248 | 0.3653 | NaN | plant |
| Hibiscus_syriacus | 0.6262 | 0.6387 | 0.3675 | NaN | plant |
| Balaenoptera_musculus | 0.8613 | 0.9032 | 0.1657 | NaN | vertebrate |
| Xiphias_gladius | 0.9132 | 0.9244 | 0.6495 | NaN | vertebrate |
| Sparus_aurata | 0.8797 | 0.8881 | 0.6009 | NaN | vertebrate |
| Desmodus_rotundus | 0.8800 | 0.8996 | 0.1927 | NaN | vertebrate |
| Pseudonaja_textilis | 0.8886 | 0.8949 | 0.1870 | NaN | vertebrate |
| Falco_naumanni | 0.8542 | 0.8916 | 0.1925 | NaN | vertebrate |
| Chiroxiphia_lanceolata | 0.8996 | 0.9189 | 0.0753 | NaN | vertebrate |
| Myotis_davidii | 0.8548 | 0.8697 | 0.1913 | NaN | vertebrate |
| Lagopus_leucura | 0.8730 | 0.9021 | 0.1498 | NaN | vertebrate |
| Rhincodon_typus | 0.8105 | 0.8347 | 0.0690 | 0.5661 | vertebrate |
| Opisthocomus_hoazin | 0.8625 | 0.8757 | 0.2853 | NaN | vertebrate |
| Bombus_impatiens | 0.8615 | 0.8628 | 0.4493 | 0.5383 | invertebrate |
| Tribolium_castaneum | 0.8640 | 0.8613 | 0.6107 | 0.6614 | invertebrate |
| Drosophila_virilis | 0.8640 | 0.8648 | 0.5307 | NaN | invertebrate |
| Apis_dorsata | 0.8454 | 0.8521 | 0.4845 | 0.4583 | invertebrate |
| Frankliniella_occidentalis | 0.7962 | 0.8097 | 0.4084 | NaN | invertebrate |
| Trachymyrmex_cornetzi | 0.8188 | 0.8223 | 0.3689 | NaN | invertebrate |
| Drosophila_albomicans | 0.8774 | 0.8748 | 0.5636 | NaN | invertebrate |
| Ctenocephalides_felis | 0.7196 | 0.7393 | 0.4535 | NaN | invertebrate |
| Atta_cephalotes | 0.8717 | 0.8710 | 0.5334 | NaN | invertebrate |
| Rhopalosiphum_maidis | 0.9008 | 0.8953 | 0.5331 | NaN | invertebrate |
| Culex_pipiens | 0.8046 | 0.8120 | 0.4476 | 0.5602 | invertebrate |
| Hyposmocoma_kahamanoa | 0.8100 | 0.8075 | 0.4953 | NaN | invertebrate |
| Caenorhabditis_remanei | 0.9098 | 0.9055 | 0.9231 | NaN | invertebrate |
| Schistosoma_mansoni | 0.8057 | 0.7956 | 0.8361 | 0.8421 | invertebrate |
| Opisthorchis_viverrini | 0.5086 | 0.4836 | 0.6328 | NaN | invertebrate |

Table S21: Fraction of complete BUSCOs for test species.

| species | reference | helixer_post | genemark | augustus | group |
|---|---|---|---|---|---|
| Neurospora_crassa | 1.000000 | 0.998681 | 1.000000 | 0.984169 | fungi |
| Coccidioides_immitis | 0.996042 | 0.997361 | 0.990765 | 0.967018 | fungi |
| Fusarium_graminearum | 0.982850 | 0.994723 | 0.990765 | 0.990765 | fungi |
| Debaryomyces_hansenii | 0.978892 | 0.982850 | 0.973615 | 0.961741 | fungi |
| Aspergillus_terreus | 0.932718 | 0.996042 | 0.990765 | 0.980211 | fungi |
| Laccaria_bicolor | 0.905013 | 0.972296 | 0.963061 | 0.912929 | fungi |
| Papaver_somniferum | 1.000000 | 0.997647 | 0.851765 | NaN | plant |
| Arachis_hypogaea | 1.000000 | 0.992941 | 0.738824 | NaN | plant |
| Brassica_napus | 0.997647 | 0.997647 | 0.995294 | NaN | plant |
| Oryza_brachyantha | 0.997647 | 0.997647 | 0.098824 | 0.720000 | plant |
| Nicotiana_attenuata | 0.997647 | 0.971765 | 0.296471 | 0.844706 | plant |
| Solanum_pennellii | 0.995294 | 0.978824 | 0.743529 | 0.868235 | plant |
| Setaria_viridis | 0.995294 | 0.995294 | 0.131765 | NaN | plant |
| Vitis_riparia | 0.995294 | 0.952941 | 0.687059 | NaN | plant |
| Triticum_dicoccoides | 0.990588 | 0.992941 | NaN | 0.736471 | plant |
| Phoenix_dactylifera | 0.985882 | 0.945882 | 0.103529 | NaN | plant |
| Coffea_arabica | 0.983529 | 0.960000 | 0.771765 | NaN | plant |
| Cannabis_sativa | 0.971765 | 0.957647 | 0.776471 | NaN | plant |
| Hibiscus_syriacus | 0.964706 | 0.962353 | 0.931765 | NaN | plant |
| Balaenoptera_musculus | 0.997904 | 0.961216 | 0.105870 | NaN | vertebrate |
| Xiphias_gladius | 0.991614 | 0.977987 | 0.905660 | NaN | vertebrate |
| Sparus_aurata | 0.991614 | 0.959119 | 0.920335 | NaN | vertebrate |
| Desmodus_rotundus | 0.988470 | 0.959119 | 0.164570 | NaN | vertebrate |
| Pseudonaja_textilis | 0.980084 | 0.930818 | 0.078616 | NaN | vertebrate |
| Falco_naumanni | 0.952830 | 0.916143 | 0.066038 | NaN | vertebrate |
| Chiroxiphia_lanceolata | 0.946541 | 0.920335 | 0.000000 | NaN | vertebrate |
| Myotis_davidii | 0.943396 | 0.903564 | 0.162474 | NaN | vertebrate |
| Lagopus_leucura | 0.937107 | 0.903564 | 0.085954 | NaN | vertebrate |
| Rhincodon_typus | 0.857442 | 0.784067 | 0.011530 | 0.578616 | vertebrate |
| Opisthocomus_hoazin | 0.779874 | 0.734801 | 0.035639 | NaN | vertebrate |
| Bombus_impatiens | 0.996855 | 0.986373 | 0.950734 | 0.706499 | invertebrate |
| Tribolium_castaneum | 0.994759 | 0.903564 | 0.938155 | 0.765199 | invertebrate |
| Drosophila_virilis | 0.993711 | 0.977987 | 0.932914 | NaN | invertebrate |
| Apis_dorsata | 0.992662 | 0.985325 | 0.970650 | 0.723270 | invertebrate |
| Frankliniella_occidentalis | 0.989518 | 0.973795 | 0.029350 | NaN | invertebrate |
| Trachymyrmex_cornetzi | 0.987421 | 0.961216 | 0.658281 | NaN | invertebrate |
| Drosophila_albomicans | 0.973795 | 0.961216 | 0.930818 | NaN | invertebrate |
| Ctenocephalides_felis | 0.969602 | 0.942348 | 0.909853 | NaN | invertebrate |
| Atta_cephalotes | 0.968553 | 0.939203 | 0.661426 | NaN | invertebrate |
| Rhopalosiphum_maidis | 0.967505 | 0.943396 | 0.828092 | NaN | invertebrate |
| Culex_pipiens | 0.963312 | 0.942348 | 0.921384 | 0.910901 | invertebrate |
| Hyposmocoma_kahamanoa | 0.923480 | 0.829140 | 0.831237 | NaN | invertebrate |
| Caenorhabditis_remanei | 0.754717 | 0.711740 | 0.729560 | NaN | invertebrate |
| Schistosoma_mansoni | 0.709644 | 0.396226 | 0.533543 | 0.459119 | invertebrate |
| Opisthorchis_viverrini | 0.647799 | 0.179245 | 0.573375 | NaN | invertebrate |

Table S22: Mapman4 annotation counts for test plant species

| %Annotated | %Classified | %Bins occupied | harmonic | annotation | species |
|---|---|---|---|---|---|
| 70.91 | 11.06 | 88.67 | 78.80 | genemark | Arachis_hypogaea |
| 85.23 | 27.86 | 96.63 | 90.57 | genemark | Brassica_napus |
| 77.29 | 13.67 | 86.79 | 81.76 | genemark | Cannabis_sativa |
| 76.39 | 20.08 | 87.64 | 81.63 | genemark | Coffea_arabica |
| 87.02 | 25.66 | 95.54 | 91.08 | genemark | Hibiscus_syriacus |
| 77.68 | 6.10 | 67.66 | 72.32 | genemark | Nicotiana_attenuata |
| 54.23 | 18.89 | 31.45 | 39.81 | genemark | Oryza_brachyantha |
| 75.20 | 12.13 | 90.30 | 82.06 | genemark | Papaver_somniferum |
| 39.62 | 9.74 | 32.30 | 35.59 | genemark | Phoenix_dactylifera |
| 49.81 | 14.22 | 36.69 | 42.26 | genemark | Setaria_viridis |
| 86.60 | 15.71 | 87.72 | 87.16 | genemark | Solanum_pennellii |
| NaN | NaN | NaN | NaN | genemark | Triticum_dicoccoides |
| 82.93 | 24.62 | 88.79 | 85.76 | genemark | Vitis_riparia |
| 92.82 | 48.92 | 97.11 | 94.92 | helixer | Arachis_hypogaea |
| 92.70 | 47.74 | 97.75 | 95.16 | helixer | Brassica_napus |
| 94.73 | 54.03 | 93.58 | 94.15 | helixer | Cannabis_sativa |
| 89.91 | 46.45 | 95.75 | 92.74 | helixer | Coffea_arabica |
| 93.49 | 46.66 | 96.75 | 95.09 | helixer | Hibiscus_syriacus |
| 86.37 | 38.39 | 96.25 | 91.04 | helixer | Nicotiana_attenuata |
| 89.48 | 52.68 | 94.66 | 92.00 | helixer | Oryza_brachyantha |
| 68.48 | 17.53 | 96.54 | 80.12 | helixer | Papaver_somniferum |
| 90.41 | 52.86 | 95.18 | 92.73 | helixer | Phoenix_dactylifera |
| 86.45 | 46.06 | 95.02 | 90.53 | helixer | Setaria_viridis |
| 90.69 | 48.44 | 95.83 | 93.19 | helixer | Solanum_pennellii |
| 71.53 | 18.53 | 94.88 | 81.57 | helixer | Triticum_dicoccoides |
| 94.60 | 53.13 | 96.23 | 95.41 | helixer | Vitis_riparia |
| 96.50 | 50.73 | 94.67 | 95.58 | reference | Arachis_hypogaea |
| 97.62 | 53.49 | 97.94 | 97.78 | reference | Brassica_napus |
| 98.35 | 50.31 | 91.65 | 94.88 | reference | Cannabis_sativa |
| 97.27 | 52.55 | 92.24 | 94.69 | reference | Coffea_arabica |
| 98.96 | 59.32 | 95.37 | 97.13 | reference | Hibiscus_syriacus |
| 93.48 | 46.75 | 94.05 | 93.76 | reference | Nicotiana_attenuata |
| 96.43 | 60.15 | 91.44 | 93.87 | reference | Oryza_brachyantha |
| 93.11 | 44.73 | 94.40 | 93.75 | reference | Papaver_somniferum |
| 97.93 | 60.07 | 88.50 | 92.98 | reference | Phoenix_dactylifera |
| 95.70 | 55.63 | 90.92 | 93.25 | reference | Setaria_viridis |
| 97.68 | 57.33 | 93.33 | 95.46 | reference | Solanum_pennellii |
| 93.94 | 52.39 | 93.55 | 93.74 | reference | Triticum_dicoccoides |
| 98.59 | 57.53 | 92.69 | 95.55 | reference | Vitis_riparia |

#### S3 Next Steps

While running the evaluation on the test set, it became evident that current best Helixer models have more issues with over-prediction (precision), while Helixer excelled at recall. Moreover, the over-prediction was not random, but concentrated in more fragmented genomes, and particularly relevant here over-prediction was occurring in large part on the small contigs below the subsequence length, i.e. those with padding.

We hypothesize that this is due to an early decision we made on how to handle small contigs without genes, which could either result from no-gene calling having been performed on the small contigs (not uncommon), or of course, no genes being present. At the time, we elected to avoid including likely erroneous data and filtered completely unannotated contigs from the training data, ignoring the bias this created in the remaining small contigs. This decision appears to have become problematic once the training species sets were expanded to include highly fragmented genomes. Thus, we have given the network a shortcut, which–like the cautionary example of a medical model making predictions based on a hospital-specific token instead of the lung image (https://doi.org/10.1371/journal.pmed.1002683)–lets the model achieve higher training accuracy by over predicting genes on contigs when it sees padding, at the expense of relying on biological features.

We report this while it is still under development for three reasons, 1) allow the earlier documentation of models that are, despite this issue, state-of-the-art, 2) inform users of current best models about likely error types, and 3) as a genomics-field example for potential Deep Learning shortcuts that modellers might face.

Both less naïve filters and data augmentation (shift and pad subsequences from well-assembled genomic regions) are being considered and tested to address this problem and the next generation of models should be available soon.
